# Metabolic resilience is encoded in genome plasticity

**DOI:** 10.1101/2021.06.25.449953

**Authors:** Leandro Z. Agudelo, Remy Tuyeras, Claudia Llinares, Alvaro Morcuende, Yongjin Park, Na Sun, Suvi Linna-Kuosmanen, Naeimeh Atabaki-Pasdar, Li-Lun Ho, Kyriakitsa Galani, Paul W. Franks, Burak Kutlu, Kevin Grove, Teresa Femenia, Manolis Kellis

## Abstract

Metabolism plays a central role in evolution, as resource conservation is a selective pressure for fitness and survival. Resource-driven adaptations offer a good model to study evolutionary innovation more broadly. It remains unknown how resource-driven optimization of genome function integrates chromatin architecture with transcriptional phase transitions. Here we show that tuning of genome architecture and heterotypic transcriptional condensates mediate resilience to nutrient limitation. Network genomic integration of phenotypic, structural, and functional relationships reveals that fat tissue promotes organismal adaptations through metabolic acceleration chromatin domains and heterotypic PGC1A condensates. We find evolutionary adaptations in several dimensions; low conservation of amino acid residues within protein disorder regions, nonrandom chromatin location of metabolic acceleration domains, condensate-chromatin stability through cis-regulatory anchoring and encoding of genome plasticity in radial chromatin organization. We show that environmental tuning of these adaptations leads to fasting endurance, through efficient nuclear compartmentalization of lipid metabolic regions, and, locally, human-specific burst kinetics of lipid cycling genes. This process reduces oxidative stress, and fatty-acid mediated cellular acidification, enabling endurance of condensate chromatin conformations. Comparative genomics of genetic and diet perturbations reveal mammalian convergence of phenotype and structural relationships, along with loss of transcriptional control by diet-induced obesity. Further, we find that radial transcriptional organization is encoded in functional divergence of metabolic disease variant-hubs, heterotypic condensate composition, and protein residues sensing metabolic variation. During fuel restriction, these features license the formation of large heterotypic condensates that buffer proton excess, and shift viscoelasticity for condensate endurance. This mechanism maintains physiological pH, reduces pH-resilient inflammatory gene programs, and enables genome plasticity through transcriptionally driven cell-specific chromatin contacts. In vivo manipulation of this circuit promotes fasting-like adaptations with heterotypic nuclear compartments, metabolic and cell-specific homeostasis. In sum, we uncover here a general principle by which transcription uses environmental fluctuations for genome function, and demonstrate how resource conservation optimizes transcriptional self-organization through robust feedback integrators, highlighting obesity as an inhibitor of genome plasticity relevant for many diseases.

## Introduction

Dynamic 3D genome organization plays important roles in guiding gene regulation, and cellular response to environmental stimuli. It also confers robustness to perturbations, and stochastic control to programmed differences between cells and between genotypes (*1*). Biophysical phase separation, on the other hand, enables both rapid formation of chromatin domains and transcriptional assemblies by phase transitions (*2–5*), in which transcriptional co-activators mediate multi-enhancer condensate-chromatin assemblies (*3, 6*). This multi-molecular cooperation has helped define hub-models of transcription, where genome activity drives cell-specific usage of chromatin architecture (*6*). The flexibility of genome organization is guided by biophysical principles including polymer-polymer interactions, diffusional motion, phase separation, and architectural constraints (*5*). Indeed, composition of transcriptional regulators along with architectural principles grant cell-specific function in response to environmental fluctuations (*3*). The ubiquitous conservation of genome structural features and the stochasticity of gene expression are hallmarks of self-organized systems. In this regard, cellular states can provide temporal stability of the system, while state transitions, influenced by environment oscillation, offer tunable feedback integration (*5*). Therefore, composition and regulation of heterotypic nuclear assemblies are important for feedback tuning of self-organization (*4*). This is seen during cellular adaptations, where compartmentalization of proteins by phase separation reduces noise from biological stochasticity (*7*). This mechanism, for example, ensures cell survival during phenotype transitions by reinforcing self-organization tuning-strategies, on top of an evolved and flexible framework (*5*). Then, transcriptional organization allows the cells to transition from different states without losing their functional identity (*1*) by coupling 3D dynamic transcriptional processes with environmental signals and with modulatory features of the chromatin-hardware (*2–5*).

Formation of self-organized assemblies synchronizes with metabolic fluctuations to better use thermodynamic states favouring phase transitions (*8, 9*). Coupling the degree of self-organization with favourable states such as cold and fasting has implications on their tuning, and more broadly on cellular and evolutionary adaptations (*8, 9*). Cold and fasting, through reduced cytosolic volume, promote cell crowding and less molecular diffusion, which increases molecular interactions and assembly of condensates (*9–13*). Since their formation is driven by entropic-forces and less by ATP (*9, 14, 15*), elucidating their range of parametrization is crucial to understand cellular adaptations (*8*). Indeed, cellular states condition the material properties of condensates, which impact the collective behavior of interactions and their viscoelasticity (*16*). For example, during energy depletion, RNA-demixing of P-granules promotes liquid-like to gel-like transitions for fitness (*8, 17*). A process optimized by evolution through the selection of protein residues favouring multi-molecular assemblies (*8, 18*). This is in line with resource-driven selection, where preservation of energy during nutritional limitation acts as a selective pressure for evolutionary adaptations (*19*). This selection is also encoded in genomic-rearrangements or cis-regulatory substitutions (*20*) (also known as human accelerated regions, HARs), and 3D optimization of chromatin interactions (*21*). It remains unknown, at small scales of cellular adaptation, how energetic states integrate tuning of transcriptional condensate transitions with genome architecture, and at larger scales, how it drives evolutionary innovation.

The nuclear architecture provides a structural framework that guides the concerted action of multi-dimensional players for genome function (*5*). Here, we use the genome architecture to supervise the integration of multi-layered information into transcriptional hub-models rendering transcriptional condensate processes. By using the probabilistic and combinatorial nature of genome activity, along with its architecture as topological coordinates, we show that integration of multi-omic information into genomic territorial hubs identifies functional dependencies between evolutionary, phenotypic, structural and transcriptional data. Our integrative topological approach reveals metabolic human acceleration domains with nonrandom higher-order location, transcriptional regulatory networks, and functional chromatin interactions mediating resilience to nutritional stress. This is regulated by heterotypic peroxisome-proliferator-activated receptor-γ coactivator-1α (PGC1A) condensates, and through the incorporation of environmental signals into genome plasticity. This mechanism optimizes transcriptional bursts of lipid-cycling genes, buffers proton accumulation and promotes endurance of cell-specific condensate-chromatin conformations. These findings uncover a general mechanism of environmental control of genome adaptations with far-reaching implications.

### Phenotypic and molecular adaptations reveal metabolic human acceleration domains

To discover human-specific adaptations in nuclear compartmentalization, including phenotypic, genetic, structural and transcriptional adaptations we used an integrative topological approach (**fig. S1 and S2**). Comparison of metabolic parameters among placenta-derived mammals has shown that primates optimized basal energy expenditure (*22*) (**Fig. 1A and S3A**). Great apes, including homo sapiens, displayed the lowest basal calorie consumption per day among mammals (**Fig. 1A and Table S1**), while humans showed the highest body fat percentage among great apes (*23*) suggesting a lineage-specific variation (**fig. S3B-D**). Specific energetic allocation and surplus of energy resources has played a crucial role in human evolution (*23, 24*). To understand the impact of fat mass on energy budget, we compared its effect on resilience to energy deficit as described before (*25*). Remarkably, compared to allometric fat mass, experimental fat mass showed species-specific acceleration to energy-deficit endurance (**Fig. 1A and Table S1**). These results indicate metabolic phenotypic differences among mammals with human lineage-specific acceleration of fat-associated endurance. To assess lineage-specific molecular signatures between humans and primates, we compared transcriptomic and epigenomic data from organs with high fat content such as brain and adipose tissue (*26, 27*), followed by functional annotation in tissue-specific networks (*28*). Consistent with phenotypic data, homo sapiens displayed enhanced transcriptional regulatory mechanisms and decreased lipid degradation (**Fig. 1C and Table S2**). To study the evolutionary and functional impact of non-coding regulatory elements, we interrogated molecular signatures from genes associated with regions with human-specific substitutions (*20, 29, 30*) (**fig. S3E and Table S3**). We found overlapping tissue-specific representations of transcriptional regulation and lipid metabolism for genes within 1Mb of HARs (**fig. S3F-H**). This suggests that phenotypic optimization of metabolism in human lineage has been parallel with molecular evolutionary adaptations in organs with high-fat content. Given the enrichment of transcriptional mechanisms, we investigated whether transcriptional regulators of HAR-genes display physical interactions, share biological processes or are HAR-associated (**fig. S4A**). ChIP-seq- and transcription factor binding sites (TFBS) enrichment analysis revealed promoters of HAR-genes are bound by physically interacting transcriptional regulators displaying common biological signatures (**fig. S4B-D and Table S4**). Targeted (1+ interactors) and untargeted (full) network reconstitution and prioritization of bound interactors showed overrepresentation of co-activators and nuclear receptors (**Fig 1D, fig. S4D-F and Table S5**). Remarkably, only CREB1 and PGC1A displayed HAR association (**Fig. 1D**), whereas annotation of these cooperative regulators revealed they control adipocyte differentiation and lipid metabolism (**fig. S4G**).

**Fig. 1.**
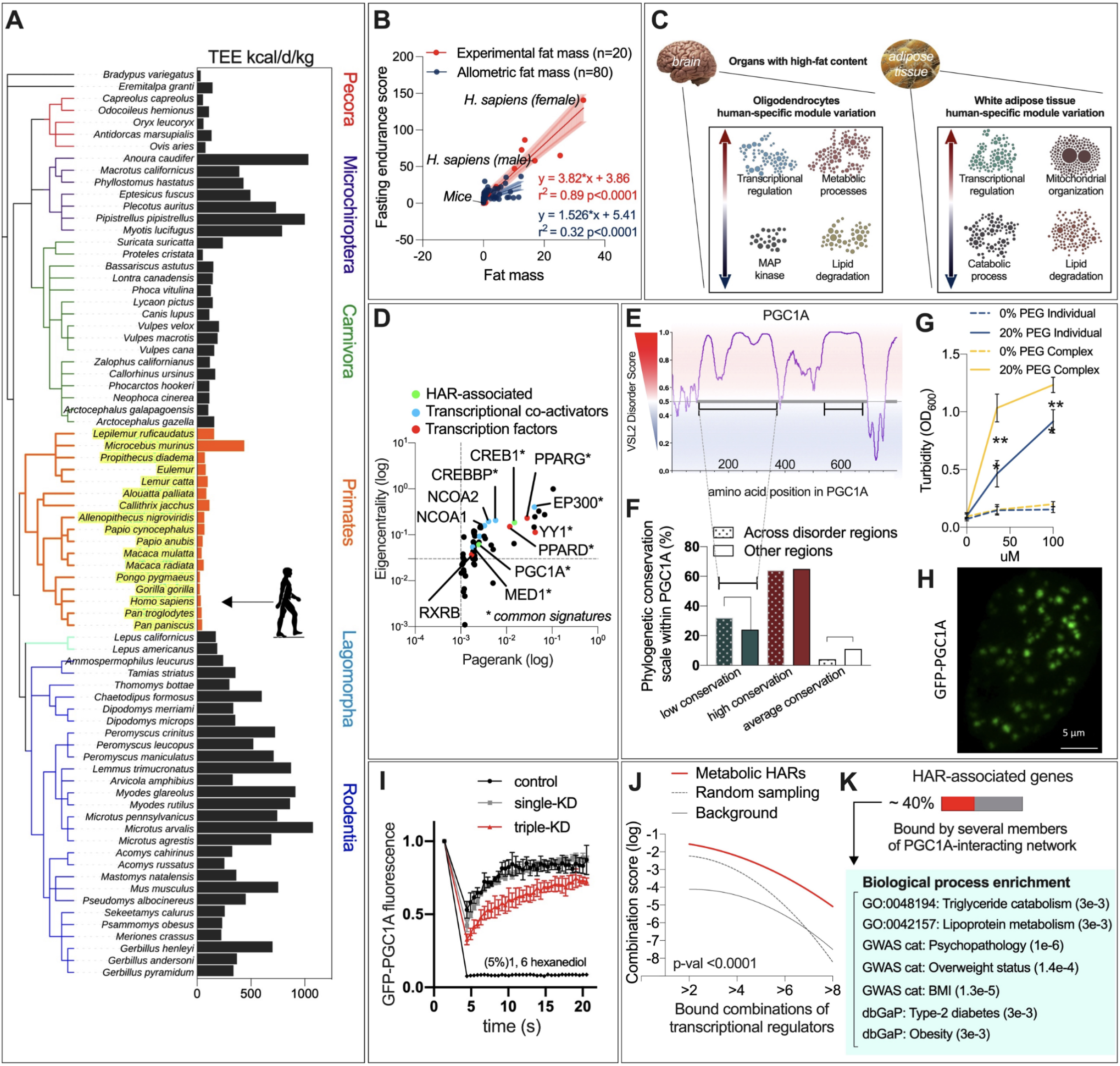
Evolutionary entanglement of metabolic phenotype and molecular adaptations. (**A**) Total energy expenditure across mammals (*22, 23*). (**B**) Comparison of energy deprivation score between allometrically scaled fat mass and experimentally measured fat mass across mammals. (**C**) Network-derived discovery of biological processes (*28*) in human enriched gene sets from tissues with high-fat content. Data derived from differential expression by RNAseq and ATACseq comparing human and primate tissues (*26, 27*). (**D**) Network-based node prioritization of recomposed interacting transcriptional regulators binding to HAR-associated genes. In green, regulators in proximity to HAR elements. (**E**) VSL2 disorder score for the transcriptional coactivator PGC1A amino acid sequence. (**F**) Proportion of amino acid conservation by phylogenetic analysis, within disorder regions and other protein domains in PGC1A. (**G**) Validation of phase separation by turbidity assays at different concentrations of PGC1A and RARA recombinant proteins with 0 or 20% polyethylene glycol, individually and in complex (n = 3 replicates). (**H**) Confocal microscopy image for GFP-PGC1A in living cells. (**I**) Fluorescence recovery and photobleaching analysis (FRAP) of GFP-PGC1A condensates in living cells treated with forskolin for transcriptional induction. Comparison of knockdown (KD) perturbations of PGC1A-interacting regulators, PPARG alone, and in combination PPARG, RXRA RARA, along with 1,6 hexanediol. (**J**) ChIP-seq integration, combinatorial regulation of HAR-associated genes by PGC1A-interacting network. (**K**) Biological enrichment of HAR-associated genes bound by a combination of several members of the PGC1A-interacting network. Data show mean values and error bars indicate SEM. Unpaired, two-tailed student’s t-test was used when two groups were compared, and ANOVA followed by fisher’s least significant difference (LSD) test for post hoc comparisons for multiple groups. * p-value < 0.05.

Combinatorial regulation of transcription includes condensates as “integrators” of heterogeneous protein organizations and genome architecture (*3*) (**fig. S5A**). Protein phase separation can be studied with the identification of low complexity intrinsically disordered regions (IDRs) (*31*). Given that PGC1A is a promiscuous interacting co-activator and a master regulator of metabolism (*32, 33*), we derived computationally, individual and cumulative phase separation scores for PGC1A-network. PGC1A is a highly disordered protein (>70% of aa sequence; **Fig. 1E, fig. S5B and Table S6**), that interacts with highly disordered co-regulators (**fig. S5C-D**). We estimated ongoing evolutionary adaptations within intrinsically disordered regions, by examining the phylogenetic conservation of PGC1A residues (**fig. S6A-B and Table S7**) (*34*). Remarkably, the proportion of residues with low conservation was higher within IDRs compared to other protein domains (**Fig 1F**). We used tissue-specific networks to assess PGC1A-network and its condensate compositional dependencies to adipose tissue and brain (*28*). More than 80% of its core members were predicted or have been validated to form phase separation (**fig. S6C**) (*35*). Further, physically interacting transcriptional regulators displayed more cumulative phase separation scores compared to random interacting permutations (Chi-square <0.001; **fig. S6D**). Turbidity assays of recombinant proteins confirmed phase separation of PGC1A and one of its interacting members RARA, individually and in complex (**Fig 1G and fig. S7A**). PGC1A Fluorescence recovery after photobleaching (FRAP) experiments in living cells activated with forskolin and perturbed with PGC1A-interactors revealed that heterotypic TF-co-activator complexes display increased stability and different dynamics than homogeneous assemblies (**Fig 1H-I and fig. S7A-B**). These results indicate evolutionary pressure within the IDRs of promiscuous co-activators, and that heterogeneous TF-co-activator complexes are more stable compositions. This prompted us to assess combinatorial transcriptional binding of top PGC1A interacting regulators. To this end, we used ChIP-seq data from the ChIP-atlas repository (*36*). Combinatorial analysis showed that PGC1A interacting regulators are co-binding the promoters (10Mb to TSS) of several HAR gene clusters (f**ig. S7C-D and Table S8**), with a large number of co-regulated genes showing co-activator dependencies (**fig. S7D-E**). This indicates physically interacting regulators with common biological signatures display specific binding density distribution to co-regulated genes (**Fig 1J**). This refinement step revealed that PGC1A-network co-regulates around 40% of genes contiguous to HARs, whose biological annotation is related to lipid metabolism (from now on metabolic-HAR domains; **Fig 1K**). Collectively, these results indicate there are evolutionary constraints affecting chromatin functionality as HARs are related to genes with similar function, and that transcriptional condensate hubs such as co-activators display low conservation of residues within IDRs.

### Metabolic-HARs exploit functional features of spatial genome architecture

Metabolism couples environmental variation to cytosolic signals controlling cellular adaptations (*37*). These functional adaptations entail the integration of genome architecture and nuclear condensates into transcriptional hubs (*5, 38*). To dissect evolutionary constraints on functional chromatin regions, we determined genome-wide structural relationships of metabolic HAR domains (**fig. S8A**). We first mapped metabolic HAR genes into topological domains using positional gene enrichments (PGEs) (*39*). Remarkably, biologically derived gene sets always recapitulated highly enriched genomic clusters as an indication of active transcriptional hubs (**fig. S8B-D and Table S9**). Functional annotation of PGEs revealed chromatin regions of similar function and related phenotype association (fPGEs) (**Fig 2A and Table S9**) To uncover genome-wide structural relationships for functional topological clusters, we interrogated their relational behavior in integrative networks for long- and short-range chromatin interactions (**Fig 2B**). Using a compendia of chromatin contacts (*40*), we found that metabolic-HAR domains are preferentially located in intrachromosomal and interchromosomal interactive regions as well as in regions with conserved topological associated domains (TADs) (**Fig 2C-D, fig. S8E-F and Table S10**). Network integration of human embryonic stem cells HiC data (*41, 42*) revealed functional metabolic-HAR domains (fPGEs) are located within hubs and compartment (cliques) forming regions. (**Fig 2E, fig. S8G and Table S11**). The location within hubs and domains improving resilience of the structural-network indicates that, throughout evolution, they have exploited topological features of genome architecture. We next investigated short-range constraints, through network integration of interactions among cis-regulatory regions (*43*) (**Fig 2F and Table S12-13**). We used module-based hierarchical models of short-range interaction frequencies (**Fig 2F and fig. S8H-I**), to assess the density of interactions, and categorized regulatory regions as social or isolated (**Fig 2G and fig. S8J-K**). Interestingly, metabolic HAR enhancers were preferentially linked to genes with social promoters (**Fig 2H**), while active enhancers displayed more elements with one interaction than genome-wide averages (**fig. S8L**). These results indicate evolutionary constraints on local genome architecture as enhancers with human-specific substitutions are especially linked to social promoters, suggesting fine-tuning of highly regulated hubs.

**Fig. 2.**
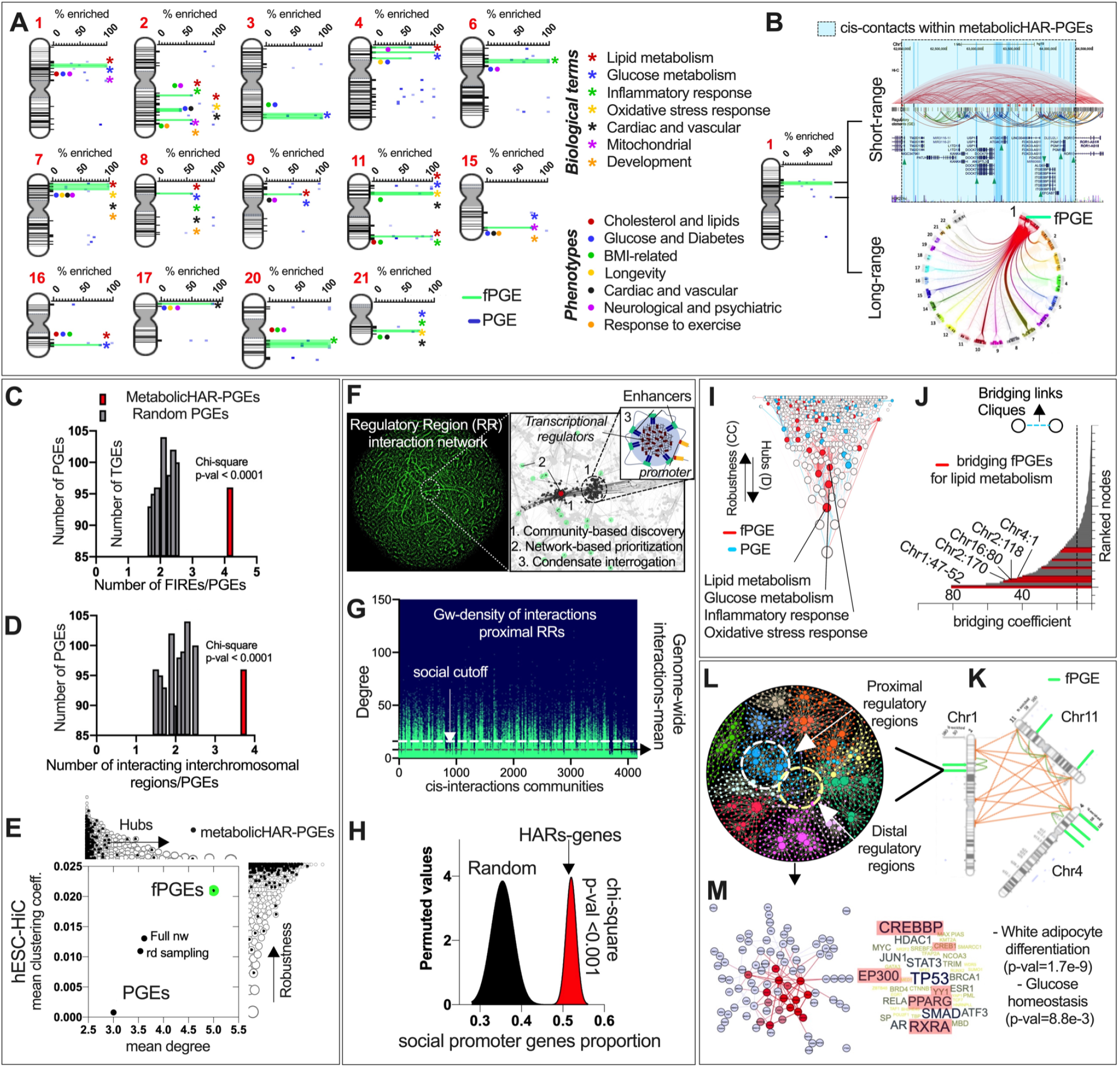
Metabolic-HAR domains exploit functional features of spatial genome architecture. (**A**) Metabolic-HAR genes positional gene enrichments (PGEs) and functional domain annotation (fPGEs). (**B**) fPGEs and PGEs domains in chromosome 1 with short- (chr1:62-65mb) and long-range structural relationships. (**C**) Overlapping frequently-interacting regions (FIREs) within metabolic-HAR regions. (**D**) Overlapping trans-interchromosomal regions from hESC-HiC data within metabolic-HAR regions. (**E**) Mean clustering coefficient (robustness) and node-degree (hubs) across nodes of trans-interacting loci-loci network from hESC-HiC interactome. Labelled, node-loci with colocalized fPGEs, PGEs, full network and random sampling. (**F**) Network-mediated interrogation of *cis*-genome-wide relationships among regulatory regions (*43*). (**G**) Node-degree of proximal regulatory regions across diverse network communities from F. (**H**) Genes associated with metabolic-HARs display highly social promoters (node-degree >16) compared to random gene sets. (**I**) Clustering coefficient (robustness) and degree (hubs) hierarchies from hESC-HiC network. Genes within top fPGE regions display metabolic function. (**J**) Bridging coefficient in hESC-HiC network. In red, top bridging chromatin regions harbouring metabolic-HAR genes with lipid metabolic function. (**K**) Hub-bridging mediated chromosome reconstruction of nuclear compartments for lipid metabolism displaying selected fPGEs. (**M**) Cis**-**regulatory network within selected fPGE-modules. (**N**) Module-mediated ChIP-seq binding enrichment within fPGE followed by PPI-network of regulators, and functional biological annotation.

Chromatin interactions cluster multiple genes with related functions into transcriptional hubs (*44*). We obtained functional nuclear compartments by assessing interactions between metabolic HAR regions of similar biological function (**Fig. S9A-B and Table S11**). Indeed, regions exhibiting high node degree and high clustering coefficients harbour HAR associated genes with lipid, glucose, inflammation and oxidative stress function (**Fig 2I**). In graph theory and network evolution, modulation of network information flow is better done through nodes bridging densely connected modules (*45–47*), rather than highly-connected nodes, as hub modulation can be costly for network integrity (*47–49*). Therefore, we investigated critical bridging HAR regions with similar biological function to hub structors. This showed several regions around chr1:48.5mb housing metabolic HAR genes with lipid and glucose function (**Fig 2J, fig. S9B and Table S11**). In agreement, chromosome interactions around other top functional connectors were pervasive in other HiC datasets (**fig. S9C-D**). To isolate critical regions that exploit information flow, and influence global genome architecture, we recomposed functional nuclear compartments enriched in top bridging nodes, guided by 3D homotypic hESC-HiC reconstruction (**Fig 2K and fig. S10**). Functional ranking showed 2 subnetworks built around top bridging nodes with fPGEs for lipid metabolism (**fig. S11A**). Collectively, these indicate that evolution optimizes metabolic functional hubs by intervening chromatin domains with potential systemic effects.

We found that, among all metabolic-HAR regions, approximately 84% harbour experimentally validated non-coding elements (*50*) (**fig. S11B**). Within the reconstructed nuclear compartment, every metabolic-HAR region showed active elements locally (**fig. S11C**). We next assessed topological hierarchies and local chromatin states, by network modular analysis of regulatory interactions (**Fig 2L, fig. S12A-B and Table S14-15**). Node prioritization showed that interaction frequencies directly correlate with chromatin states, as isolated enhancers are enriched in repressive epigenetic tags such as H3K9me and H3K27me (**Fig 2M and fig. S12C-D**). Consequently, we used ChIP-seq enrichment of transcriptional regulators bound to both social enhancers and promoters to assess regulators within the topological domains (**fig. S12E-F**). We then filtered binding regulators that physically interact, followed by network functional enrichment (**Fig 2M and fig. S12G**). This revealed that local topological territories can be classified by function as an indication of regional transcriptional activity (**Fig 2M and Table S14**). Subregions with metabolic HARs were bound by interacting regulators controlling lipid and glucose metabolism, with several members of the PGC1A-network (**Fig 2M and Table S14**). Interestingly, every submodule showed local interaction decay with distance constraints for submodular communication (**fig. S12H**). To assess if modular-territories in the regulatory network are representations of structure-based domains such as TADs and chromatin contact domains (CCDs), we investigated global and local CTCF-mediated 3D reconstruction (**fig. S13A-B**) (*51, 52*). Locally, chromatin interaction analysis by paired-end tag sequencing (ChIA-PET) and CTCF-mediated reconstruction from several cell lines (*51*) revealed a clear overlap between CTCF-cluster graphs and network submodule territories (**fig. S13C**), along with intra and inter-domain (e.g. TAD) 3D effects of structural variants on the metabolic-HAR region (**fig. S13D-E**) The overlap between structural regions and regulatory frequency networks suggests the existence of intrinsic properties of genome architecture.

Functional enrichment of chromatin territories and interacting regions produced complex functional hierarchies (**Fig 2K**). This suggests that structural dependencies might confer genome robustness as novel substitutions can be tolerated while being tested. Likewise, substitutions that lead to DNA misplacement affect evolutionary adaptations and disease (*53, 54*). We thus assessed territorial functional and phenotype association, including structural variants (*55*) and genomic-range GWAS (*56*) (**fig. S14A-D and Table S16**). This revealed interacting hierarchical maps of trait enrichments including lipid, glycemic and BMI in metabolic-HAR regions of selected nuclear compartments (**fig. S14D-F**). To obtain genomic-range trait enrichments with deeper resolution, including tissue-specific enhancer modules with trait enrichments and dependencies, we used data from EpiMap (epigenomic integration across multiple annotation projects) (*57*). Hierarchical network analysis yielded the polyfactorial penetrance of metabolic and lipid-related traits, and locally with traits in our reconstructed nuclear compartments (**fig. S15A-B and Table S17**). For those traits within the same modules, we built hub models of topologically related traits and tissues (**fig. S15A-C**). This showed tissues with lipid functions are drivers of disease and multifactorial lipid related traits are bridging across other traits with hierarchies following structural functionality (**fig. S15B-D**).

### Metabolic-HAR activity and heterotypic nuclear compartments control fasting endurance

During cellular adaptations, phase transitions entrain the integration of environmentally mediated signals into self-organized conformations (*5, 58*). Environmental signals such as solvation and salt concentration, for example, affect self-organization through intrachain electrostatic repulsion-attraction (*59*). We thus used CIDER to dissect context-dependent condensate conformations from amino acid sequences (*59*). Among members of the PGC1A-network, YY1, PPARG and PGC1A displayed context-dependent globule conformations (**Fig 3A**) with well distributed charged residues and increased proportion of basic residues in PGC1A IDRs (**fig. S16A-B**). Given that PGC1A coactivators and interacting regulators control energy homeostasis (**fig. S5C**) and that species-specific variations of metabolic parameters are related to energy endurance and fat tissue (**Fig 1**), we interrogated nuclear compartmentalization during fasting in human adipocytes. First, we observed that glucose fasting promotes cytosolic pH acidification (**Fig 3B**). This is in line with protonation-mediated phase separation of PGC1A, individually and in complex (**Fig 3C and fig. S16D**), along with increased expression of PGC1A and its interacting members in fasting (**fig. S16E**). Transcriptional condensates control chromatin interactions by limiting stochastic motion and increasing persistent interactions (*60*). For this reason, we used structuring proteins identified by ChIP-MS bound to both enhancers and promoters (*61, 62*). In PPI networks, we found that PGC1A-network associates with structuring candidates CTCF, and YY1, both mediating enhancer-promoter loops (*62*) (**Fig 3D and Table S18**). Given that genomic clusters formed by binders within heterogeneous condensates might be larger than isolated chromatin binders (*6, 63–65*) (**Fig 3D and fig. S16F**), we tested PGC1A influence on hub models of nuclear compartmentalization. This revealed that fasting promotes CTCF expression and CTCF-mediated interactions around metabolic-HAR of selected nuclear compartments (**Fig 3E and fig. S16G**). In the same conditions, PGC1A bound to ncHARs within chromatin regions of interest (**fig. S16H**), while HAR-associated genes showed increased regional transcriptional activity for top mediators such as ATG4C, PGC1A, INSIG2, EIF4EBP1 and CAT (**fig. S16H-I**). Accordingly, fasting increased YY1-mediated ncHAR enhancer-promoter loops (**Fig 3F**).

**Fig. 3.**
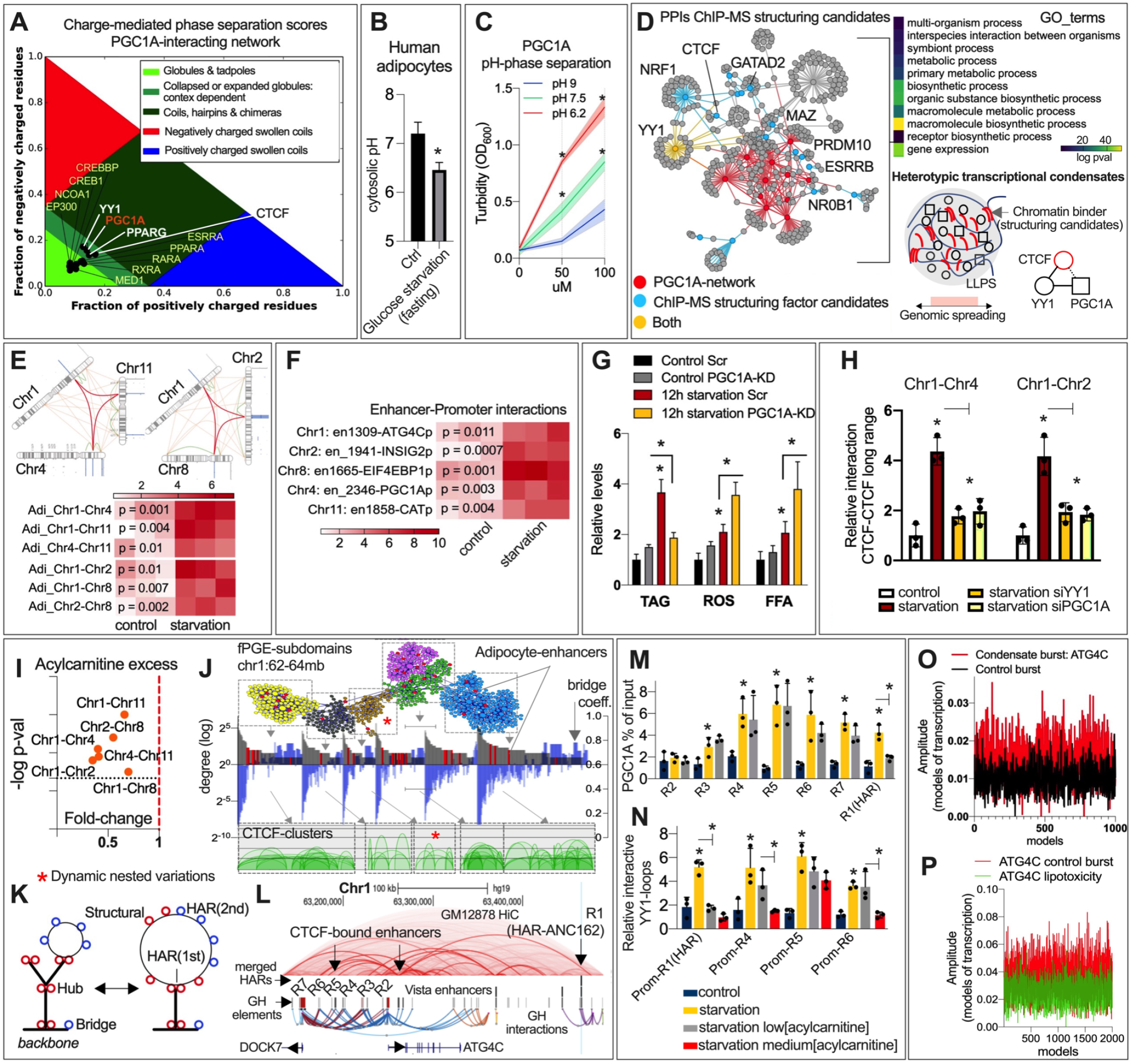
Metabolic-HAR nuclear compartmentalization controls fasting endurance. (**A**) Context-dependent phase separation score by CIDER (*59*) of PGC1A-interacting regulators, and CTCF chromatin binder. (**B**) Cytosolic pH variation after 12h of glucose starvation in human adipocytes. (**C**) Turbidity assays of recombinant PGC1A phase separation mediated by pH variations. (**D**) Transcriptional regulator protein-protein interactions for PGC1A-network and structuring transcriptional candidates selected by ChIP-MS (*61*), their functional annotation and illustration of cooperative heterotypic regulation of condensates. (**E**) CTCF-mediated ChIP-loop of fPGE-fPGE interaction within selected nuclear compartments as in B. (**F**) YY1-mediated enhancer-promoter loops between HAR-elements and starvation-regulated genes within fPGEs as in B. (**G**) Triglycerides, reactive oxygen species, and free fatty acid functional assays in human adipocytes after starvation and with siRNA for PGC1A. (**H**) CTCF-mediated long-range chromatin interactions by ChIP-loop after 6 hours of starvation and with targeted siRNA. (**I**) CTCF-mediated long-range ChIP-loop interactions in human adipocytes after starvation with acylcarnitine (AC) excess (20uM). (**J**) Network-based model of regulatory region interactions (top panel) and submodules overlap with CTCF-mediated chromatin contact domains (CCDs) clusters (top down panel). Chr1-fPGE submodules with adipocyte-specific enhancers in red and ATG4C submodule with an asterix. Hub degree and bridge divergent specialization of regulatory regions (middle panel). (**K**) Tree nested model representation showing dynamic chromatin variations with hierarchies of structural (CTCF-bound) and non-structural regulatory regions. (**L**) Human genome track showing ncHARs, regulatory elements and interactions, GM12878-Hi-C data, promoter locations, and structural and non-structural enhancers (R1 to R7). (**M**) ChIP-qPCR binding of PGC1A to regulatory elements in human adipocytes during starvation + low concentration of acylcarnitines (1uM). (**N**) ChIP-loop enhancer-promoter interactions in the same conditions as in **M** (medium AC = 10uM). (**O**) Amplitude of Monte Carlo models of transcription burst built from dynamic expression data for ATG4C and control gene in the same conditions as in **M**. (**P**) Amplitude of transcriptional burst models as in **O** but with high concentration of acylcarnitine (20uM). Bars show mean values and error bars indicate SEM. Unpaired, two-tailed student’s t-test was used when two groups were compared, and ANOVA followed by fisher’s least significant difference (LSD) test for post hoc comparisons for multiple groups. * p-value < 0.05.

ATG4C promotes fast generation of fatty acids through lipophagy (*66*). DGAT1, on the other hand, mediates fatty acid re-esterification as a protective mechanism to lipotoxicity (*67, 68*). Interestingly, DGAT1 loci is located at the tail of chromosome 8, a highly interacting region within the nuclear compartment for lipid metabolism (**fig. S16K**). Accordingly, fasting increased DGAT1 expression, PGC1A-binding to DGAT promoter and CTCF-mediated interactions between metabolic-HAR fPGE and DGAT region (**fig. S16K**). Reflecting DGAT1 enzyme activity, functional assays showed elevated esterification of fatty acids (**Fig 3G**). In agreement, knockdown of PGC1A blunted this effect while leading to excessive levels of oxidative stress (**Fig 3G**). To assess if other metabolic-HAR genes within the compartments control lipotoxicity, and whether this response is dependent on mitochondrial CPT1A, we performed functional assays after gene perturbations and after CPT1A-inhibition. Perturbation of genes within the compartments led to similar phenotypes (fatty acid accumulation and oxidative stress), which were exacerbated by CPT1A-inhibition (**fig. S16L-M**). To determine if compartment formation is dependent on heterogeneous condensates enriched in PGC1A, we investigated the time after fasting when compartment formation precedes phenotypic malfunction. We found that, although PGC1A gene perturbation has no effect on functional assays after 6h of fasting (**fig. S16N**), it blunts both long- and short-range chromatin interactions (**Fig 3H and fig. S16O**). Nuclear compartment formation is dependent on liquid-like condensates, as 1,6 hexanediol blocked chromatin interactions (**fig. S16P**). Similar results were obtained by blocking glycolysis, which led to increased ROS, FFA, and more acidic pH (**fig. S17A-B**).

Incomplete oxidation of fatty acids leads to accumulation of acylcarnitines (*67*), which are associated with obesity and insulin resistance (*69*). Indeed, acylcarnitine excess inhibited fasting-mediated long- and short-range chromatin interactions (**Fig 3I and fig. S17C**). This effect was dependent on uncontrolled cytosolic acidification as activators of proton pump extruders improved chromatin interactions (**fig. S17D-E**). To investigate the topological penetrance of acylcarnitine and pH variations on heterogeneous looping of regulatory regions, we assessed whether structural hierarchies within TADs or subdomains contribute to genomic and condensate stability. To this end, we focused on the ATG4C-fPGE domain (**Fig 3J**). Besides the convergence between CTCF-cluster graphs and network submodules (**Fig 3J**), our integration revealed some pervasive architectural principles such as interaction frequency specialization of regulatory regions (**Fig 3J**). Decay on interaction probabilities was paralleled by an increment on bridging score for nodes within submodule boundaries (**Fig 3J and Table S14**). Since this is in line with polymer interaction models where nested-tree structures represent intra TAD variability (*70*) (**Fig 3K**), we evaluated if regulatory regions bound by structural proteins condition resilience to metabolic stressors. Interestingly, a large proportion of genes controlled by CTCF are promiscuous or social promoters, supporting a role for structuring candidates as anchors of transcriptional hub stability (**fig. S17F**). We first evaluated, during starvation, cell-type specific enhancers (*57, 71*) (**Fig 3J and Table 19-20**). Adipocyte-specific enhancers within the fPGE displayed higher levels of H3K27Ac epigenetic modification, despite not being bound by PGC1A during fasting (**fig. S17G-H**). This indicates that transcriptional activity is fine-tuned at different levels despite domain stochasticity and cell-type variation. For example, ATG4C structural enhancers bound by PGC1A exhibited different sensitivity to acylcarnitine excess (**Fig 3L-M**). HAR-element linked to ATG4Cp was highly sensitive to low levels of acylcarnitine, which might indicate an exposed 3D conformation during starvation (**Fig 3M**). On the other hand, loops formed on structural enhancers were resilient to elevated levels of acylcarnitine (**Fig 3M**), and, similar to local cell-specific enhancers, were bound by histone acetyltransferase EP300 and CTCF (**fig. S17I-J**). Perturbation of EP300 and CTCF during starvation led to loss of PGC1A-binding to HAR enhancer, and increased sensitivity to fatty acids as structural loops lost their resilience along with target gene expression (**fig. S17K-M**). Similar to previous reports (*72*), these results indicate structural hubs limit stochastic chromatin motion while following hierarchical principles of organization during genome function.

Since multi-enhancer hubs regulate the kinetics of transcriptional bursts (*73, 74*), we used a probabilistic approach to model burst profiles from dynamic gene expression. This showed that dynamic fluctuations of gene expression can be captured by nonlinear sine-wave models and Monte Carlo simulations. With this, we discriminated, in similar conditions, transcriptional burst kinetics for genes within the same TAD (**fig. S175N**). Compared to genes with isolated promoters such as ANGPTL3, dynamic fluctuations of HAR-associated genes with social promoters such as ATG4C, display higher amplitude and more frequent burst kinetics (**Fig 3O and fig S175O-Q**). Accordingly, these were susceptible and blunted by metabolic overload through acylcarnitine excess (**Fig. 3P**). Finally, we observed similar dynamic nuclear conformation during fasting in human neurons (fi**g. S18A-G**). Transcriptional profiles from human adipocyte precursor cells (APC) (*75*) and snRNAseq from human cortex (*76*), showed that high lipid turnover APCs and oligodendrocytes display elevated territorial expression of HAR-fPGEs genes (**fig. S18H-I and Table S21**). Collectively, these results underscore an ongoing evolutionary pressure for burst stability, as ncHARs are linked to social promoters with specific burst kinetics. At the molecular level, these topological constraints allow the identification of functional feedback transcriptional hubs controlling metabolic endurance. (**fig S17Q**) (*61, 62*).

### Metabolic-HAR domains shows species-specific convergence and tuning

To assess principles of transcriptional hub activity across mammals, we first determined syntenic conservation of metabolic HAR domains in mice. This revealed a high degree of chromosomal synteny (**fig S19A and Table S22**), with some domains being split in murine chromosomes (**Fig. 4A-B**). Interestingly, metabolic HAR regions displayed higher conservation than chromosomal TAD boundaries (breakpoints; around 54% of human boundaries shared with mice) (*41*), and less than syntenic blocks (**Fig.4B**). Regarding the transcriptional networks mediating fasting endurance in humans, regions in chromosome 2 and 4 were split in murine chromosomes. These results indicate a high degree of preservation of metabolic-HAR blocks but also convergent rearrangements in humans for distant HAR-regions in mice (hChr2, 4, 7 and 18). Since compartmentalized hubs coordinate territorial transcriptional activity (*3*) (**Fig 3**), we interrogated whether phenotypes from mammalian perturbation data represent territorial activity with congruent phenotypes. Using mice knockout data displaying phenotype and gene relationships followed by network integration (**fig S19B and Table S22**), we found that metabolic HAR genes form hierarchical modules of overlapping phenotypes (**Fig 4B and S19B-C**). Accordingly, metabolic-HAR genes mediating metabolic endurance are located within metabolic, immune and CNS modules (**Fig 4B and fig. S19D**). Unbiased node prioritization confirmed several metabolic phenotypes among the most influential (**fig. S19E-F**). On the other hand, top drivers of metabolic, immune and CNS phenotype clusters were PGC1A, NFE2L2, and EN1 respectively (**Fig 4C and fig. S19E**), whereas module interaction analysis showed their topological dependency (**Fig 4D**). We thus assessed whether this phenotype relationship translates into specific genomic clusters. Remarkably, compared to random combinations or individual enrichments, combined positional enrichment of gene drivers for these related phenotypes showed increased topological coalescence with an increased number of regions harbouring > 5 genes (**Fig 4E and fig. S19G**). These highlight the heterotypic nature of transcriptional hub models where stochasticity, chromatin interactions, and condensate composition translate into territorial genome function.

**Fig. 4.**
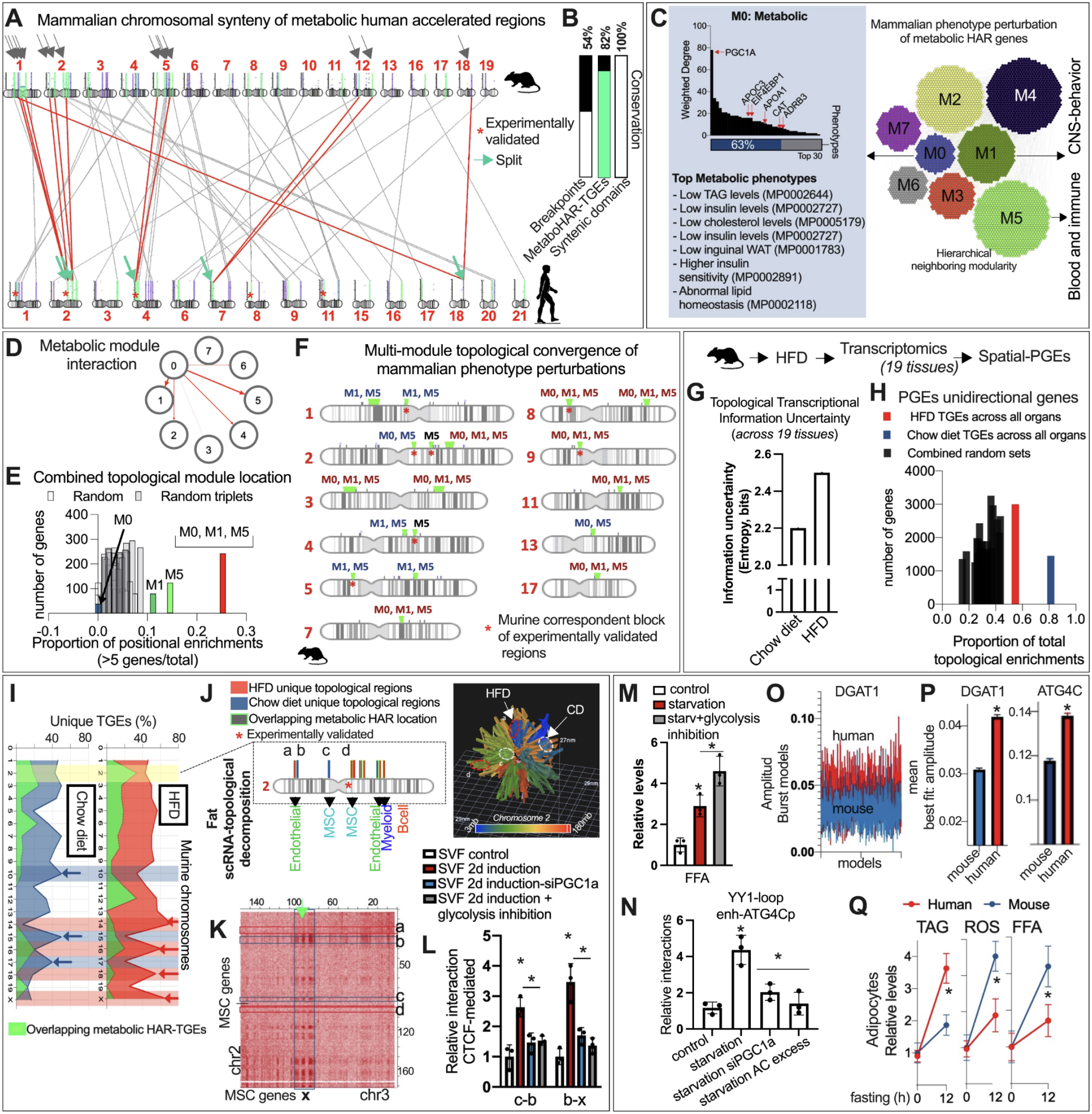
Species-specific convergence and tuning of nuclear compartmentalization. (**A**) Chromosomal synteny of human metabolic-HAR domains in mice. In red lines, split chromosomal regions. (**B**) Rate of mice-human conservation across different chromosome domains. (**C**) Network integration of mammalian genetic perturbations for metabolic-HAR genes. Hierarchical modularity (right panel) and node degree of gene knockouts within modules (left panel). (**D**) Metabolic inter-module interactions. (**E**) Proportion of PGEs (> 5 genes in genomic cluster) for genes in individual modules and metabolic interacting multi-module PGEs. (**F**) Multi-module topological convergence of metabolic interacting modules in murine chromosome regions. (**G**) Mammalian HFD perturbation followed by RNAseq in 19 tissues and spatial-PGEs analysis. Shannon entropy for number of PGEs (>5 genes) and total number of genes with genomic-cluster enrichment (PGE) across all organs. (**H**) Proportion of total PGEs for genes regulated by either HFD or chow-diet. (**I**) Murine chromosomal proportion of unique PGE hubs by HFD or chow-diet and overlapping syntenic metabolic-HARs. (**J**) Chromosome 2 location of HFD and chow-diet unique hubs (left panel) with 3D reconstruction and positional location of discrepant hotspots (right panel). scRNA-topological decomposition in fat using tabula muris dataset (*78*). (**K**) mESC-HiC chr2-chr3 contact frequencies showing hubs for mesenchymal stem cells from fat (fat msc) scRNA-decomposition. Light green arrow shows the location of metabolic-HAR regions. (**L**) CTCF-mediated compartments for fat-MSCs PGEs shown in **K** from induced stromal vascular cell fraction from subcutaneous adipose tissue. (**M**) Free Fatty Acid levels in differentiated murine adipocytes. (**N**) ChIP-loop YY1-mediated enhancer-promoter interactions for ATG4C gene in murine adipocytes. (**O**) Amplitude variation of transcriptional burst Monte Carlo models from dynamic gene expression data for murine and human DGAT1 expression. (**P**) Mean amplitude of burst models for DGAT1 and ATG4C. (**Q**) Murine and human divergent response to starvation by functional assays in adipocytes. Bars show mean values and error bars indicate SEM. Unpaired, two-tailed student’s t-test was used when two groups were compared, and ANOVA followed by fisher’s least significant difference (LSD) test for post hoc comparisons for multiple groups. * p-value < 0.05.

To understand the influence of diet on localized transcriptional gene-programs, we performed transcriptomic profiles in 19 tissues after diet-induced obesity. Differentially expressed genes per tissue were then interrogated for spatial positional enrichments (**fig. S19H and Table S23**). Comparison of tissue-specific and global PGEs demonstrated a territorial transcriptional coalescence as the proportion of chromatin regions with > 5 genes increased drastically in combined enrichments for HFD and chow diet (**fig. S19I and Table S24**). These analyses also revealed that HFD exhibits higher topological enrichments for regions with 2-genes and isolated gene positions (**fig. S19H, and Table S24**). To compare the HFD impact on transcriptional stochasticity and uncertainty, we derived Shannon entropy using tissue-specific ratios between PGEs harbouring > 5 genes and the total number of regulated genes. HFD promoted a more pronounced transcriptional uncertainty observed in random positional enrichments (**Fig 4G**). By filtering gene-programs specific to HFD or chow diet (**fig. S19J**), and combined positional enrichments, HFD-specific genes were more randomly located, while chow diet genes formed transcriptional hubs (**Fig 4H**). A chromosomal map with unique highly enriched topological regions per phenotype revealed diet constraints on overlapping metabolic-HAR domains and a larger number of unique positions in small chromosomes (**Fig 4I and fig. S19K-L**). We found more divergence of unique regions in large chromosomes (**fig. S19K**), along with experimentally validated metabolic HARs mediating endurance (**fig. S19K**). To evaluate tissue-specific discrepancies in transcriptional hubs, we focused on gene-programs in adipose tissue subtypes. We found overlap of top positional enrichments across adipose tissues and tissue-specificity for less enriched domains (**fig. S20A and Table S25**). Furthermore, HFD constrained adipose transcription as there was a global shift on hub regions, with an increased proportion of inefficient PGEs harbouring only 2 genes (**fig. S20A**).

### Species- and cell-specific tuning of fasting endurance

Despite the heterogeneity of local genome architecture during transcription in neighboring single cells (*1, 77*), single cell transcriptional maps have revealed a degree of homogeneity. Given the relationship between global genome architecture and genome function, we interrogated if transcriptional regions can be decomposed into cell-specific hubs using scRNAseq data (*78*) (**fig. S20B and Table S26**). Compared to random permutations, chow-diet hubs overlap with cell-specific positional hubs derived from fat scRNAseq data (**fig. S20C**). In agreement with regional hub models, this approach confirmed cell-specific and shared clusters of topological activity: around 70% of chow-diet domains and 51% of HFD regions intersected cell-specific clusters (random overlap is 20-35%; **fig. S20D and Table S26**). Interestingly, mesenchymal stem cells and endothelial cells displayed the highest topological convergence with HFD-mediated hubs (**fig. S20D**). This result confirmed the pervasive nature of diet-excess on random transcription overload, at the tissue- and cell-specific level (**Fig 4G**). To validate diet loss of cell-specific hubs harbouring syntenic metabolic-HARs, we focused on the topological behavior of chromosome 2 in fat tissue (**Fig 4I**). Indeed, diet-dependent topological regions showed 3D discrepancies despite being neighboring, and cell-enriched (**Fig 4J and fig. S21A**). For example, closely located cell-specific domains for endothelial and mesenchymal stem cells displayed diet-mediated shifts (**Fig 4J and fig. S21A**). Using HiC data, we found that these neighboring regions form homotypic interactions with distant domains harbouring homogeneous cell-specific hubs and metabolic-HARs (**fig. S21B**). Remarkably, HFD promoted a cell-specific shift in transcriptional activity from heterotypic interacting regions to neighboring isolated domains (**fig. S21C**). For example, interacting interchromosomal regions (chr3-chr2) with cell-specific hubs (MSCs) showed transcriptional activity in chow-diet, which was lost after HFD (**Fig 4K and fig. S21C**). Accordingly, heterotypic and homotypic interactions increased in adipose stromal vascular fraction after induction of differentiation (**Fig 4L**). This effect was blunted by perturbation of PGC1A and inhibition of glycolysis (**Fig 4L**). Underscoring its fatty acid dependency, PGC1A knockdown caused further accumulation of fatty acids, which was exacerbated after glycolysis inhibition (**fig. S21D**). Similar functional results were observed in differentiated murine adipocytes (**Fig 4M**). Interestingly, formation of syntenic nuclear compartments mediating metabolic endurance in human cells after fasting was incomplete in split domains in mice (**fig. S21E**). For regulated loop interactions, condensate hubs and chromatin interactions were abolished by PGC1A knockdown and acylcarnitine excess (**Fig 4N**). To elucidate species-specific burst kinetics, we focused on local top regulatory regions harboring ATG4C, DGAT1, and PGC1A, which showed PGC1A-binding to linked HARs was absent in murine orthologue regions after fasting (f**ig S21F**). Burst kinetics showed that frequency and amplitude are elevated in humans compared to murine adipocytes (**Fig 4O-P and fig. S21G-H**). This human-specific tuning translated into more fasting-endurance with increased fatty acid esterification, less oxidative stress and less acidic cytosolic pH (**Fig 4Q and fig. S21I**). Overall, these results imply evolutionary optimization of burst kinetics during nutrient limitation. Indeed, tuning of species-specific regulatory enhancers in the form of anchors seems to increase the stability of self-organized mechanisms that improve the functional control of cellular endurance (**fig. S21J**).

### Metabolic-HAR domains share metabolic disease-variants hubs

Obesity and type 2 diabetes are multifactorial diseases caused by the cumulative effect of lifestyle and small genetic variants dispersed throughout the genome. To understand structure-function relationships of polygenic metabolic traits, we interrogated the topological hub-contribution of causal genes mediating the effects of genetic variations in non-coding regulatory regions (**fig. S22A**). To this end, we used our approach (**fig. S2B**) on data derived from novel causal inference methods that integrate GWAS traits with tissue-specific cis-eQTLs to build trait-, tissue- and gene module-specific maps for metabolic disease polygenic dependencies (*79*) (**fig. S22A-B and Table S27**). We focused on commonalities and specificities between BMI and T2D polygenic and mediation maps among diverse tissues (**fig. S22B-C**). This showed multi-tissue contribution to polygenic scores, with 44% of gene modules coming from tissues with metabolic function (**fig. S22C-D**). Among those, we found trait- and tissue-specific contributions in visceral fat, skeletal muscle, stomach, cerebellum and cortex (**fig. S22D-E**). Given our observations on territorial transcriptional activity in diet and genetic perturbations (**Fig 4**), as well as genomic-range genetics (**fig. S14-S15**), we determined global and tissue-specific territorial hubs for metabolic disease (**fig. S22F**) In agreement, we found clusters of topological coalescence as tissue-specific genes further enriched optimal PGEs with > 5 genes (**fig. S22G**). Interestingly, we observed trait-specific discrepancies as global BMI-hubs from combined multi-tissue genes converge more into high-risk topological hubs (70% for T2D and > 95% for BMI; **fig. S22G-H and Table S27**). Given the global territorial difference between BMI and T2D polygenicity and their risk-disease causal association, we interrogated local chromosomal divergence of merged-PGEs (**fig. S23A**). This revealed unique and shared positional hubs as well as colocalization dependencies with metabolic HAR PGEs (**Fig 5A and fig. S23A-B**). BMI-PGEs and metabolic-HAR regions mediating stress-resilience were enriched in larger chromosomes while T2D-PGEs and shared-PGEs in small chromosomes (**Fig 5A and fig. S23B**). Chromosome size and trait-specific territorial hubs suggest structure-function principles of radial chromatin organization as BMI-T2D divergence could potentially explain risk factor-to-disease functional topological transitions.

**Fig. 5.**
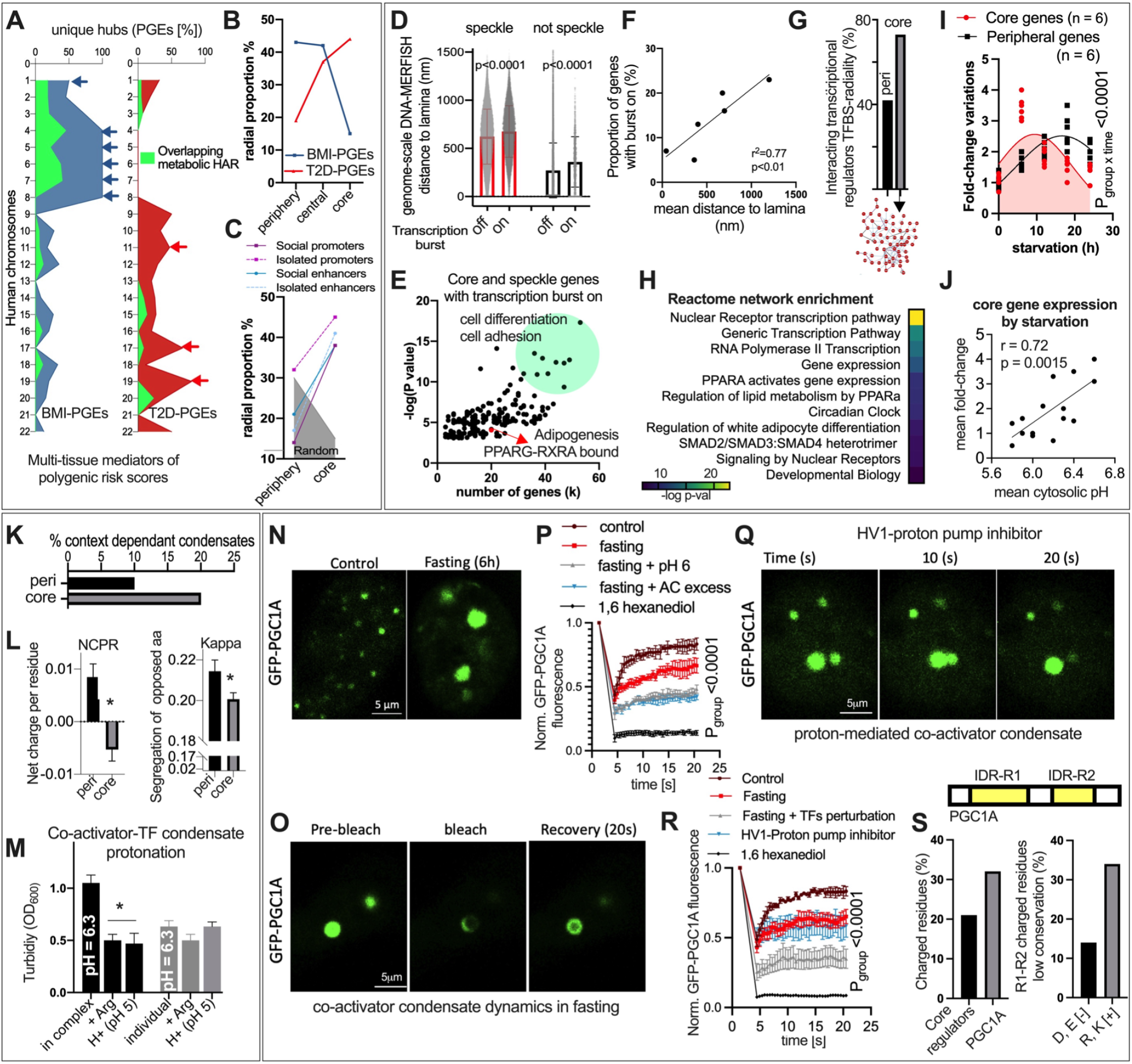
Chromatin radial divergence and gel-like condensate transitions control fasting endurance. (**A**) Chromosomal proportion of unique disease hubs from multi-tissue mediator genes of polygenic risk for BMI and T2D disease variants. (**B**) Radial proportion of disease variants hubs for BMI and T2D using GPseq-radial loci data (*80*). (**C**) Radial proportion enrichment for network-based categories of regulatory regions interaction density. (**D**) Distance to lamina of speckle associated genes with transcriptional burst on/off from genome-scale multiplexed FISH (*77*). (**E**) Functional annotation of core and speckle associated genes with transcriptional burst on from multiplexed FISH data (*77*). (**F**) Correlation between the proportion of genes (not associated with nuclear lamina) with transcriptional burst on and their distance to nuclear lamina from multiplexed DNA-FISH (*77*). (**G**) Proportion of protein-protein interactions among transcriptional regulators with TFBS radial enrichment (*80*). (**H**) Network reactome enrichment for core interacting transcriptional regulators. (**I**) Fold-change variation for genes with specific radial enrichment during starvation at different time scales in human adipocytes. (**J**) Correlation between mean fold-change variation and mean cytosolic pH during starvation in human adipocytes. (**K**) Proportion of transcriptional regulators (with TFBS radial enrichment) with amino acid sequences showing context dependent phase separation scores using CIDER (*59*). (**L**) Net charge per residue (left panel) and segregation of oppositely charged residues (right panel) in amino acid sequences from core and peripheral regulators using CIDER (*59*). (**M**) Phase separation by turbidity assay of PGC1A and RARA recombinant proteins in complex, individually, and with arginine and acidic-pH excess. (**N**) GFP-PGC1A confocal microscopy in living cells. (**O**) Confocal microscopy of FRAP-experiments in the same conditions as in N. (**P**) FRAP time of recovery and PGC1A condensate dynamics in living cells in control and starvation states. (**Q**) GFP-PGC1A confocal microscopy in living cells after HV1-proton pump inhibition. (**R**) FRAP time of recovery and PGC1A condensate dynamics in living cells in control, starvation, target perturbations and HV1-inhibitor. (**S**) Top panel shows PGC1A protein and consensus disorder regions. Proportion of charged residues and degree of conservation for positively and negatively charged residues in PGC1A. Bars show mean values and error bars indicate SEM. Unpaired, two-tailed student’s t-test was used when two groups were compared, and ANOVA followed by fisher’s least significant difference (LSD) test for post hoc comparisons for multiple groups. * p-value < 0.05.

### Radial genome organization pairs structure and function in metabolic adaptations

To evaluate the impact of radial nuclear divergence on trait-specificity, we used high-resolution data of chromatin radial positions (*80*) (**fig. S23C and Table S28**). Remarkably, BMI-PGEs were preferentially located in the nuclear periphery while T2D-PGEs were enriched in the nuclear core (**Fig 5B**). Furthermore, HARs displayed both peripheral and nuclear specific regions, while validated metabolic-HARs localized in central to peripheral regions (**fig. S23D**). Indicating a higher degree of tuning during transcriptional activity, clusters of social regulatory regions were enriched at the nuclear core, with some exceptions for peripheral isolated promoters (**Fig 5C**). To integrate functional radial divergence during transcriptional activity, we used multiplexed FISH imaging data derived from single cells active loci (*77*). This revealed direct relationships between nuclear organelle association, distance to nuclear lamina and transcriptionally active loci (**Fig 5D-E, fig. S23E-F and Table S28**). Interestingly, functional annotation of active nuclear core loci associated with nuclear speckles showed transcription of genes regulating cell differentiation, commitment, and cell adhesion (**Fig 5E**). Similar to functional enrichments for metabolic-HARs, we observed PPARG-mediated adipogenesis in core active genes (**Fig 5E**). We next determined functional network enrichment of interacting transcriptional regulators sharing TFBS-radial correlations (*80*) (**fig. S23G)**. Regulators with core TFBS-associations were highly social proteins as they display a larger proportion of physical interactions (**Fig 5G and Table S29**) Network reconstitution and functional enrichment revealed that core transcriptional regulators are related to nuclear receptors, PPARs, lipid metabolism, circadian clock, cell commitment and adipocyte differentiation (**Fig 5H and Table S29)**. Notably, interacting peripheral regulators were enriched in anti-inflammatory pathways mediated by several cytokines such as IL21, IL7, IL15, IL4 and IL13 (**fig. S23H**). Interrogation of radial dynamic expression during metabolic challenge revealed transcriptional peak differences between core and peripheral genes, with acute core activity and prolonged transcriptional behavior for peripheral genes (**Fig 5I and fig. S23I-J**). Similar to nuclear-cytosolic pH discrepancies (*81*) and supporting a nuclear core susceptibility to excessive protonation, their dynamic behavior was concomitant to elevated acidification of cytosolic pH and a threshold-dependent correlation of core activity with pH variations (**Fig 5J**). Collectively, these results indicate that structure-function dependencies translate environmental variations into topological specialized adaptations. This is supported by observations on radial divergence of risk-to-disease polygenicity (peripheral-to-core), location of metabolic-HAR endurers and spatial dynamic bursts influenced by pH-gradient.

### Transition from liquid-like to gel-like condensates modulates fasting endurance

We next determined radial condensate composition by assessing context-dependent conformations from regulators with TFBS-radial enrichment. Core-enriched regulators showed a higher number of sequences whose condensate globule behavior depends on local environment, such as pH variation and heterotypic interactions (**Fig 5K and Table S30**). In agreement with evolutionary adaptations to protonation (*8, 16*), correlation of IDR-ensemble parameters showed that core regulators display less segregation of charged residues and more negatively charged amino acids (**Fig 5L and fig. S23JK-L**). These enrichments highlight radial adaptations to environmental variations without precluding their activity in both compartments as observed in some metabolic-HAR regions and in DNA-MERFISH data. Among the environmental parameters, we confirmed that protonation regulates heterotypic co-activators and transcription factors condensates, which were perturbed by excessive protonation (**Fig 5M**). These results suggest that nutrient deprivation promotes adaptations at different levels to preserve cellular function (**Fig 3**). For example, fasting-mediated cytosolic acidification prompts heterotypic condensate formation guided by genome architecture to regulate the expression of genes dampening excessive cellular acidification (**Fig 3**). With both aspects showing evolutionary optimization (**Fig 1, 2 and 4**). This, however, does not fully explain if additional nuclear adaptations give mammals their ability to withstand chronic periods of nutrient deprivation. To elucidate this physiological process, we assessed condensate formation and dynamic properties during nutritional stress in living cells. Remarkably, FRAP experiments revealed that 6h of fasting promotes the formation of large and still dynamic PGC1A condensates (**Fig 5N-O**). Formation of condensates with multivalent interactions and different material properties have been previously reported during diverse stress conditions (*8, 82, 83*). For example, gel-like phase transitions by p-granules promote cellular fitness (*8*). Strikingly, fasting shifted PGC1A condensates dynamic behavior and recovery (**Fig 5P**), suggesting a fasting transition from liquid-like to gel-like conformations (*8, 16*). To confirm that these liquid-to-gel conformations are regulated by environmentally tuned electrostatic interactions, we found that their recovery decreased with addition of 1, 6 hexanediol, excessive acidification and acylcarnitine surplus (**Fig 5P**). In addition, we observed that, during fasting, co-activator condensates are much larger in size, and are regulated by pH-mediated ionic strength and salt concentration (**fig. S24A**). Similar to our observations, high-to-low entropic shifts suggest that gel-like formation ensures less diffusion of the transcriptional machinery required for physiological endurance (*8, 16*). To test if protonation alone can modulate these phase transitions, we blocked the cell-surface proton pump extruder HV1, previously shown to display metabolic functions (*84*). Similar to fasting, HV1 inhibition provoked cytosolic pH acidification (**fig. S24B**), dynamic gel-like condensate formation (**Fig 5Q**), and activity of selected metabo-HAR genes (**fig. S24C**). Next, we tested the heterotypic nature of these assemblies after perturbation of interacting transcriptional regulators. This revealed that, similar to fasting, HV1 inhibition promoted PGC1A gel-like organizations with longer times of recovery, which further decreased when interacting structuring regulators were perturbed (**Fig 5R and fig. S24D**). Suggesting a threshold-dependent effect of protonation on co-activator assemblies, condensates in prolonged fasting alone (18 hours) were still dynamic, while together with proton pump inhibition blunted their recovery (**fig. S24D**). By inhibiting condensate formation with hydrotropes such as ATP, we further showed that they buffer cytosolic pH acidification during resource conservation (**fig. S24E-F**). This threshold susceptibility indicates that optimization of heterotypic organizations buffering environmental variations might be favoured by evolution. Indeed, our results on PGC1A IDRs low conservation (**Fig 1F**), along with core-specific transcriptional adaptations (**Fig 5J**), prompted us to assess the fraction of charged residues and their degree of conservation (**fig. S24G-I**). Remarkably and contrary to core-enriched regulators (**Fig 5L**), PGC1A-IDRs displayed a higher proportion of positively charged residues (**Fig 5S and fig. S24H-I**), which, compared to negative residues, were 3 times as likely to exhibit low conservation (**Fig 5S**). This suggests evolutionary innovation on charge optimization for hub co-activators helps withstand acidic environments while preserving dynamic cooperative behavior.

### Metabolic resilience by robust genome plasticity

Genome architecture acts as a framework to guide the activity of transcriptional assemblies in response to environmental stimulus (**Fig 3-5**). Association of transcriptional factories with trans-interacting chromatin domains increases cell-specific contacts, ensuring the preservation of cellular function through the regulation of distant gene programs (*85*) (**Fig 2, 3 and 5**). To address the dependencies between disease hubs and dynamic architectural regions, we interrogated hub-regions of polygenic risk in relation to trans-interacting chromatin regions (**fig. S25A**). First, we defined social interchromosomal regions (SIRs), by using Hi-C data (*42, 86*) and data from crosslinked-based genome-wide detection of interaction hubs (*87*) (see structural prioritization of metabolic disease hubs). We complemented this by curating trans-interacting domains across different cell types, and obtaining a likelihood-based score for these social chromatin regions (**fig. S25B and Table S31**). Similar to metabolic-HAR regions, disease hubs from polygenic metabolic traits are preferentially located in SIRs (**fig. S25C**). Reconstruction of SIR locations into a chromatin-chromatin loci network of interacting disease hubs and metabolic-HAR regions (**fig. S25D-E**), revealed hierarchical maps of codependent disease hubs with potential systemic effects (**Fig 6A and fig. S25F**). Interestingly, metabolic associated FTO loci was found to interact with top trans-interacting nodes and top driver regions of metabolic-related phenotypes (**fig. S25D-G and fig. S15D**). BMI and T2D GWAS loci, located distantly, were topologically related between them and with validated metabolic-HAR domains (**fig. S25H**). This underscores the dependencies of polygenic metabolic hubs, their inter-trait relationships and their hierarchies. In addition, we found an overall enrichment for binding combinations of the PGC1A-interacting network to metabolic diseased genes within interacting hubs (**Fig 6B and Table S32**), and with overlapping metabolic-HAR regions (**fig. S26A**). In agreement with our observations (**Fig 2 and 5**), ATG4C-associated domain was found to integrate peripherally-enriched BMI hubs and shared BMI-T2D hubs with nuclear core locations (**fig. S26A**). For PGC1A-regulated genes during fasting, we found that the transcriptional activity within interacting disease hubs is higher compared to active genes within neighboring non-interacting regions (**fig. S26B-C**). Evaluation of disease variant hubs with mammalian phenotype-gene perturbation networks in syntenic domains revealed metabolic clusters, with several gene drivers (**fig. S26D-H and Table S31**). Module interaction analysis, syntenic conservation, and combined positional gene enrichments revealed territorial coalescence for metabolic phenotypes in specific regions (**fig. S26I-K**). Interestingly, this showed species-specific domains as only 15% of human disease hubs display syntenic regions with metabolic phenotypes in mice (**fig. S26L-N**).

**Fig. 6.**
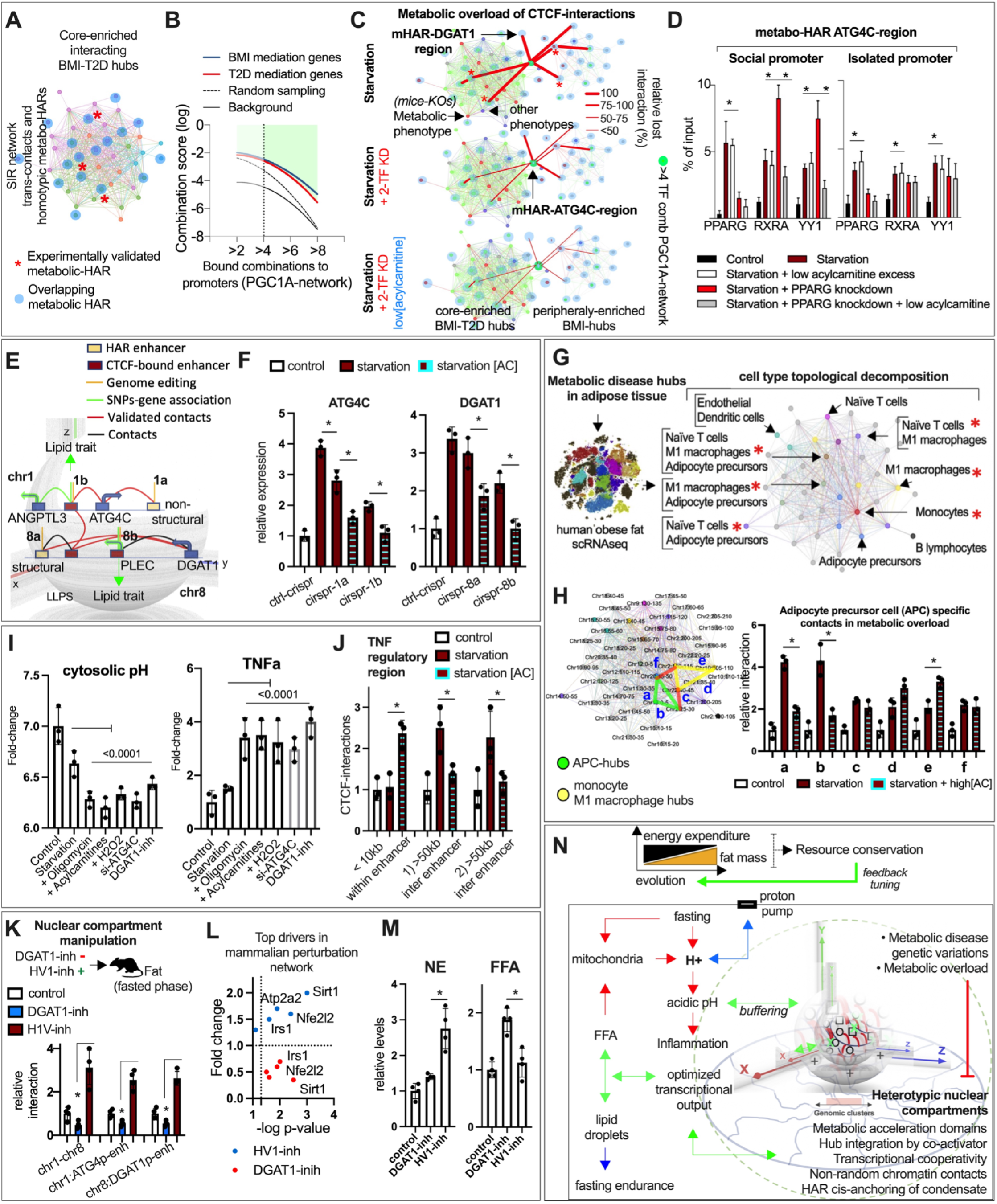
Resilience through robust genome plasticity. (**A**) Cell-naïve chromatin loci-loci network of trans-domains interactions with colocalized metabolic disease hubs, and related metabolic-HAR regions (in blue). With an asterix, experimentally validated HAR-domains. (**B**) Combinatorial regulation of mediator genes of polygenic risk for BMI and T2D by PGC1A-interacting regulators. (**C**) Human adipocytes CTCF-mediated long-range contacts in starvation with siRNA knockdowns (PPARG and RXRA) and low acylcarnitine overload (1uM). (**D**) ChIP-qPCR of social and isolated promoters within ATG4C-PGE region. Human adipocytes after starvation, with siRNA-knockdowns and metabolic overload with acylcarnitine (1uM). (**E**) Network reconstruction of structural dependencies between metabolic-HARs ATG4C and DGAT1 regions, location of related metabolic variants with associated genes, regulatory elements and coordinates (1a, 1b, 8a and 8b) for genome editing with CRISPR/Cas9 system. (**F**) Gene expression in human adipocytes displaying targeted mutations, in starvation and with acylcarnitine (10uM). (**G**) Single-cell topological decomposition of interacting (through trans-domains) metabolic disease hubs with scRNAseq from obese human subjects (*90*). With an asterisk, inflammatory cell hotspots. (**H**) Left panel, selection of cell-specific hubs subnetwork for adipose precursor cells (APCs in green) and topological neighboring subnetwork for inflammatory cells (in yellow). CTCF-mediated long-range interaction validation in human adipocytes in starvation and with acylcarnitine excess (20uM). (**I**) Cytosolic pH measurement in human adipocytes after starvation and starvation with oligomycin, acylcarnitine, H2O2, DGAT1-inhibitor and ATG4C-knockdown. Left panel, TNFa gene expression in human adipocytes in the same conditions. (**J**) CTCF-mediated interactions in TNF regulatory region. (**K**) Top panel shows illustration for chronic in-vivo intervention in mice by intraperitoneal injections with either DGAT1-inhibitor or HV1-inhibitor (2mg/kg). In the lower panel, CTCF-mediated (HAR regions ATG4C and DGAT1, and local enhancer-DGAT1_promoter) and YY1-mediated (enhancer-ATG4C_promoter) interactions in adipose tissue. Gene expression profile (**L**), norepinephrine and free-fatty acids (**M**) in adipose tissue in the same conditions as in K. (**N**) Summary illustration. Genome plasticity encodes metabolic resilience. Bars show mean values and error bars indicate SEM. Unpaired, two-tailed student’s t-test was used when two groups were compared, and ANOVA followed by fisher’s least significant difference (LSD) test for post hoc comparisons for multiple groups. * p-value < 0.05.

To assess the effects of metabolic overload and combinatorial perturbation of regulators on interacting disease hubs, we evaluated long-range contacts after fasting. Remarkably, nuclear compartment formation helped withstand both metabolic overload by acylcarnitine and gene perturbations of cooperative regulators (**Fig 6C and fig. S27A-B**). Targeted validations on ATG4C-domain showed that, in social promoters, single perturbation of PGC1A-interacting transcription factor increased the binding of other cooperative regulators (**Fig 6D**). Notably, this resilient response was lost by added metabolic overload with acylcarnitine (**Fig 6D**). Consistent with our results (**Fig 2-3**), isolated promoters do not display this genome plasticity (**Fig 6D**). Strikingly, genes from mediation analysis of polygenic risk scores and those regulated by PGC1A-network were preferentially social promoters (**fig. S27C-D**). Transcriptional burst profiles with Monte Carlo models showed that genes with social promoters are more susceptible to acylcarnitine excess during fasting (**fig. S27E**). Genes with social promoters and located at the nuclear core exhibited resilience to moderate acylcarnitine and a steep loss of burst amplitude by increased metabolic overload (**fig. S27F**). On the other hand, genes with social promoters located at the periphery displayed resilience to high acylcarnitine levels during fasting (**fig. S27F**). Moreover, perturbation of ATG4C and DGAT1 genes sensitized adipocytes to metabolic overload, impairing chromatin contacts and functional response to starvation (**fig. S27G-K**). Further dissection of this circuit by genome editing of HARs, as well as interacting CTCF-bound enhancers with related GWAS associations (**Fig 6E and figs. S28-S29**), showed that HARs enable the cells to withstand metabolic overload by keeping key structural enhancers in place for gene expression (**Fig 6E and fig. S30A-C**). Mutation of key structural enhancers impaired functional response to starvation, low expression of feedback regulators, and promoted a transition from transcriptional hub regulation to expression of genes with isolated promoters (**fig. S30I-J**). Remarkably and underscoring the functional plasticity of this regulatory switch, these genes are pro-inflammatory, inhibitors of lipolysis, and are causal mediators of lipid accumulation disorders (*88, 89*) (**fig. S30J-K**). Thus, perturbation of social genome plasticity revealed a switch to transcription within remote areas, seen locally in ANGPTL3, and in non-interacting neighboring segments (chr1:55mb) with PCSK9 expression (**fig. S30K**). This shows nutritional stress integrates structure-function adaptations, and how perturbation of social genome plasticity leads to expression of disease associated genes.

### Metabolic overload inhibits tissue- and cell-specific genome plasticity

To evaluate the cell-specificity of interacting polygenic hubs, we decomposed BMI-T2D fat hotspots with scRNAseq data from obese individuals (*90*) (**fig. S31A-B and Table S33**). Single cell hubs overlapping with the disease-SIR network revealed cell-specific, heterogeneous and promiscuous transcriptionally active trans-domains (**Fig 6G, fig. S31C-D and Table S33**). Given the overrepresentation of immune inflammatory hotspots, shared with APCs, we tested ectopic transitions during metabolic overload (**Fig 6G and fig. S31D**). Remarkably, during fasting, cell-specific transcriptional activity increased the interactions in contiguous trans-domains, which were lost and switched to immune hubs by metabolic overload (**Fig 6H**). This suggests cell-specific transcriptional regulators condition the contact probability of underlying architecture where ectopic expression of nonspecific genes might lead to tissue pathology (*91*). Given that ectopic expression of inflammatory genes in stromal or precursor cells affects tissue homeostasis (*92*), we determined APC function in additional harsh environments. Interestingly, metabolic overload, mitochondrial inhibition, ROS, ATG4C perturbation and DGAT1 inhibition, all led to excessive cytosolic pH acidity and elevated inflammatory gene expression (**Fig 6I**). Interestingly, TNFA, an inflammatory cytokine expressed in adipose tissue and related to insulin resistance (*93*) is located within the only nuclear core genomic-region of Chr6 (**fig. S12D**) Consistent with environmental regulation of inflammatory gene expression, burst kinetics of TNFA were similar to isolated genes, despite being located on a social regulatory region (**fig. S31E**). This indicates inflammatory genes exhibit transcriptional resilience to harsh environments likely due to homogeneous transcriptional regulation and specific genome plasticity. Remarkably, fasting metabolic overload by acylcarnitine excess showed a shift from long-range interactions to short-range enhancer-promoter entanglement in TNFA social regulatory region, suggesting again a social to nonsocial structural plasticity in metabolic overload (**Fig 6J and fig. S31F**).

To test heterotypic nuclear compartment formation in vivo, especifically in adipose tissue and APCs, mice undergoing fasting were treated intraperitoneally with an acute dose of HV1 proton pump and DGAT1 inhibitors. Fasting promoted metabolic-HAR ATG4C-DGAT1 domain interactions, and short-range contacts (**fig. S31G**). These were blunted by co-treatment with proton pump and DGAT1 inhibition, suggesting a gradient protonation threshold (**fig. S31G**). Chronic manipulation of compartment formation with 7 days daily-dose of either inhibitor revealed proton pump inhibitor increases selected metabolic-HAR compartments in adipose tissue (**Fig 6K**). Given that phenotype-gene perturbation networks of syntenic diseased hubs revealed top mediators genes including SIRT1, NFE2L2, and IRS1 (**fig. S26F**), we found that, in agreement with their function on metabolic homeostasis (*94–96*), HV1 inhibition increased their expression in adipose tissue (**Fig 6L**). We also observed opposing results with chronic HV1 inhibitor improving sympathetic tone and metabolic parameters, along with anti-inflammatory, mitochondrial and metabolic-HAR gene expression profiles (**Fig 6L-M and fig. S31H-L**). Chronic DGAT1-inhibition promoted a robust shift to pro-inflammatory and ER-stress gene expression (**fig. S31L**). These results confirm that re-esterification of lipids during metabolic overload is a mammalian response to increase resilience and fitness through protection of nuclear compartment tuning (**Fig 6N**).

## Discussion

In conclusion, we provide evidence of evolutionary coupling between phenotypic and molecular adaptations. Human acceleration of energy storage in fat tissue demonstrates the impact resource conservation has had on organismal adaptations seen in foraging behavior, cognition, physical endurance and reproduction (*97*). Coupling metabolic endurance with molecular optimization across transcriptional condensates, 3D architecture and cis-regulatory regions provides a framework for understanding evolutionary innovation, for example, in tuning nuclear self-organization. By studying these transcriptional dimensions, in relation to domains harbouring human specific substitutions, we show that environmental variation integrates components of metabolism into genome function. In addition, we show this has direct effects on genome plasticity and robustness, which could have many implications on a wide range of disease processes. We also demonstrate that fasting can recapitulate at small scales molecular adaptations taking shape throughout evolution. The pervasive nature of genome architecture coupled to cell-specific transcriptional machinery offer a way to manipulate specific subnetworks with systemic effects. It also provided us with a structural framework for multi-modal data integration. By using the probabilistic and combinatorial behavior of genome function, including protein-protein, protein-chromatin, chromatin-chromatin interactions, we use graph networks and genome proximity for integration of multi-genomic data. These nested structural representations allow us to model and funnel computational insights into experimentally testable hypotheses. With these, we demonstrate how genome function follows intrinsic architectural principles and drives cell-specific usage of the interacting architecture. For example, by integrating metabolic-driven pH variations into large heterotypic assemblies, cells optimize the transcriptional output by increasing genomic spreading through cooperative behavior with structural binders CTCF and YY1. This optimization is coupled with the usage of trans-interacting chromosomal regions harbouring functional related gene programs. One of these programs shown here, lipid-cycling genes, also harbouring human specific substitutions, is relevant for resilience to nutrient limitation. Indeed, fasting-mediated fatty acid mobilization leads to fatty acid oxidation, which cannot be sustained unhindered. This promotes the accumulation of fatty acids, inhibition of mitochondrial activity, and further decrease of cytosolic pH. Integrating proton excess to chromatin-condensate stability for the regulation of gene programs controlling cytosolic acidification provides evidence of a robust feedback integrator. The evolutionary tuning of chromatin structure and heterotypic condensates for optimal bursts of lipid cycling genes promotes metabolic and oxidative stress endurance. It also demonstrates the encoding of resource conservation in genome function, and more broadly evolutionary innovation in parametrization of self-organization.

Mammalian comparison of syntenic HAR-domains show some degree of convergence in long genomic-ranges, and acceleration of tuning in short-range scales. It also shows that territorial transcriptional activity is preserved in mice and is congruent with genomic-range phenotype associations, similar to metabolic disease variant hubs. Our topological hub integration reveals that high-fat diet promotes loss of transcriptional control by metabolic overload through accumulation of fatty acids, and excess of acidification. Moreover, we provide evidence that radial chromatin organization pairs structure and function. Resource-driven adaptation to protonation is observed in the enrichment of negatively charged amino acids in nuclear core transcriptional regulators. Despite this radial plasticity, the co-activator PGC1A, acting as a condensate-integrator in acidic milieu during energy conservation, shows more evolutionary innovation for positively charged residues. This suggests tuning of condensate endurance as excessive protonation impairs its dynamic properties. These features license the formation of large heterotypic organizations with gel-like properties that buffer proton excess, withstand less molecular diffusion and remain reversible self-organizations. We show that their reversibility and dynamic properties depend on the presence of heterogeneous interactions as perturbation of interactors reduces their transition ability. Moreover, HAR substitutions have been shown to be enriched in AT-to-GC transitions near telomere regions, which is in line with observations on nuclear core enrichments (high in GC-content). Interestingly, specialized metabolic-HAR regions regulating fasting endurance are located towards central-peripheral nuclear regions. Their radial 3D location implies their need to withstand acidic excess during activity to promote a resilient phenotype. Below a physiological threshold, acidity not only dissolves heterotypic assemblies and chromatin interactions, but also leaves unaffected pH-resilient inflammatory gene programs. The latter is relevant as transcriptional assemblies have been found in proximity to interchromosomal regions. We therefore modeled the topological dependencies of metabolic disease hubs from mediator genes of polygenic risk with a cell-naïve trans-interacting chromatin network. This shows the pervasive nature of structural architecture in classifying hierarchical disease hubs with systemic effects and their inter-trait relationships. We further decompose these hotspots into cell-specific transcriptional hubs using single-cell RNA seq data. At this resolution, we show that cell-specific transcriptional activity drives cell-specific usage probability of underlying chromatin interactions. This cooperative plasticity, in both long-range convergence and short-range tuning, was dependent on metabolic overload and cytosolic pH. We further demonstrate, in vitro and in vivo, that pH reduction within physiological ranges through HV1 pump inhibition can promote a metabolic resilient response with heterotypic condensates, and genome architecture, which promotes organismal, tissue and cell-specific homeostasis. On the other hand, inhibition of fatty-acid esterification, and perturbation of social genome plasticity through HAR-circuit genome editing, block selected nuclear compartments in adipocytes and adipose tissue, promoting inflammation and metabolic malfunction.

In summary, our work uncovered mechanisms of evolutionary innovation in tuning transcriptional self-organization. While these mechanisms are recorded through evolution into transcriptional hubs within chromatin architecture, dynamic mechanisms such as stress-dependent phase-separation allow the integration of these uni-dimensional regions into multi-dimensional platforms consisting of metabolic-HAR domains, their master regulators and local thermodynamic states. The present study therefore highlights the existence of structural and topological mechanisms unraveling hierarchically-organized functions stored within the genome. In particular, it revealed robust feedback integrators of resource-driven adaptations, and general approaches for manipulating genome function, for example, through the use of compartmentalized hub-models for precision medicine and machine learning. Such computational approaches could benefit from a better understanding of the principles governing functional compartmentalization, for example, of stress-dependent transcriptional hubs and how these hubs are used as a feedback to stress. In parallel work (*98*), we have investigated mathematically and computationally, principles through which cellulhar self-organization, and more generally compartments, can be manipulated in stress conditions at different entropic states, to optimize resilience to pressure inputs, while acquiring properties of self-supervised organization (*98*). In all, our work suggests that metabolic input overload offers a therapeutic opportunity for obesity, metabolic disorders and many pathological processes by promoting stress resilience through genome plasticity.

## Supporting information

Supplementary Materials for Metabolic Resilience is Encoded in Genome Plasticity

## Acknowledgments

We thank all the members of the computational biology group at MIT, the Femenia lab members at the institute for Neuroscience in Alicante, and the Novo Nordisk research unit at Seattle for discussions and help throughout the research activities described here.

## Author contributions

Project design and Conceptualization: LZA, RT. Computational Methodology: LZA with feedback from RT, MK, PWF, NAP, YP, TF, NS, BK, KG. Polygenic risk scores and mediation analysis: YP with feedback from LZA, MK. Experimental design and validation: LZA and TF with contributions from CL, AM, RT, SKL, LH, KG, PWF, KG, BK, MK. Supervision of the work: LZA, MK, RT, TF. Funding acquisition: LZA, MK, RT, TF. Writing of original draft: LZA. Writing & editing: LZA, TF, RT, MK. All authors reviewed the manuscript.

## Data and materials availability

Data is available in the supplementary materials. Accession numbers to any data relating to the paper will be deposited in a public repository

