## Supplementary Materials for Metabolic Resilience is Encoded in Genome Plasticity for "Metabolic resilience is encoded in genome plasticity"

#### This pdf file includes

Materials and Methods
- Experimental methods
- Computational methods

Figures S1 to S31
Tables S1 to S33
Captions for Tables S1 to S33
References and notes

### Materials and Methods

#### 1. Experimental material and methods

##### Animal experiments

All animal experiments were performed in accordance with internationally accepted guidelines and principles for the use of laboratory animals, and were approved by the Novo Nordisk Research Centre Seattle Institutional Animal Care and Use Committee and the Novo Nordisk Ethical Review Committee, Novo Nordisk Research Center in China, and by the regional ethics committee of Alicante, and Universidad Miguel Hernández-CSIC Spain.

Adult Wild-type male C57BL/6J mice aged 6-12 weeks were obtained from Jackson Laboratories (Stock # 000664). Mice were housed 4-8 per cage with ad libitum access to water and food (Type IV Scanbur Cages; 1800 $cm^2$ floor area). Animal holding rooms temperature ranged from 20-26 °C and humidity ranged from 30-70%. Lighting was on an automated light/dark cycle with lights on between 6:00 AM and 6:00 PM. Mice were provided with Alpha Dri bedding material to just cover the bottoms of cages cage, one Nestlet square, and two 8 g Bed-r'Nest "pucks" or similar. Environmental enrichment included a plastic hut, a paper hut, one wood gnawing block, and a tunnel suspended from the top of cages. Mice received a hydrogel cup upon arrival until familiarized with the automatic water system which supplied reverse osmosis purified water via the Scantainer rack system. Standard rodent chow diet [PicoLab Rodent Diet 20 Lab Diet #5R53 (24.5% Protein, 13.1% Fat, 62.4% Carbohydrate)] was provided to the mice upon arrival until diet was switched from chow to high fat (described below). All food and water were available ad libitum. Mice were acclimated to housing for one week and then marked with 2D barcode ear tags.

<u>High fat diet experiments</u>: At 4 weeks of age mice were placed on a 60% high fat diet (High-Fat Diet #D12492 (20.0% Protein, 60% Fat, 20% Carbohydrate) during 10 weeks and they reached dietary induced obese status (DIO). After this point, mice were moved from group housing to 2 mice per Type IV Scanbur cage, separated by an opaque cage divider. Bedding and environmental enrichment was as described above. Food and water were provided ad lib regardless of housing condition. Mice were sacrificed and tissues of interest were extracted and kept at -80 °C until RNA extraction protocol. Once RNA per tissue was obtained it was sent for RNA-sequencing at the Covance Genomics Lab, Redmond, WA. cDNA libraries were prepared with Illumina TruSeq stranded mRNA kit and sequenced on an Illumina HiSeq. 2000 using paired-end, 100 nucleotide reads with an average depth of 20 million reads per sample. Transcriptomic profile from 19 tissues, comparing control chow diet and HFD diet interventions were carried out. Differential expressed genes from each tissue were obtained following previously established protocols (*98*). This was followed by integrative topological genomic protocols described in this paper.

Wild-type mice of 12 weeks received intraperitoneally 1 daily dose during 1 week of either HV1 proton pump inhibitor (2.mg/kg 2GBI, Sigma Aldrich), DGAT1 inhibitor (2mg/kg Sigma Aldrich, CatNo:A922500) or control. On day 7, during their fasting phase mice were sacrificed. This was followed by tissue extraction and further processing for molecular experiments. In addition, wild-type mice of 12 weeks were fasted for 12 hours. In parallel, a subgroup of mice received intraperitoneally 1 single dose of either HV1 proton pump inhibitor (2.mg/kg 2GBI, Sigma Aldrich), DGAT1 inhibitor (2mg/kg Sigma Aldrich, CatNo:A922500) or control. At the end of the fasting period mice were sacrificed. This was followed by tissue extraction and further processing for molecular experiments.

##### Mammalian cell culture

Human embryonic kidney 293 cells (HEK293) were cultured on glass-bottomed MatTek dishes. Cells were grown in Dulbecco's modified Eagle's medium (Gibco, Thermo Fisher Scientific), 10% fetal bovine serum (Thermo Fisher Scientific) and 10U/mL penicillin-streptomycin (Thermo Fisher Scientific) and incubated at 37ºC and 5% $CO_2$ in humidified incubator. Cells were transfected when they reached 70% confluence with plasmid DNA GFP-PGC1 (Addgene, Massachusetts, USA) using Lipofectamine 3000 (Invitrogen, Thermo Fisher Scientific) and removed 24h post-transfection, according to standard manufacturer protocols. Transfected cells were imaged 24-48 h post-transfection. Human primary pre-adipocytes (ATCC) were differentiated into mature adipocytes and used for fasting and chemical treatment experiments. For mice primary adipocytes, SVF from inguinal fat depots of 6-12-week-old male mice were isolated and prepared as previously described (*99*). The differentiation cocktail was used during the first 2 days of culture, followed by rosiglitazone and Insulin for 6 to 8 days.

##### Cell and chemical treatments

Indicated cell cultures were kept in different condition media: Dulbecco's without glucose for fasting condition or Dulbecco´s high glucose medium for control. Adipocytes under differentiation were transfected using lipofectamine 3000 (Invitrogen, Thermo fisher technologies) with siRNA for PPARα (Applied biosystems), with siRNA for YY1 (Applied biosystems), with siRNA for PPARG (Applied biosystems), with siRNA for PGC1A (Applied biosystems), siRNA for RARA (Applied biosystems), siRNA for RXRA (Applied biosystems), with siRNA for CTCF (Applied biosystems), with siRNA for

EP300 (Applied biosystems), with siRNA for ATG4C (Applied biosystems), with siRNA for DGAT1 (Applied biosystems), with siRNA for INSIG2 (Applied biosystems), with siRNA for CAT (Applied biosystems), and scramble control. Once they were differentiated, they were processed for analysis of gene expression, or follow up experiments (as indicated). HEK293 cells were transfected as previously indicated with siRNAs or scramble control 24h previous to downstream experiments. Cells were treated when indicated with 2-Deoxy-Glucose at 0.25mM (Sigma Aldrich) to inhibit glycolysis. 2-guanidinobenzimidazole at 200uM (2GBI, Sigma Aldrich), was used to inhibit the proton pump extruder HV1 as indicated. DGAT1 specific inhibitors were used as indicated (Sigma Aldrich, CatNo:A922500). C:16-acylcarnitine was used as representative long-chain acylcarnitines for cellular experiments at different concentrations as indicated (Sigma Aldrich 61251 - Palmitoyl-L-carnitine). Etomoxir (Sigma Aldrich, 50uM) was used to block CPT1A-B mitochondrial fatty acid import channel as indicated. 1,6 Hexanediol at 2.5mM was used to dissolve condensates (240117, sigma aldrich)

**Analysis of Gene Expression**

Total RNA was isolated from cells or tissues using Isol-RNA Lysis Reagent (5 PRIME) according to manufacturer's instructions. Amplification Grade DNase I (Life Technologies) was used to treat 1 µg of RNA, from which, 500 ng were used for cDNA preparation following Applied Biosystem Reverse Transcription Kit (Life Technologies). Quantitative Real-Time PCR was performed in a ViiA 7 Real-Time PCR system thermal cycler with SYBR Green PCR Master Mix (both Applied Biosystems). Analysis of gene expression was performed using the ΔΔCt method and relative gene expression was normalized to hypoxanthine phosphoribosyltransferase (HPRT) mRNA levels. Gene expression analyses were expressed as mRNA levels relative to controls. Primer sequences available upon reasonable request.

**ChIP immunoprecipitation**

ChIP experiments were performed as previously described (*100*) with the following modifications: Cells were plated and (5-10million) pooled before homogenization. Cells were homogenized in PBS and crosslinked with 1% formaldehyde during 10min. For tissue, 50mg were homogenized in PBS and crosslinked with 1% formaldehyde for 10 min. Glycine (125 mM) was added, followed by centrifugation at 4000 rpm for 5 min. After aspirating the supernatant, the pellet was washed with 1 ml cold PBS followed by centrifugation at 4000 rpm for 5 min. This step was repeated with buffer 1 (0.25% Triton X-100, 10mM EDTA, 0.5mM EGTA, 10mM HEPES, pH 6.5) and buffer 2 (200mM NaCl, 1mM EDTA, 0.5mM EGTA, 10mM HEPES, pH 6.5). The pellet was then resuspended in 300 µl lysis buffer (1% SDS, 10mM EDTA, 50mM Tris-HCl, 1X protease cocktail inhibitor, pH 8). Samples were sonicated on ice using a Bioruptor (Diagenode) 30 times for 30 seconds ON and OFF intervals. After the lysates were sonicated to shear the DNA to fragment lengths of 300bp, the complexes were immunoprecipitated with antibodies specific for CTCF (Abcam), YY1 (Abcam), PGC1A (Abcam), PPARG (Abcam), RXRA (Abcam), EP300 (Abcam), and H3K27Ac (Abcam). 10-20 ul of antibody was used for each immunoprecipitation. No antibody controls were also included for each ChIP assay and no precipitation was observed by quantitative Real-Time PCR (qPCR) analysis. Input samples were processed in parallel. The antibody/protein complexes were collected by either salmon sperm DNA/protein A agarose slurry or Protein A/G PLUS agarose beads (Santa Cruz sc-2003) and washed several times. The immuno complexes were eluted with 1% SDS and 0.1 M $NaHCO_3$ and samples were treated with proteinase K for 1 hour and DNA was purified by phenol/chloroform extraction, ethanol precipitation and resuspended in 20 ul $H_2O$. Input DNA and immunoprecipitated DNA were analyzed by quantitative PCR. Primer sequences used to detect each DNA region are available upon reasonable request. Primers are available upon reasonable request.

**ChIP-loop immunoprecipitation of chromatin interactions**

ChIP-loop-qPCR protocol is based on ChIP immunoprecipitation assay previously described with the following modifications: After sonication of the complexes, the lysate was digested HindIII (NEB) following manufacturer's recommendation. Then the digested complexes were immunoprecipitated with specific antibodies (see ChIP-immuno). The immunoprecipitated complexes were resuspended in 200ul of 1x T4 ligase buffer with 400 U of T4 ligase (NEB). Then the samples were collected by protein A/G Plus agarose beads, and DNA samples were purified with phenol/chloroform extraction and ethanol precipitation. Ligation efficiencies were determined using the qPCR assay compared with control templates. Specific primers for computationally selected loops were designed to validate chromatin-chromatin interactions. When comparing different biological samples, highly interactive chromatin regions were used as a control for normalization of sample effect. Positive interactions were matched by ChIP-binding of the regulator. Primers are available upon reasonable request.

**Kit measurements**

Triglyceride (Abcam ab65336), Free Fatty Acid (ab65341), ROS Detection (ab139476), NE elisa kits (myBiosource), and cellular pH (thermo fisher) quantification from biological samples were done using commercially available kits, following manufacturer's instructions.

**Protein sample preparation and turbidity assays**

Protein aliquots were thawed above the UCST to ensure the protein was soluble. Phase separation was induced by mixing (stored in 500 mM NaCl and 20 mM Tris, pH 7.5) with no-salt buffer (0 mM NaCl and 20 mM Tris, pH 7.5) to obtain a solution containing 150 mM NaCl and 20 mM Tris, pH 7.5. Protein concentrations were adjusted as necessary by diluting further with the same buffer conditions. Crowding was induced with different concentrations of PEG 8000 (Sigma Aldrich). Proteins were dissolved in potassium phosphate buffer (pH 7) to a concentration of 160 μM/800 μM as stock and diluted with potassium phosphate buffer (pH 7) in the assay. Then, PGC1A and RARA were mixed with 0, and 20% PEG 8000 solution with a volume ratio of 1:1 at room temperature to obtain various concentrations. Arginine (R) was dissolved in a potassium phosphate buffer (pH 7) to a concentration of 2, 1.5, and 2 M as stock, respectively. PGC1A-RARA combined or individual solutions were mixed with amino acid solution first to obtain various concentrations of R. Then, 20% PEG 8000 was added to the mixture to a volume ratio of 1:1. PGC1A-RARA combined or individual solutions were desalted into a potassium phosphate buffer of pH 5, pH 6.2, pH 7.5, or pH 9. Protein solutions were then concentrated into 320 μM/800 μM as stock and diluted with corresponding buffers in the assay. Then, protein solutions were mixed with 20% PEG 8000 solution in a volume ratio of 1:1 at room temperature to obtain various concentrations. Turbidity assays were performed on a UV-Vis spectrophotometer (Thermo Scientific, Evolution 350 UV-Vis Spectrophotometer). Samples (in quartz cuvettes with 1 cm path length; Thorlabs) were first equilibrated, and the instrument was blanked. Throughout the experiment, absorbance was measured at $\lambda = 600$ nm. All samples were examined in triplicate ($n = 3$).

**Live cell imaging and Fluorescence Recovery After Photobleaching (FRAP)**

FRAP was performed using an inverted confocal super-resolution microscope Zeiss LSM 880-Airyscan Elyra PS.1 with a 63x oil immersion objective. Cells were maintained in the microscope chamber at 37ºC and 5% $CO_2$. A circular region of interest (ROI) 23 mm was selected on GFP-PGC1 nuclear condensates and bleached with Argon laser at 488 nm wavelength by 10 consecutive bleaching iterations (1ms). Fluorescence intensity was recorded at baseline, during and after the bleaching and corrected by ROI corresponding to the background. Different media conditions were used as indicated; Dulbecco's without glucose (6-8h) for fasting condition or Dulbecco´s high glucose medium. When indicated cells were transfected with targeted siRNA for knockdown experiments followed by the aforementioned protocol. 1,6 Hexanediol at 2.5mM was used to dissolve condensates (240117, sigma aldrich). ATP at 1mM was used as a hydrotrope to evaluate condensate dissolution when indicated. Image processing and analysis was performed with ImageJ.

**Extracellular Flux Analysis (Seahorse) Assays**

Cells were differentiated and treated as described above. When indicated, mitochondrial oxidative phosphorylation from human adipocytes was analyzed using extracellular flux analysis (XF24; Seahorse Biosciences) in DMEM buffer (pH 7.4; Sigma Aldrich). Baseline oxygen consumption rates (OCR) were measured every 7 min for control and cells undergoing glucose fasting as previously described.

**Genome editing of regulatory elements by CRISPR/Cas9**

Circuitry dissection: Our integrative genomic approach revealed chromatin segments with nonrandom higher-order locations. We have characterized their functional role in resource-conservation endurance across many experimental systems. We have focused on nuclear compartments with similar function, with functional interactions both at short- and long-range levels, along with their metabolic disease risk associations. The latter is observed in unbiased genomic range genetics, and in unbiased epigenomic integration of loci-trait-tissue relationships, where both underscored the importance of inter-trait-disease dependencies for metabolic-related traits, especially lipid associated modules. We next tested in polygenic risk scores and downstream causal genes, the topological relations between T2D and BMI variant-hubs, and how these colocalize with metabolic-HAR domains, interacting chromatin regions, cell-specific transcriptional hotspots, driver regions of metabolic disease, especially lipid disease, and within selected functional compartments. This approach showed functional nuclear compartments for lipid cycling genes, fine-tuned by cis-regulatory human accelerated regions, heterotypic condensates and local thermodynamic states, such as resource limitation. The same functional regions display strong topological relationships to T2D-BMI polygenic risk variant hubs, bridging segments of chromatin architecture and regions of inter-trait and inter-disease dependencies by lipid-related modules. Our experimental validation confirmed this, and helped find top functional dependencies during resource conservation, such as chr1-ATG4C region and chr8-DGAT1 region. During fuel restriction, we showed that HAR-enhancers are more sensitive to metabolic overload by acylcarnitines, while key structural enhancers (CTCF-bound) formed promoter loops that are resilient to acylcarnitine excess. Regarding the latter, even though they preserved basal transcriptional activity of target genes in starvation, fatty acid excess decreased their burst profile. For these reasons, we dissected the local circuitry of this heterotypic compartment using genome editing of key regulatory elements (HARs and structural) in human adipocytes, followed by functional, structural and transcriptional assessment of their genome perturbation in starvation states. Our initial goal is to test our previous findings and find causality as to how HARs offer stability to social genome plasticity, and how structural enhancers are key for intra-domain stability during resource conservation. Second, we want to find context-dependent causality for loss of genome plasticity transitions, and how these are relevant for phenotype outcomes of metabolic genetic variants within the circuitry. Thus, we focused on the local interacting circuitry between HAR elements and promoter regions of target genes, followed by isolation of key CTCF-bound enhancers (this is in addition to experimentally validated loops such as YY1-mediated), with genetic variations related to metabolic traits, especially lipid disorders (as previously described in genomic-range genetics, epimap integration and mediators of polygenic risk for

T2D-BMI). ATG4C domain circuitry: following our validations, we focused on the HAR ANC162 that forms interactions with the promoter region. We carried out C-to-T genome editing using CRISPR/Cas9 in the binding motif (see transcriptional pipeline for TFBS-assessment) of a PGC1A-interacting target, YY1. As shown in our results in fig. S28 on circuit dissection, there is a genetic variation rs17124210 in that position, related to waist-hip-ratio (mutation name CRISPR-1a). Next, we focused on our previously validated key enhancer linked to ATG4C promoter, R5 (fig. S28). In starvation, this enhancer forms YY1-loops, and it is bound by CTCF as well. Given our results on heterotypic cooperative behavior of interacting transcriptional regulators, and how CTCF as a chromatin binder participates in this plasticity, we focus on finding CTCF-binding sites with genetic variations or in proximity. Using the DNA sequence of the whole regulatory element span, we scanned it for CTCF-motifs using Ziebarth et al. approach (*101*). Considering the syntax effect and influence of neighboring motifs, we selected for genome editing the closest CTCF-motif to the strongest genetic variation to lipid disorders, rs12130333. This position is associated with the ANGPTL3 gene, which is located within the ATG4C submodule (Fig 2 and 3). In this motif, we also carried out C-to-T genome editing using CRISPR/Cas9 in the binding motif (mutation name CRISPR-1b). DGAT1 domain circuitry: we have shown in our experimental validations that DGAT1 location in the tail of chromosome 8, lies in a social interchromosomal segment, and belongs to our selected nuclear compartment. We found in starvation, long-range CTCF mediated interactions both homotypic to CAT-domain and heterotypic to metabolic-HAR ATG4C domain. By using the network reconstruction of short-range interactions (Fig 2), we obtained DGAT1 territorial module and submodules. We then focused on the HAR-HACNS71 element as a candidate for DGAT1-circuitry, which, due to distance constraints (1mbp), was validated in Fig 3 as CTCF-mediated interaction. In close proximity to this HAR element, lies an active validated element_558 (VISTA), forming cis and trans interactions, as well as being bound by ChIP-seq experiments to CTCF and YY1, both members of PGC1A-heterotypic condensates. Therefore, we mapped top CTCF-motifs in this HAR element to perform C-to-T genome editing using CRISPR/Cas9 (fig. S29; mutation name CRISPR-8a). Next, we scanned regulatory elements within the DGAT1 submodule for genetic variations associated with phenotypes congruent with the trans-interacting ATG4C domain. We found in a regulatory cluster, a genetic variation rs55831924 associated with lipid disorders and related to PLEC gene. Unbiased mapping of CTCF-motifs in the regulatory element DNA sequence, revealed that the genetic variant colocalized with a top CTCF-motif. This motif was selected to perform C-to-T genome editing using CRISPR/Cas9 (mutation name CRISPR-8b). Full network reconstruction of circuitry dissection can be found in supplementary figures S28 and S29, and Fig 6E. Genome editing: we performed genome editing in human adipose cells as previously described (*102*). In brief, all mutations in selected regulatory elements (some with genetic variations; see previous section) were from CC to TT alleles. hCas9 and guide RNA (gRNA) vectors were used (Addgene). C-to-T change was done using site-directed mutagenesis Q5 kit (New England Biolabs). Guide RNAs were designed using the CRISPR design tool available at http://crispr.mit.edu/. GFP, Cas9 with sgRNAs, homology vector, and pMACs 4.1 plasmids were co-transfected in adipose progenitor cells using Amaxa-nucleofector (lonza). Cell sorting was done using MACSelectTM (Miltenyi Biotec), followed by cell culture for 6 days, and propagation of selected clones for downstream experiments. These were functional assays (FFA, ROS, TG, pH, OCR), targeted chromatin interactions (cis and trans within circuitry), expression of target genes (ATG4C and DGAT1) and topological outcomes from loss of social genome plasticity in resource-conservation states. For the last one, we were guided by the interaction frequencies among regulatory elements within the domain (defined in Fig 2 as social or isolated regulatory regions). Network or topologically-based prioritization of interacting modules was used to extract isolated regulatory regions (fig. S30I), followed by functional annotation, and detection of strong genetic variants in contiguous segments such as PCSK9 in Chr1:55mb (rs11591147). We then tested in cells with mutations in key structural enhancers (1b and 8b), if loss of social genomic plasticity in nutrient limitation leads to transition of transcriptional activity to remote areas or isolated regulatory regions within domains of interest. We selected top isolated region genes and quantified their transcriptional activity in the aforementioned conditions. Primers are available upon reasonable request.

**Intraperitoneal glucose tolerance test (IP-GTT)**

Intraperitoneal GTT was performed on overnight fasted mice. Blood glucose levels were measured at basal state (0 min) and then at 30, 60, and 120 min after i.p. injection of glucose (1.5 mg/kg body weight). Blood glucose concentrations were measured using the Accu-Chek Aviva monitoring system (Roche, Basel, Switzerland).

**Mitochondrial mass**

Plated cells were pooled and homogenized with PBS and Mitochondria Isolation buffer (225 mM Mannitol, 75 mM Sucrose, 30 mM Tris-HCl pH 7.4, 0,1mM EGTA). For animal experiments, 10 g of tissue was used from each animal to measure mitochondrial mass. Tissues were homogenized with Mitochondria Isolation buffer. Tissues were slowly homogenized with strokes to break them up to 80%–90%. They were spun down at 500 xg for 10’ to bring down unbroken cells and cell debris. The supernatant was then transferred to weighted Eppendorf tubes. This was followed by centrifugation at 7000xg for 10’ at 4°C to bring down mitochondria. Finally, the Eppendorf was weighted again and the difference between the last weight and the previous one gave us the total mitochondria mass.

**Mitochondria/Genomic DNA Ratio**

DNA was extracted using Isol-RNA Lysis Reagent (5 PRIME) according to the manufacturer's instructions. After DNA quantification, 3 nmol of DNA were used to quantitative real-time PCR using SYBR-green. We used primers for mitochondrial DNA and primers for hypoxanthine-guanine phosphoribosyltransferase (HPRT) as a gene specifically transcribed in the nucleus. We used the ratio between the mitochondrial DNA and genomic DNA as an indication of the mitochondrial DNA per sample. Primers are available upon reasonable request.

#### 2. Computational methods

##### Integrative topological genomics (ITG)

##### General description

Genome features including activity, structure and diversity are being measured at an exceptional pace. These multi-layered representations are modeled with single statistical approaches or with ensemble machine learning methods. Despite recent advances, biological systems, data generation, and analysis are imprinted with stochasticity, which coerces the generation of general models. On the other hand, multi-models of stacked single tasks are poised to better capture this diversity, as they funnel computational inferences into guided representations of genome proximities. To elucidate genome structure-function dependencies, and given that nuclear architecture provides a structural framework for genome function, we use here structural representations of genome distances to supervise the integration of multi-layered information (e.g. structural features, protein interactions, binding of regulatory regions, heterotypic condensates, tissue and cell-specific gene expression profiles, genome substitutions). These pipelines are categorized in several classes (evolutionary, transcriptional, structural, comparative genomics, disease hubs, and radial genome), each with its own focus, but complementing each other (**fig. S1 and S2**). Every pipeline has points of transition between them, and points complemented by experimental validation of computational inferences (**fig. S1 and S2**).

##### Integrative topological genomics I (ITG-I): evolutionary relationships

- **Multi-species metabolic phenotypes**

Data for multi-species comparisons of metabolic phenotype were obtained as follows: total energy expenditure (TEE) from, including hominidae fat and lean mass, and (*22*, *23*), experimental fat mass measurement across mammals (*24*) (table S1).

- **Multi-species endurance score**

To calculate energy endurance score, we followed previous work by Lindstedt et al. (*25*), where they generated a fasting endurance score (survival days in nutritional deficit) as a function of body size in mammals. We used a similar approach but with allometric (10% of body weight) and experimentally measured fat mass. Briefly, endurance score is based on the relationship between fat mass energy levels (in kilojoules) and the amount of energy consumed per day from TEE data (table S2). It is then derived from resource availability, assuming energy intake is low. This gives a representation of the number of days a given species can last solely on the energy stored in fat tissue. After obtaining an endurance score per species where data was available, we plot a XY graph together with fat mass. Fat mass is derived from both experimentally measured and allometric fat mass. We then performed linear regression to compare both experimental and allometric models.

- **Primate-Human comparison**

We used previous published work to compare primate and human data: from brain cells (*27*) and adipose tissue (*26*). We re-analyzed the data following protocols described there (GEO: GSM3494237–GSM3494249), and used differential signatures for downstream analysis.

- **Network-based module discovery**

To assess functional gene-gene dependencies in tissue-specific networks, we performed a targeted approach of overlapping network representation using data from tissue-integrated genome-scale analysis (*28*) (archived in http://giant.princeton.edu and https://hb.flatironinstitute.org/). This was followed by functional clustering of dependencies.

- **Integration Human accelerated regions (HAR)**

Human accelerated regions and HAR associated genes were obtained from previous work (*29*) (table S3). Full HAR-associated genes were completed with subsequent studies (*20*). Enhancer coordinates and HAR-genes were used for downstream analysis. HAR-genes were used as input for network-based module discovery in specific tissues, followed by interrogation of overlapping molecular signatures with results from primate-human high-throughput comparisons. Human gain enhancer positions were obtained from Uebbing et al. with TFBS enrichments and target regulated genes (*30*)

#### ITG-II: transcriptional relationships

- **Transcriptional regulators promoter-binding**

To assess transcriptional binding to promoters of HAR-associated genes, we used a combined enrichment approach including motifs and binding of regulators. For a given geneset we used conserved common regulatory motifs enrichment (archived in http://www.gsea-msigdb.org/gsea/msigdb/index.jsp). Enrichr in R was used to perform functional enrichment analysis based on the database: ChEA 2016 and ENCODE consensus. FDR <0.01 and p-value <0.05 were used as a threshold to select the significant enrichments (*28*, *103*) (archived in https://maayanlab.cloud/Enrichr/). Enrichment results from each class were followed by venn-diagram filtering of overlapping regulators. These were used for downstream analysis. Prediction tool for TFBS from DNA sequences can be found at http://alggen.lsi.upc.es/. Tool for prediction of CTCF-binding sites can be found at https://insulatordb.uthsc.edu/ (*101*).

- **Functional interacting transcriptional regulators (f-ITR)**

Common regulators from our previous step were used as inputs to identify physical interactions in Protein-Protein interaction (PPI) data stored in several repositories. First, we obtained physical interactions from regulators using human protein interaction atlas (archived in http://www.interactome-atlas.org/ (*104*)). This was followed by filtering of regulators displaying common molecular signatures. These were used for network recomposition

- **Network recomposition and modelling of f-ITRs**

Next, we carried out a 1+ node reconstitution (partial) and full network reconstitution of all possible interactors from our filtered targets. To do so, we used both HuRI and Biogrid PPI-data (archived in https://thebiogrid.org/ (*105*)). Partial and fully reconstituted networks were used for downstream analysis. We performed global and local network analysis with clustering and modularity followed by node-prioritization with eigenvector centrality and random-walks algorithms. Nodes with the highest influence on the network were used for functional annotation embedding using DAVID clustering (*106*) and HAR-association with our curated dataset. Top ranking nodes were filtered for downstream analysis.

- **Individual phase separation scores**

We next identify computationally intrinsically disordered regions in the amino acid sequences of f-ITRs. Disorder scores from protein amino acid sequences were obtained using predictors of naturally disorder regions software (*107*, *108*), inference of intrinsically unstructured proteins (IUPred) with Anchor (binding regions within IDRs) package (*109*)(*110*), multivalent pi-pi interactions in non-aromatic residues of folded proteins (*111*) and curated experimentally validated phase separation databases (stored at http://www.pondr.com/, https://iupred3.elte.hu/, https://mobidb.bio.unipd.it/ and http://db.phasep.pro/) (*35*). From PONDR, we obtained VSL2- and hydropathy-scores plotted against mean net charge of proteins of interest. The hydropathy-score in PONDR uses a linear discriminate function to define boundaries between disordered and ordered proteins (*112*). We obtained genome-wide phase separation scores based on highly frequent pi-interactions in small amino acids with exposed peptide backbone within IDRs from Vernon et al. (*111*). These pi-contacts were found to be important for protein-protein interaction formation, hydrogen-bond stabilization and solvation, which is directly coupled to the metabolic environment of the cells. Based on these pi-pi contact frequencies in folded proteins, and their correlation with reversible self-aggregation, the authors developed a predictor of pi-mediated phase separation (*111*). We downloaded their proteome wide pi-scores and ranked the proteins with high likelihood of phase separation. As described by the authors, we cross-examined their prediction using other predictors of phase separation. For proteins with high pi-pi phase separation, we assessed whether they displayed more physical interactions and compared it with counterparts displaying the lowest phase separation score. This revealed, top pi-pi phase separated proteins formed more heterogeneous interactions by interrogating their physical partners using PPIs databases (see interacting transcriptional regulators). We next performed functional annotation of those top pi-pi proteins using Enrichr in R and found they displayed metabolic associated functions. Finally, we examined whether PGC1A-interacting network were among top pi-pi based phase separated proteins.

- **Context-dependent phase separation scores**

Physical conformations between intrinsically disordered proteins vary depending on different principles and environments (*59*). Interestingly, solvent-mediated electrostatic repulsions and attractions have been shown to depend on linear sequence confirmation and charge segregation within intrinsically disordered regions (*113*). Then, the proportion of linear sequences with charge segregation in IDRs partitioned structural representations of proteins into categories such as globules, context-dependent globules, coils and chimeras. These conformations form an ensemble of parameters for new phase separation classifiers that complement well established low-complexity regions predictors. We then used classification of intrinsically disordered ensemble regions (CIDER) to obtain parameters related to phase conformational transitions (archived in http://pappulab.wustl.edu/CIDER/about/). We implemented local-CIDER in python 3 (stored at http://pappulab.github.io/localCIDER/), to compute bulk- conformational variations in large data samples. We first focused on globules and context-dependent globule conformation classification (*113*). These were shown to be influenced by environmental variations such as salt, charge, ligand-binding, cis-interactions and solvation (*59*). Among the parameters obtained that are at the basis of their conformation classifier, charge-mediated transitions can be recapitulated by net charge and mixing of opposing residues within IDRs. A list of parameters evaluated by CIDER is as follows: K or Kappa represents segregation of opposing charge residues with 0 being well mixed or intertwined while 1 being separated stretches of positive or negative residues; Fraction of charge residues (FCR); net charge per residues (NCPR); f- negative fraction; f+ positive fraction; hydropathy (*114*) and disorder promoting scores. Fasta files of protein sequences of interest were downloaded from the NCBI repository.

- **Cumulative phase separation scores of f-ITRs**

To quantify phase separation score in the network of interacting transcriptional regulators, we used adipose-tissue and brain tissue-specific network background to interrogate their collective associations. We then discriminated between individual nodes that have been experimentally validated in phase separation and nodes that are computationally predicted to display a high degree of IDRs. We next quantify network z-enrichment with nodes predicted or validated in phase separation (p-value <0.05). To complement this approach, we used data containing a global genome-wide disorder score based on planar pi-pi interactions (*111*). For the selected f-ITRs with observed association in tissue-specific networks we summed up their individual phase separation score to a cumulative phase separation score. We then carried out permutations with random proteins and quantified their cumulative phase separation score. This was followed by Chi-square tests comparison and p-value determination.

- **Experimental validation of heterotypic condensates**

See turbidity assay and live imaging FRAP.

- **Phylogenetic amino acid conservation:**

Phylogenetic analysis was performed on amino acid sequences of proteins of interest such as PGC1A. We used the ConSurf software package to obtain conservation scores and multi-species comparison (*34*). Briefly, it estimates the degree of conservation of amino acid sequences based on iterative comparison with homologous sequences. The conservation rate of a sequence depends on its importance for the structural function. The evolutionary relationship is dictated by the similarity between species comparison in the substitution matrix using previously established Bayesian or maximum likelihood (ML) approaches (*115*). Here below is a brief description of ConSurf stepwise analysis. First, given your amino acid sequence, consurf uses a heuristic algorithm to search default homologous sequences using BLAST across different databases and with known 3D structural features (*116*). A multiple sequence alignment is constructed with these followed by phylogenetic tree building using neighbor-joining algorithms (*115*). Conservation score or evolutionary rate in each residue is imputed using rate4site, and divided in different scales for the degree of preservation. Residues evolving slowly are referred to as conserved and those evolving rapidly are referred to as variable. The degree of preservation depends on different levels of purifying selection, which can be folding constraints, enzymatic activity, ligand-binding or protein interactions. For the rate of evolution, the Bayesian approach takes both into account the stochasticity of evolutionary change and the phylogenetic comparison (*117*). This probabilistic model of amino acid replacement estimates how likely a residue is to influence structure and function of protein domains. 3D structures of proteins are obtained from updated databases such as the protein data bank (*118*). It uses simulation based estimation of amino acid substitutions within the tree branch and compared to known structural dependencies of the protein (*117*). For each residue within the sequence, consurf gives a normalized score of evolutionary conservation from 1 with lowest conservation to 9 with the highest conservation. In addition, consurf mapps the estimated conservation score within proteins, to known 3D models of protein structure (*118*), giving a homology approximation of 3D structural features for the residues. It uses HHPred, which exploits a hidden markov model to look for 3D templates from homologous sequences or similar proteins using PDB (*119*) and extrapolate them to the query sequence with structural features. The features for each residue are the degree of 3D exposure, hiddenness and functionality. The evolutionary phylogenetic tree is calculated using the WASABI platform (*120*) which is integrated with multi-species display of amino acid sequences and the degree of evolutionary conservation based on the aforementioned methods. We then evaluated the phylogenetic tree and the degree of conservation for PGC1A sequence. We obtained for each amino acid a conservation scale, with structural features from known 3D representations. We used this to compare the degree of preservation of intrinsically disordered regions within PGC1A. The IDRs were defined as consensus disordered domains within the protein. We then extracted structural features and the degree of evolutionary rate. To compare this domain with the rest of the protein we first defined following consurf estimations: conserved residues scoring as higher than 7 and law conservation rates as lower than 3. As previously described, from the consensus disordered predictions, we obtained 2 IDR domains in PGC1A.

One close to the N-terminal and the other in proximity to the RNA-binding domains. For both IDRs we quantified the number of residues with low conservation score, high conservation, and average conservation. We then obtained the proportion for each category relative to the total number of residues. We performed the same approach for the rest of the protein and compared the estimation of evolutionary rate between IDRs vs the rest of the amino acid sequences. A similar approach was carried out to assess evolutionary innovation in charged residues within the IDR domains. Final scoring was obtained as the proportion of charged residues (positively charged: arginine[R], lysine[K], and negatively charged: glutamate[E], aspartate[D]) within IDRs with low conservation score, in function of the total number of residues.

- **Combinatorial binding density distribution**

To assess binding combinations of f-ITRs to promoters of HAR-associated genes, we used an integrative ChIP-seq database containing >100000 experiments (archived in https://chip-atlas.org/) (*36*). First, we selected top ranked nodes from the interacting transcriptional regulator network. Next, we parsed and integrated all ChIP-seq experiments for each regulator, binding ±10kb from transcription start site (TSS) of target genes. Given that the peak calls in each experiment are based on MACS2 score as previously described (*36*), we used the average MACS2 score from the available experiments. To filter out all possible combinations of co-regulated target genes, we used a cutoff with an average score >50. Next, we built overlapping maps using venn-diagrams in R from filtered target genes. With this, we quantified the number of co-regulated genes by pairwise combinations of f-ITRs. To assess the frequency and dependencies of pairwise combinations we hot-encoded them and used manifold unsupervised learning to cluster their relationships. We complemented this approach with the use of sieve diagrams or contingency tables as previously described (*121*). With the sieve diagrams, we obtained the dependencies of transcriptional regulators combinations (n=881 and dependency discrimination p-value < 0.001). We next selected top combinations for downstream analysis by overlapping manifold clusters and top-dependencies from sieve diagrams. We then used these frequent pairwise-combinations for f-ITRs to select transcriptional regulators with most dependencies and derive a combinatorial-score for a given gene set. As a background set are all possible combinations with all possible target genes for the selected transcriptional regulators. We then used venn-diagrams in R to quantify the number of genes for each combination (classes are >2, >4, >6, and >8 combinations). The combinatorial score for the background set is then the log fraction between the number of combinations in each combination class and the number of overlapping target genes for selected f-ITRs (comb score = Class # of combinations / Total number of overlapping genes with >2 combinations). We then evaluated the combinatorial score from the binding of the selected f-ITRs to HAR-associated genes as follows: HAR-genes comb-score is the log fraction between the number of combinations in each binding combination class and the overlapping HAR-associated genes (number of HAR-genes with >2 combinations). We compared our target gene set with random sampling gene sets with the same number of genes to our query set. Finally, we performed hypergeometric enrichment to compare our query to the background and random sampling, and defined statistical significance at p-value <0.01. HAR-associated genes bound by several combinations of the selected f-ITRs were functionally annotated by using EnrichR in R followed by selection of common biological terms. We also calculated the PGC1A-network combinatorial score for mediators of polygenic risk for BMI and T2D (see combinatorial disease hubs pipeline).

- **ChIP-MS structuring candidates**

To evaluate regulators that bind active enhancers and promoters and their relationships with PGC1A-network, we used previously published datasets (*61*), and a similar implementation (*62*). Briefly, structuring candidates were identified with chromatin immunoprecipitation followed by mass spectrometry. Antibodies targeting epigenetic tags of active enhancers (H3K27ac) and promoters (H3K4me3) (*122*), were used to precipitate binding regulators. Mass spectrometry was used in those complexes to identify putative candidates. As previously described by Weintraub et al. binding candidates were filtered using the log2 ratio of the IP over the IgG with a cutoff of log2 > 1.

- **PPIs structuring candidates and PGC1A-network**

For the resulting structuring candidates binding both active enhancer and promoters, we performed network propagation with PGC1A-network candidates whose expression was increased after fasting in adipocytes. To this end, we used protein-protein-interaction reference networks (at HuRi) filtered for adipose specific enrichments. Network recomposition (node + 1 interactor) was performed for all direct interactors in both datasets, and network analysis parameters and functional enrichment were obtained as previously described (see ITG for transcriptional relationships).

#### ITG-III: functional relationships within genome architecture

- **Functional embedding of positional gene enrichments (PGE)**

To identify spatial chromatin regions with genes displaying topological dependencies, we used previously established methods of positional gene enrichments (*39*, *119*). Briefly, PGEs exploit topological features of gene locations to extrapolate dosage sensitive active chromatin regions. This approach is based on calculating the hypergeometric distribution along genomic distances for gene sets inputs. For a given region, it corresponds to the probability of having observed genes in that region. To test statistical significance for that region, cumulative p-value distributions are used on random simulations or false discovery rate on large gene-sets. This works as the probability of achieving p-value enrichments that are better than chance estimates and use this to define topological enrichments. The additional constraints to positively categorize a genomic region displaying positional enrichments for a given set of genes are as follows: having at least 2 genes; no smaller region was found by random permutations and cumulative p value distributions; no bigger regions with more target genes were found. This algorithm defines genomic regions by distance of query genes, followed by estimation of p-value distribution to compare chance expectation. This implies that query lists containing genes located in the same chromosome but distantly, might give false positive enrichments. For this reason, we defined target genomic ranges for positional enrichment following structural information related to the approximate size of homotypic highly interacting TADs (*123*). The range for topological regions was then set to 15-mb, where PGEs with larger domains were filtered out. These topological dependencies were defined as enriched regions within the aforementioned span. Given that local homotypic chromatin interactions are more enriched within cytogenetic bands (*123*), we filtered out PGEs without cytogenetic band enrichment, using a curated list of cytogenetic band-gene dependencies (archived in http://www.gsea-msigdb.org/gsea/msigdb/index.jsp). To obtain a global algorithmic performance determined by the number of genes being queried, we additionally performed permutations with random gene sets (same number of inputs), followed by statistical comparisons using Chi-sqare test and p-value discrimination. For every topological gene cluster in the form of PGEs, we derived functional enrichments using Enrichr in R. FDR <0.01 and p-value < 0.05 was used as a threshold to select the significant enrichments for biological terms and phenotypes associated with a given region. PGEs with common biological enrichment across domain-catalogues stored in Enrichr, were categorized as class fPGEs (functional PGEs) for downstream analysis. These hierarchical topological chromatin domains were used as seeds for prioritization of structural chromatin data when indicated, and as inputs for determining long- and short-range structural relationships. Combinatorial-topology PGEs: To identify colocalized enrichments or topological improvements of enrichment between single gene sets and combined gene sets we devised a combinatorial pipeline variation. For example, for cross-tissue hubs of transcription within each biological group, we combined gene sets composing PGEs per group and evaluated a combined-PGE hub over-enrichment. To this end, we used optimal-PGEs, which were defined as PGEs with more than 5 genes (see transcriptional information uncertainty with Shannon entropy and mice phenotype networks). We then compared the proportion of PGEs with >5 / total regions, in combined by group, individual by group and combined random permutations of PGEs with similar parameters for either comparison. To discriminate between the average of expected proportions for individual and random combinations, with the proportions from biological combinations we used chi-square test and p-value estimation.

- **Hi-C Data Analysis**

Representative Hi-C data from diverse cells was procured from (4DN hESC-REF, human lymphoblastoid cells, colon cancer, hs2-hi-c) (*40*, *124*) and uploaded using straw (*125*). To determine A/B compartments in chromosomes, we performed previously established protocols (*126*). To identify TADs, we used previously established protocols (*40*, *127*). Briefly, boundaries were defined at 40-kb, 1-mb genomic regions, and 200-kb window for delta vector calculation. Identification of significant Hi-C contacts at 40-kb resolution was done with a previously established protocol called Fit-Hi-C (*128*). Within a 2-Mb genomic distance p- and q-value were calculated for each bin pair with FDR<1e-5 threshold for peak-calling.

- **Conserved TADs and PGEs embedding**

We used a comprehensive published compendium of 21 human tissues and primary cells Hi-C data downloaded from the GEO database with accession number GSE87112 (*129*), along with previously established protocols described therein (*129*). Briefly, data was normalized using HiCNorm (*130*), vanilla coverage (*124*) or ICE (*131*). For comparative analyses, quantile normalization was used to normalize differences. First line of analysis including compartment A/B and TAD calling were done as previously described (see Hi-C data analysis). TAD boundaries were previously described using established protocols (*129*). Briefly, TAD boundary regions were defined at 40-400-kb, as previously described (*41*). TAD overlaps were defined between samples if 80-kb TAD boundaries were shared, followed by chi-square test to discriminate significance among TAD overlaps. Once conserved segments were obtained, these conserved TADs were plotted as chromosome positions in each chromosome. We next quantified conserved TADs within chromatin domains defined by PGEs as previously described. These chromatin regions were in this case defined by the chromatin regions derived from metabolic-HAR genes or metabo-HAR PGEs. We next calculated the mean of conserved TADs within PGEs and compared it to the mean number of conserved TADs within PGEs from random gene sets. We defined each random permutation with the same number of genes composing the metabo-HAR PGEs. This was followed by chi-square test to evaluate statistical significance and area under the curve of ROC to evaluate the discrimination threshold between comparisons.

- **Frequently interacting regions (FIREs) and PGEs embedding**

FIREs regions have been previously described by Schmitt et al (*129*), and were obtained from the same integrative compendia of human Hi-C data (GEO: GSE87112). Briefly, to identify FIRE bins a poisson regression model (HiCNormCis) was fitted as previously described (*130*), where intrinsic data biases are taken into account as defined by Yaffe et al. (*132*). With this approach the total normalized cis-interactions were obtained, and a gaussian distribution approach was used to estimate the local cis contacts, followed conversion of values to -ln(p-value) denoted as FIRE score and used in our pipelines for downstream analysis. Once conserved segments were obtained, these FIREs were plotted as chromosome positions in each chromosome. We next quantified FIREs within chromatin domains defined by PGEs as previously described. These chromatin regions were in this case defined by the chromatin regions derived from metabolic-HAR genes or metabo-HAR PGEs. We calculated the mean of FIREs within PGEs and compared it to the mean number of FIREs within PGEs from random gene sets. We define each random permutation with the same number of genes defining the metabo-HAR PGEs. This was followed by chi-square test to evaluate statistical significance and area under the curve of ROC to evaluate the discrimination threshold between comparisons.

- **Trans-interacting regions in hESC Hi-C data**

To identify the relationship between metabo-HAR domains and trans-interacting regions in human chromosomes we used data from hESC GEO: GSE35156 (*41*). To assess the trans-interacting interchromosomal regions, we used previously established methods with a few modifications (*42*). Briefly, Hi-C data was normalized with hicpipe, and a probabilistic model was followed to calculate chromatin segment contact maps (*132*). Contact maps are based on quantifying systematically intrinsic biases within the Hi-C data. For contact call significance a binomial distribution p-value estimation was used as previously described (*133*). Bins were defined at 500-kb and contact probabilities were normalized against chromosome length. Trans-interacting networks were then formed with segments as nodes and edges as presence of interaction. An undirected graph was built and used to interpolate metabo-HAR domains. Where a metabo-HAR was present in a given chromatin segment, we labeled that node as either PGE and fPGEs if they had a functional embedding. In addition, we quantified the number of trans-interacting bins within chromatin domains defined by PGEs as previously described. These chromatin regions were defined by the chromatin regions derived from metabolic-HAR genes or metabo-HAR PGEs. We next calculated the mean of the number of trans-interacting bins within PGEs and compared it to the mean number of bins within PGEs from random gene sets. We define each random permutation with the same number of genes defining the metabo-HAR PGEs. This was followed by chi-square test to evaluate statistical significance and area under the curve of ROC to evaluate the discrimination threshold between comparisons.

- **PGE-domains in trans-interacting-network**

Trans-interacting networks were formed with segments as nodes and edges as presence of interaction as previously described (*42*). An undirected graph was built and used to interpolate metabo-HAR domains. Where a metabo-HAR was present or colocalized in a given chromatin segment, we labeled that node as either PGE and fPGEs if they had a functional embedding. We used the NetworkX library in python to perform subsequent topological analysis of the embedded network. We quantified average node degree, mean clustering coefficient, centrality measurements such as betweenness and closeness, information flow parameters such as bridging centrality, and node-prioritization with pagerank and eigenvector centrality. Graph layout such as force atlas, openord, circular, were used for representations when indicated (in figure legends). We performed randomization with similar identified parameters and assessed global structural properties with hESC trans-interacting network. To compare the topological location properties between PGEs and fPGEs, we averaged the clustering coefficients and node degrees for the interpolated chromatin segments or nodes. Again, we compared it to full network averages and random sampling with a similar number of nodes to either PGEs or fPGEs. With the embedded functional annotation obtained before, we identified top hubs and robust nodes with a specific functional cluster. The functional annotation of top clusters were obtained with the functional embedding of PGEs (see above). To filter nodes that both display a specific functional annotation and highly influence information flow through the network, we used bridging coefficient and overlapping functional annotation to hubs and robust segments. Top bridging chromatin segments harbouring fPGEs of interest were selected for downstream nuclear compartment reconstruction. Complementing the network-based prioritization, we functionally annotated genes within top trans-interacting chromatin segments using DAVID functional clustering analysis (https://david.ncifcrf.gov/) (*106*).

- **Integration of short-range chromatin relationships**

To study genomic regulatory element relationships at high resolution, we used previously established methods (*43*). Briefly, GeneHancer map is a genome-wide regulatory elements relationships, comprising around 284000 integrated elements from different databases: ENCODE, the Ensembl regulatory build, the functional annotation of the mammalian genome (FANTOM) project, the VISTA Enhancer Browser, dbsuper super-enhancers, EPDnew species-specific databases of experimentally validated promoters, and UCNEbase ultra-conserved noncoding elements. The approach by the Cohen Lab links regulatory elements to genes, using a combination of relationships: correlation between genes and enhancer RNAs (*134*), enhancer-targeted transcription factor genes, expression quantitative trait loci (eQTLs) (*135*), promoter-capture Hi-C (*136*), and distance-based associations based on immediate adjacency (*43*)(*50*). GeneHancer map of regulatory elements uses a likelihood-based score for each enhancer–gene association. This generates a hierarchical pairing that is defined by the number of sources supporting it (*43*). Given our interest in studying both global and local topological representations and the fact that human-accelerated regions are elements that might not have known links, we included all linking data for downstream analysis.

- **Relationships in short-range linking network**

Cis-interacting short-range chromatin networks were formed as an undirected graph with regulatory regions defined as follows: enhancer elements as distal regulatory regions and genes or promoters as proximal regulatory regions. Regulatory regions were defined as the nodes while the edges were the presence or linking association between them. We then interpolated metabo-HAR domain information as follows: HAR-associated genes described before (see pipeline ITG-I) were labeled in the proximal regulatory regions, while non-coding HAR element coordinates were overlapped in the distal regulatory elements. We next used the NetworkX library in python to layout the graph and perform subsequent topological analysis of the network. We quantified average node degree, mean clustering coefficient, centrality measurements such as betweenness and closeness, information flow parameters such as bridging centrality, and node-prioritization with pagerank and eigenvector centrality. We also performed a modularity based clustering approach to find communities of regulatory-region dependencies. When indicated, the graph layout was hierarchical neighboring modularity. In addition, we performed randomization with similar identified parameters and assessed global structural properties. Using hub-based and influence-based node prioritization, we ranked regulatory regions by their density of interactions. We derived a genome-wide density of interactions for proximal regulatory regions as the global mean degree. We categorized this threshold as a social cutoff, and defined nodes as social if they had 2 times the genome-wide mean degree (>16). Likewise, isolated nodes were defined as nodes displaying < 8 degree interaction. To compare the topological location properties for HAR-associated genes, we calculated a hypergeometric enrichment from the metabo-HAR-genes on the list of social promoters derived from our genome-wide social network ranking approach. We compared this with permutations of PGEs built from random gene sets of equal parameters. This was followed by chi-square tests to evaluate statistical significance. To evaluate distal regulatory dependencies of non-coding active elements, we first classified all regulatory elements by their density of interactions in function to the proportion of enhancers in each group. We then compared the genome-wide behavior of enhancer linking with the linking density of non-coding active elements (see selection from vista enhancer database). We next fitted gaussian curves in the data followed by sum-of-squares F-test to assess variability between the 2 models.

- **Nuclear compartment reconstruction and functional embedding**

Once we evaluated global relationships for the trans chromatin segments of interest harbouring gene clusters from PGE decomposition, we took advantage of the network-based analysis approach to select with high confidence structural representations valuable for the network global dependencies, yet sharing functional annotation. This protocol can be used to study any PGE constructed from any given gene set and it is based on selecting trans-interacting chromatin segments that both display topological qualities and harbour functional-PGEs (fPGE) of similar function (see PGEs in trans-interacting-network). Briefly, top bridging trans-interacting segments harbouring fPGEs that shared similar functional annotation to top hub trans-interacting segments were chosen for nuclear compartment reconstruction and further experimental validation. Given that our combinatorial transcriptional based selection (see ITG-I) and our topological decomposition (see PGE functional embedding) both showed lipid metabolism as a top functional candidate, highly trans-interacting segments (with top bridging scores) harbouring fPGEs for lipid metabolism were chosen for reconstruction. Following any possible interaction in our trans-interacting chromatin network from hESC Hi-C data, we iteratively sought for other segments with high bridging score and with a functional embedding for lipid metabolism. To evaluate if these interactions were present in other Hi-C datasets, we tested their presence in other cell types (see Hi-C data analysis). Because we were looking for likelihood of interactions for downstream experimental validations in cell- and tissue-specific context, we only highlighted their behavior as binary, either present or absent without any other integrative method. They were displayed as shared contacts across studies. To fully reconstruct highly ranked and functionally embedded trans structural relationships, we used the top bridging segment, harbouring metabolic-HAR genes, Chr1:48.5mb to derive other local homotypic interacting segments, and other interacting heterotypic segments with high bridging coefficient and related function, such as Chr1:62mb, Chr2:170mb, Chr2:118mb, Chr4:42mb, Chr4:1mb and Chr11:34mb. Interestingly, we found a high density of interacting homotypic regions around chr1:48, including chr1:43, chr39, displaying top bridging coefficient scores in the network. This highlights this intertwined region of the genome as a modulator of information flow within chromatin architecture. As shown later, this region contains genes related to lipid metabolism (fig. S10), and displays high-density genetic variations related to lipid metabolism as well (fig. S14 and S15). We displayed our selection as Chr1-based sub-compartment triplets with Hi-C derived homotypic and heterotypic interactions, as well as the location of PGEs and fPGEs with similar functional annotation within the composing chromosomes. This subnetwork of compartments was used for downstream genomic-range topological genetics and experimental validation in biological systems of interest.

- **PGE-domains at high resolution in short-range linking subnetworks**

Once we evaluated global relationships for the regulatory elements of interest within the short-range network, we took advantage of a sub-module clustering approach to study at high resolution structural representations, yet keeping its global dependencies. This method can be used to study any PGE constructed from a given gene set. First, we selected functional-PGEs (fPGE) following our topological filtering method (see PGEs in trans-interacting-network and nuclear compartment reconstruction). Briefly, top bridging trans-interacting segments harbouring fPGEs that shared similar functional annotation to top hub trans-interacting segments were chosen for nuclear compartment reconstruction and further experimental validation. For the selected trans-compartments, we labeled the modules where local fPGEs reside, using the short-range linking network, and used these modules for downstream analysis. To embed both network and biological qualitative information in the modules of interests, we used a social network based approach supported by epigenetics and transcriptional binding from experimental data. First, every module was further classified by sub-modules using the module cluster algorithm. This generated territorial clusters of associations for HAR-domains (or any PGE) with intra-submodular and inter-submodular interactions, as well as homotypic inter-modular interactions. We then interrogated iteratively binding enrichments of epigenetic chromatin tags given territorial composing regulatory elements (for each module and submodule). To do this, we used the integrative ChIP-atlas data (see combinatorial binding density distribution), to calculate binding enrichments for enhancer coordinates that form topological clusters of associations. When necessary we used liftover function to match coordinates between different assemblies (USCS genome browser at https://genome.ucsc.edu/cgi-bin/hgLiftOver). To calculate background ChIP-enrichments per module, we used histone modifications (n=16641 experiments) across all cell types (n=61679 experiments), with a threshold of significance of 50 and random permutations for comparison. Data was subsequently presented as fold-enrichment to p-value followed by word-cloud generator showing most frequent histone modifications. We next embedded topological qualitative information from our network-based analysis to classify social and isolated enhancers (see relationships in short-range linking networks). We next differentiated chromatin states between social and isolated enhancers. First, we quantified from the peaks with significant binding enrichment the number of peak differences for all histone marks between both enhancer classes. Given that histone repressive tags govern cell-specific transcriptional identity by blocking unspecific gene programs (*137*), we subsequently compared its binding enrichment for social and isolated enhancers. We quantified the total number of significantly enriched repressive peaks (H3K9me and H3K27me) for social and isolated enhancers in a given module, followed by comparison of the total number of enhancers with the repressive tags. This step allowed us to prioritize social enhancers to further dissect active transcriptional elements and transcriptional regulators for a given module. We then selected coordinates for social enhancers to study binding enrichment analysis using the ChIP-atlas dataset. For this we used TFs and regulators (n=15217 experiments) and similar parameters as indicated before. Binding enrichment was defined as fold-change > 2 and p-value <0.05. Once we established the top regulatory binders to territorial social enhancers (modules or submodules), we assessed the binding enrichment for proximal regulatory regions (±10kb from TSS) for genes within the module or submodule of interest. For a given submodule or module, we then selected top ranked transcriptional regulators binding both social enhancers and promoter regions, and generated a hierarchical list of likely active promoter-enhancers links with their intertwined inter-submodular dependencies. This unbiased approach exploited both integrative and topological associations, which allowed us to generate qualitative information for downstream analysis. In this case, we used for each selected module our previously described pipeline for interrogation of interacting transcriptional regulators (see ITG-II for transcriptional dependencies). Finally, for interacting transcriptional regulators territorially enriched, we highlighted those that recapitulated locally our global transcriptional inferences. This was followed by word-cloud representation (based on number of interactions) and network functional enrichment. We complemented this approach by interrogating local topological behavior of metabo-HAR domains as follows: with a targeted manually curated strategy, we used the ucsc genome browser (archived in https://genome.ucsc.edu/) to find the metabo-HAR domain window containing genes and enhancers of interest. We highlighted active elements and assessed by ChIP-regulatory-binding and TFBS motifs, whether members of the PGC1A-interating network were present among the regulators.

- **Selection of active non-coding elements within PGEs**

To select PGE or fPGE regions that contained experimentally tested activity of regulatory elements, we used the VISTA enhancer resource containing in-vivo validated enhancer information (archived in https://enhancer.lbl.gov/frnt_page_n.shtml) (*50*). For any given PGEs, in this case, PGEs derived from HAR-associated genes (table S3), we evaluated the number of active non-coding elements within the chromatin region. For example, around 84% of metabo-HAR PGE regions displayed active elements. We next assessed the number of active elements for the fPGEs that reside within chromosomes composing nuclear compartments of interest (see nuclear compartment reconstruction). We then labeled these elements for experimental validation in biological systems of interest. Using the short-range linking network, we were able to prioritize genes (promoters or proximal regulatory regions) that were associated with active elements. We next overlapped ncHARs (archived in http://docpollard.org/research/) (*20*), with top ranked active regulatory elements within the network module and submodule of interest. We used these coordinates for downstream experimental validation.

- **Chromatin 3D-reconstruction**

To assess global and local 3D conformations from topological data of chromosomes composing the reconstructed nuclear compartments and within chromatin domains derived from PGEs, we used previously established protocols (*138*)(*139*)(*52*,

*139*) (archived in as follows: GSDB at http://sysbio.rnet.missouri.edu/3dgenome/GSDB/; Spacewalk at http://3dg.io/spacewalk/). Briefly, for global chromosome reconstruction GSDB and spacewalk modelling was used to explore the architecture and homotypic interacting clusters where PGEs reside. GDSB is a comprehensive repository of 3D modeling algorithms and Hi-C data. Spacewalk is an application interface with interacting animation for the navigation of genomic-ranges. Using the GSDB application, we first compare the performance between 2 distance-based algorithms for 3D Hi-C reconstruction. We obtained structure evaluation for representative Hi-C at 40-kb resolution of 3D reconstruction using 3D maximum likelihood algorithm (*138*), and LorDG which uses a nonlinear Lorentzian objective function to optimize realistic interactions (*140*). In addition to the same Hi-C data previously used for isolating trans-interacting segments (hESC HiC; GSE35156), we used colorectal adenocarcinoma cells (GEO: GSE105318), human brain pericytes (GSE105513), astrocytes from the spinal cord (GSE105957), human fibroblast IMR90 (GSE35156), brain microvascular endothelial cells (GSE105544), and human hepatic endothelial sinusoid cells (GSE105988). To compare the structure reconstruction between these distance-based algorithms, we used the average spearman correlation of Expected Distance vs. Reconstructed Distance (*139*) on the chromosomes that composed our selected nuclear compartments and the small chromosome 19 as a control. Next, we represented the data in a XY plot as performance score (expected vs reconstructed distances) vs chromosome size (log). We then evaluated the performance across different cell types by performing a linear regression followed by F-test statistics. This algorithmic selection showed that 3DMax maximum likelihood algorithms represent better in 3D Hi-C data including long chromosomes. Therefore, we next evaluated in spacewalk 3DMax pdb-files to assess the exposure of metabo-HAR PGEs, homotypic interactions, homo-heterotypic relationships, as well as the genomic range of their cluster association.

- **CTCF- and RNAPII-mediated chromatin contact domains**

To complement the global reconstruction of 3C chromatin interactions, we evaluated targeted chromatin contact domains with previously published approach (*51*). Briefly, CTCF-mediated modeling is done from chromatin interaction analysis and paired-end tag sequencing (ChIA-PET) from several cell lines (GEO:GSE72816). Tang et al. employed a hierarchical method to integrate specific structure candidates such as CTCF ChIA-PET data with all chromatin contacts from Hi-C data into higher resolution chromatin folding structures. First, PET inter-ligation products with a genomic span of 8-kb were used for inter-region interactions. These were expanded 500-kb along the genome to calculate the interaction frequencies and to define PET clusters for sequencing as previously described (*51*). These inter-ligation PET cluster products were used to call singletons as enriched interactions between discrete chromatin loci (*141*) (*142*). To identify binding peaks MACS pipeline was used (*143*). For intra-chromosomal normalization data was partitioned into 1-mb bins, where targeted quartile normalization was applied as described by Tang et al. (*51*). CTCF ChIP-seq of ENCODE data and STORM CTCF motif interrogation was also used to find overlapping foci for filtering of CTCF loops. To identify chromatin contact domains (CCDs), data was clustered into loops and gaps, followed by aggregate calculation of CTCF loop coverage along all chromosomes and filtering out of regions with a span <10-kb. CTCF anchors were defined using CTCF binding peaks and motifs as previously described (*51*). To map transcription functionality into CCDs defined by CTCF, RNAPII (RNA-pol II) ChIA-PET was analyzed similarly to the CTCF approach as previously described. RNAPII loops were filtered following GENCODE promoter (TSS), transcription end site (TES) and enhancers overlapped data. Briefly, RNAPII loops were then overlapped with CTCF-CCDs and found an 85% approximate inclusion of RNAPII loop clusters within CTCF domains. Single- and multi-gene loop complexes were defined and identified as previously described (*144*) (*145*). This depends on the amount of promoters involved; 1 for single and more for multiple complexes. In addition, this finding allowed us to define with confidence the rule for node notation in our short-range linking network, where genes were identified as proximal regulatory regions and enhancers as distal regulatory regions. Multi-RNAPII complexes showed promoter regions in heterotypic clusters can behave as distant regulators of transcription within CTCF-domains (*51*). The topological domains are displayed as CTCF-graph clusters with loops as arcs within the genomic range. Because depending on the genomic position CTCF-cluster might vary, statistics on the genomic range of interest are displayed as loci distribution and cluster length distribution. We used these PET-based domains and the superimposed RNAPII-complexes to compare the distribution and boundaries of these CCDs with our short-range linking network (see above). More specifically, we compared the submodule boundaries of the integrative short-range network with CCDs-based boundaries, and found a high degree of overlapped boundaries within our PGE domains. Altogether, the integrative topological comparison by Tang et al. between a targeted CTCF-mediated and untargeted methods for topological domain reconstruction (CCDs and TADs respectively), showed a high degree of superposition between both methods. This underscores the relevance of CTCF cis- and trans chromatin binder regulation while regulating local complexes for transcriptional tuning through multi-RNAPII loops. Given that functional domain of transcription are CCDs and TADs, and that intra-domain stochasticity have been represented in loop-extrusion and polymer-inspired models (*70*), we used submodular hierarchical node ranking from our high-resolution short-range network to interrogate loop anchors with CTCF enhancer binding discrimination (see hierarchical enhancers prioritization).

- **CTCF-mediated 3D-reconstruction**

To complement the global reconstruction using 3C chromatin interaction frequency, we evaluated multi-scale local 3D modeling of PEGs and segments of interests with previously established protocols (*51*, *52*, *146*) (archived in https://3dgnome.cent.uw.edu.pl/). Briefly, the authors used a hierarchical tree based model of ChIA-PET derived CCDs

(ChIA-PET data stored at GEO: GSE72816). The chromosomes are then modeled following a tree-based representation where parent-child node relationships are defined by the resolution. They used a beads-on-a-string polymer model where beads are genomic coordinates of 1-2 mb genomic range with PET-clusters within. Anchors and subanchors were defined by CCDs and gaps between them respectively. The hierarchical clustering algorithm aggregate then neighboring interacting segments following genomic size constraints. This approach allowed it to go from different levels of resolution; for example from low with singleton units or CCDs to high resolution with intra-CCDs ChIA-PET defined loops. For structure reconstruction a monte carlo simulated annealing function is used to minimize the energy in each level. At low level the singleton data is used to minimize simulations while at higher resolution PET-loops are used to optimize local interactions. To translate loop arc data to physical genomic distances a inverse power law approach was used, where the number of interactions in a given PET genomic range reflects physical distances. Finally, 3D models are tuned depending on loop directions and singletons size. With this approach we then obtained CTCF-mediated 3D models of local PEG domains and their relative structural genomic range.

- **CTCF-bound regulatory regions**

To identify gene promoters bound by the structuring protein CTCF we used the ChIP-atlas dataset as previously described (see combinatorial binding density distribution). For the genes identified as CTCF-bound, we assessed whether they were classified as social proximal regulatory regions by obtaining their association degree from our integrative regulatory network (see short-range linking network). This was followed by hypergeometric enrichment of CTCF-bound promoters within promoters defined as social. We used the Chi-square test to evaluate statistical significance. In parallel, we interrogated ChIPseq-enrichment of CTCF in enhancers defined as isolated or social (see short-range linking network), using again the ChIP-atlas dataset. In addition, we used other database compendia of enhancers for interrogating CTCF-bound elements in local chromatin regions (see structural enhancer prioritization).

- **Structural enhancers prioritization**

We next sought to identify structural enhancers within genomic ranges defined by fPGEs (in the present study as metabolic-HAR regions). We used enhancer coordinates within submodule territories of the integrative short-range linking network. To prune our global network approach with enhancer datasets including structuring-based experiments (ChIA-PET and Hi-C) and identify TF binding candidates across several cell lines, we used the HACER (human active enhancer to interpret regulatory variants) database (*147*) (archived in http://bioinfo.vanderbilt.edu/AE/HACER/index.html). In brief, HACER is a comprehensive database that incorporates interaction frequency experiments to the linking association derived from GROseq, PROseq and CAGE profiles. Enhancers in HACER are cross-referenced with VISTA enhancers (*50*), ENCODE elements, the ensembl regulatory build (*148*), and chromHMM (*149*). Enhancer interactions in HACER are defined using similar parameters to the GeneHancer regulatory network used here to perform network propagation of short-range association. These are genome distance, eQTLs, and FANTOM. Besides these, the authors used structural-mediated linking to improve their associations with experimental interactions from 4D genome consortium (*150*) and chromatin contacts derived datasets (*51*, *136*, *145*). Moreover, HACER has integrated ENCODE ChIP-seq data from several cell lines and tissues. This includes > 700 000 TF-enhancer binding and 156 TFs. In order to interrogate TF-binding of structuring candidates as well as experimental structural validations of enhancer-gene associations, we used as inputs the coordinates of regulatory regions within fPGEs. We then used HACER assemblies as test datasets for our global networks, where enhancers bound by CTCF in HACER were classified as social or isolated with the network parameters of the short-range linking networks (see short-range linking network). With this approach we were able to estimate the average number of links that structuring enhancers display within selected chromatin regions. In sum, this was a qualitative step to filter short-range network representations within regions of interest, in submodules for downstream experimental validation. For the filtered enhancers within fPGEs, we used HACER GWAS-integration of regulatory regions displaying SNPs for experimental validations (*151*).

- **Cell-specific regulatory regions**

To identify cell-specific regulatory elements in our global short-range association networks, we used 2 different approaches relying on previously established protocols and datasets. ***Enhancer atlas**:* First, we used the enhancer atlas (archived in http://www.enhanceratlas.org/index.php) (*71*). Briefly, this atlas contains regulatory elements for > 500 tissue and cell types across several species. It incorporates high-throughput experiments including chromatin states, DNAse-seq, ATAC-seq, ChIA-PET, GRO-seq, STARR-seq and MPRA among others. To combine different tracks from each dataset and generate a consensus assembly the authors used an unsupervised approach that weights each track for the global track (*152*). Peak filtering is done at 2500bp and merging with the average summit in the size of the average peak width (ASW) (*153*). To focus on the degree of overlap between regulatory elements, Tianshun et al. used the Jaccard index with intersection over union (*71*). Using this resource we downloaded cell-specific enhancer coordinates and mapped them with our short-range regulatory network, highlighting their location and network statistics. We then used this qualitative filtering step for experimental validation of cell-specific activity during fasting. To this end, we performed ChIP-qPCR with epigenetic tags of enhancer activation (H3K27Ac), and transcriptional regulatory binding. Other tissue-specific modules with motif enrichments were obtained from Epimap (*57*).

- **Structural variants density within PGEs**

Structural variants (SVs) promote large genomic rearrangements that have profound impact on disease and evolution (*54*)(*55*). These rearrangements can affect dosage of protein-coding genes and cis-regulatory function (*55*). Until recently, single-nucleotide variants (SNVs) have been the reference in genome-sequencing population genetics. However, new efforts to generate maps of SVs from diverse populations have highlighted their potential impact on functional genomics. With this in mind, we have integrated into our pipelines approximate 15630 SVs data from the reference resource compendia developed by Collins et al. (*55*) (archived in https://gnomad.broadinstitute.org/). As previously described and for our purposes, we used common SVs (allele frequency >1%) that displayed strong linkage disequilibrium (LD) ($R2 \geq 0.8$) with common short variants with a reported association to GWAS catalogue and UK-biobank (*55*). These selected SVs in LD with GWAS variants were found to be enriched in genic and intergenic regulatory regions. To evaluate the density of SVs within PGEs located in the chromosomes composing nuclear compartments of interest, we partitioned SVs location by cytogenetic band genomic range (resolution 850 bphs). We then displayed the number of SVs per cytogenetic band within the selected nuclear compartment. We next located the band with fPGEs harbouring HARs, as well as other neighboring and interacting bands composing both topological domains and 3D interacting homotypic regions.

- **Genomic region enrichment annotation tool (GREAT)**

To assess how pervasive the functional annotation of HAR genes, and specifically the metabo-HAR regions obtained by our ITG-pipeline, we performed GREAT analysis (*154*), to find ontologies of the genes within the same genomic regions. Briefly, HAR coordinates (table S3) were used as an input into the GREAT software (available at http://great.stanford.edu/public/html/index.php), which evaluates biological process enrichment for neighboring genes compared to genomic background, followed by bonferroni-corrected test to estimate p-values.

- **Genomic-range genetics**

To find metabolic disease associations for a given genomic-range, we used the curated common metabolic disease portal. (archived in https://hugeamp.org/) This integrated database allowed the analysis of human genetics linked to common metabolic diseases. To evaluate genetic association enrichments for fPGEs or genomic regions of interest, we used previously established methods (currently used at hugeamp) (*56*). Briefly, this protocol is based on meta-analysis of associations that provide an estimate of variant effect by integrating information from diverse studies. By taking into account the sample overlap among datasets, this approach weights in the study contribution for the final estimate. The meta-analyses are implemented in METAL which is a tool for genome-wide scans that was described by Willer et al. (*56*). Their approach is based on p-values and subsequent z-statistics for the impact of the association. Given that all studies used are aligned to the same allele, z and p-value are obtained from the weighted sum of individual statistics. The weights are then the square-root of the number of individuals in each sample. Following their streamline implementation of METAL and genome-wide scans we interrogated the cumulative association of genomic-regions to metabolic disease traits. For this, we used fPGEs in chromatin segments composing heterotypic compartments including those in Chr1, Chr2, Chr4, Chr8 and Chr11. Next, we assessed genetic associations to homotopically interacting chromatin segments and displayed p-value of association for a given region. We obtained from this, trait groups and top traits within groups. We displayed the integrative results for all genomic-range traits from metabo-HAR regions as a wordcloud representing the number of appearances of groups and trait descriptions.

- **Epimap integration**

To integrate epigenomic data with high-resolution into our pipelines, we used previously established methods developed recently (archived in http://compbio.mit.edu/epimap/) (*57*). In brief, Epimap includes > 800 biosamples with > 10000 epigenomic maps from Encode and Roadmap epigenome consortium (*155*, *156*), all processed uniformly and integrated following a machine learning approach to impute chromatin states, tissue-specific enhancer modules, upstream and downstream regulators (*57*). This comprehensive compendium of functional enhancer modules with motifs, and GWAS trait enrichments revealed patterns of disease dependencies and comorbidity. We downloaded results from this resource including enhancer modules with tissue-specific enrichments, enhancer-gene links, GWAS enrichments, tissue-trait, and trait-enhancer relationships for downstream network analysis. We next built undirected relational graphs and performed hierarchical neighboring analysis to identify module dependencies. Each module was classified depending on its phenotype composition, and ranked by node-degree. We next used multidimensional scaling and node prioritization to uncover module drivers and information flow bridging nodes (bridging coefficient). We focused on metabolic- and lipid related traits to carry out network propagation and find out the degree of trait permeation within modules and with interacting modules. For top associated enhancers enriched in driver tissues, we extrapolated enhancer-gene information. In addition, we displayed inter-related trait dependencies in function of bridging coefficient score (mean per module) within obtained modules, which showed their lipid-trait hierarchical dependencies for general network-flow bridging score coefficients. For related genes with modular, or bridging dependencies we assessed topological enrichments to identify spatial positional hubs. For each inter-related module, we quantified positional hubs of driver genes, individually (per module), and combinatorially (for top interacting modules; see functional embedding of positional gene enrichments). We then assessed if individual module hubs, and topological top driver hubs, enriched in relational traits obtained here, were colocalized with other interacting metabolic-HAR hubs in general, and those within reconstructed nuclear compartments.

- **Aggregated topological expression from scRNAseq and bulkRNAseq**

For aggregate topological expression we used previously published data from human brain snRNA (*76*) and human APC-bulk expression data (*75*) (see human brain single nuclei RNA seq analysis and transcriptomic profile of human adipose precursor cells). We then partitioned the datasets into fPGEs of interest, in this case our selected metabo-HAR domains within the reconstructed nuclear compartments. For the genes composing a given fPGE or PGE, counts of expression were aggregated per biosample or cell subtype defined by the clustering representation. We next represented local transcriptional activity within the PGE by the aggregate expression number for each gene (y axis) vs an approximated location of corresponding genes (x axis) following the human genome assembly Hg19. Gene location number was defined depending on the appearance following a 5' to 3' direction. To compare topological transcriptional activity between groups we used Area under the curve (AUC). To give an estimate of topological regional expression, we simulated the aggregated counts within each biological group.

#### ITG-IIIa: transcriptional burst models

- **Dynamic gene expression data**

To quantify dynamic gene expression changes in cellular systems of interest, we used intrinsic fluctuations captured by targeted gene expression (see analysis of gene expression) between challenged (i.e. fasting) and control states (RNA extraction and cDNA library preparation protocol was performed as previously described). The intrinsic fluctuations of dynamic gene expression were used to build statistical models using a Monte Carlo approach (see Monte Carlo modeling). Modifications in cell culture for broad capture of gene expression were as follows: cells were plated in 96 well plates in equal cell numbers (1x10e4 cells/well). Induction and differentiation were carried out as previously described (see mammalian cell culture). Once differentiated, fasting (low glucose medium) was induced in half of the wells followed by dynamic time tracking, where every plate was destined to track 1 different time point. After 3 hours of fasting, cells were collected (1 plate per time point) and RNA extraction protocol was performed every 10 minutes for at least 10 different times (T0 + 10 additional measurements; where T0 = 3 hours after fasting induction). We collected at least 6 biological replicates per time point followed by qPCR gene expression analysis as previously described. To compare the expression of genes within the same TADs, CCDs or submodules as previously defined (see ITG for structural chromatin relationships and short-range link networks). For the selected TADs within PGEs of interest, we quantified fold-change distribution across time variations in genes with social or isolated promoters. These were defined by network integration and propagation analysis of short-range link networks. In brief, we obtained fold-change variations (and distributions) by comparing the expression between control and fasting in every time point evaluated. These dynamic fold-change fluctuations between fasting-induced genes were used to build downstream statistical models. We were interested in studying, for differentially expressed genes within the same TADs during fasting, how much their gene expression fluctuates over time comparing it with control bursting fluctuations. We observed that, for genes regulated during fasting, there were time variations in fold-change over control, indicative of specific bursting patterns (**fig. S17N**). Indeed, genes that were defined as isolated by our network approach did not display dramatic fluctuations as those defined as having social promoters (**fig. S17N**). Their fold-change variation over time was stable and constant during fasting (**fig. S17N**). On the other hand, active genes with social promoters recurrently exhibited bursting fluctuations in several experimental setups indicative of different kinetics of expression. To control for primer effect we tested several sets and obtained consistent results. In addition, we observed that perturbing genomic plasticity during fasting, blunted this fluctuation in social genes, while preserving their constant and stable pushed expression.

- **Non-linear sinusoidal models on dynamic gene expression**

In order to capture these variations and model statistical differences we investigated non-linear models. Experimental observations showed an intricate oscillating pattern of expression for some genes within fPGE domains. Their behavior was reminiscent of sinusoidal spectral patterns where fluctuations seem to capture energy discrepancies at different frequencies. This oscillatory dynamic behavior has been observed in complex biological systems where synchronicity of population behavior has been exploited to extrapolate principles for non-stationary variability (*157*). Moreover, gene expression burst kinetics have been shown to differ between enhancer- and promoter-encoding burstings as well as models for condensate-control of transcription (*73*)(*74*)(*74*, *158*). With this in mind, we used non-linear sinusoidal regression (sine-wave with nonzero baseline model in graph pad) to model our experimental observations. We fitted models for each target gene fluctuations to assess their best-fit parameter. Non-linear least squares model was used to fit the regression without weighting the data points (*159*). To discriminate the best-fit parameters between models we used the Akaike's information criteria (AICc), which assumes non-nested inferences. Goodness-of-fit was evaluated by $R^2$ and standard error of the estimate. Parameters obtained from the sine-wave models are as follows: amplitude defines the height of the waves from the baseline; frequency is the number of cycles per time unit; baseline is the Y value where the curve oscillates. We next compared the best-fit parameters from biological experiments with randomly generated values that preserved amplitude distributions. To this end, we generated random values per time point followed by sinusoidal non linear regression, and used AICc to evaluate goodness-of-fit between them. In addition, we also modeled basal expression detection of both house-keeping and unstimulated genes. This showed discrepancies between the models, with amplitude and frequencies displaying the largest variation. Burst profile models of social genes exhibited higher frequencies, amplitude and baseline parameters. We therefore used these quantified parameters to simulate in large scale sinewave behavior constrained by these model variations. We used an in-built simulation option in graph pad where XY table values were selected. Time-series variations were on the X values with similar intervals as our experimental data set (>1000 interval simulations). Y values were generated using sinewave non zero baseline model equations with amplitude, frequency, phaseshift and baseline constants from experimentally derived models. Gaussian random error was used to generate random scatter values. Once the simulations were obtained we performed nonlinear sinewave regression on simulated values to get a better estimation of confidence intervals and estimate best-of-fit and oscillatory parameters. We used these statistical approximations for Monte-Carlo simulation followed by outcome discrimination of the models recreated.

- **Monte Carlo modeling of dynamic gene expression**

To estimate the distribution and range of variation among oscillatory parameters in models from different experimental biological observations, we used a Monte-Carlo approach on large-scale simulated data from our previously described step (see non-linear sinusoidal models) (*160*) (*161*). From the simulated sinusoidal models, we obtained best-fit estimates of models constrained by experimental observations. Using these best-fit estimates, we modeled >1000 paralleled models using Monte-Carlo. We defined sinewave equation parameters for each group followed by nonlinear sinewave regression evaluation. Outcome discrimination on each models was carried out with AICc and standard error of the estimate from experimental groups and random generated models. We next tested distribution discrepancies for oscillatory parameters (amplitude, frequency, and baseline) from Monte-Carlo models. Mean distributions between biological groups from which the models were devised, were used to make statistical comparisons between model fluctuations and estimate p-value for model assumptions.

#### ITG-IV: Topological comparative genomics: human-mice convergence and tuning of nuclear compartmentalization

- **Mice-human chromosomal synteny**

To evaluate genomic conservation regions between humans and mice, positional gene enrichments of mice orthologues metabo-HAR genes were obtained following the same criteria as indicated before (see functional embedding of positional gene enrichments). We quantified the number of optimal-PGEs (> 5 genes) between human and mice topological enrichments and compared them to the number of PGE derived from random permutations. Subsequently, mice PGEs regions of < 15-Mb were selected for downstream analysis. We next evaluated whether metabolic-HAR regions (mPGEs and hPGEs) were located in syntenic domains, using the synteny portal (archived in http://bioinfo.konkuk.ac.kr/synteny_portal/) (*162*). This revealed that 82% of PGEs were in syntenic regions and 18% were located in split regions in mice. Of these, some belonged to our network-based nuclear compartment for lipid metabolism and were experimentally validated. We also displayed the level of genomic conservation in breakpoint regions and in fully syntenic blocks (*41*).

- **Mice phenotype networks**

For a given gene set of interest (in this case, metabolic-HAR genes), we identify gene and phenotype relationships from data stored in Jackson laboratory containing mammalian gene perturbation experiments. This dataset has been parsed and is stored in ENRICHR as mammalian perturbation catalogue (*103*). Network reconstruction was done with all phenotypes associated with gene knockout information followed by network integration and analysis. To identify phenotypic modules we used modularity and to identify gene drivers per module we used node degree, pagerank and eigencentrality. To identify phenotype drivers we used node prioritization followed by quantification of phenotype representations by proportion frequencies (top 30 phenotypes) Intermodule communication was performed by quantifying interactions between modules. Positional gene enrichment coalescence of gene drivers within top interacting modules was performed using positional gene enrichments for individual genes within modules and combined genes from top interacting modules. Proportion of positional enrichments by coalescence was defined by comparing positional enrichments with > 5 genes per region (see functional embedding of positional gene enrichments).

- **Transcriptional hubs (PGEs) in chow-diet and High-fat-diet across 19 organs**

We performed positional gene enrichments of differentially expressed genes derived from RNAseq in mice models of obesity (see animal experiment; HFD intervention). In brief, mice either on chow-diet or HFD were sacrificed and 19 tissues were collected for RNAseq profiling. Multi-tissue differentially expressed genes were classified as HFD regulated or Chow-diet regulated. These gene sets were used as inputs for hub PGE-analysis. We displayed PGE hubs as the proportion of total PGE by dividing the number of regulated genes in topological enrichment with the total number of regulated genes. We carried out this approach in every tissue per biological group, and compared them with hub-PGEs formed from random sets (same distribution parameters as the number of genes composing PGEs of interest). To identify overlapping or cross-tissue hubs of transcription within each group, we combined gene sets per group and evaluated spatial hubs. We assessed optimal-PGEs, which were defined as PGEs with more than 5 genes in genomic cluster (see transcriptional information uncertainty with Shannon entropy and mice phenotype networks). We next compared the proportion of PGEs with >5 genes / total regions, in combinatorial hubs by group, individual and control random permutations (with similar parameters for comparison). To discriminate between the average of expected proportions for individual and random combinations, with the proportions from biological combinations we used chi-square test and p-value estimation. We also compared optimal-PGE hubs from genes that were only regulated by HFD or chow-diet (see transcriptional information uncertainty by Shannon entropy). To this end, we used proportions of total topological enrichment to get an estimate of full transcriptional burden by treatment. Proportion of total topological enrichment was defined as the number of genes in PGEs / total number of regulated genes. We plotted representative optimal-PGEs in each chromosome by group (genomic range defined by start and end of chromosomal PGE region) and defined colocalized regions of activity. Metabolic-HAR domains and experimentally validated HAR-regions in human cellular models were highlighted. In addition, we calculated the proportion of unique PGEs in each chromosome, by quantifying the total number of PGEs irrespective of treatment and the ratio contribution per state. To show in full-display this topological coalescence across similar tissues and by physiological state, we plotted all PGEs derived from adipose tissues (subcutaneous, visceral, mesenteric, retroperitoneal and brown fat) from chow-diet and HFD. This revealed both territorial cross tissue transcriptional activity and tissue-specific domains. For those chromatin regions showing convergence, we hierarchically ranked them by p-value of enrichment. Top hubs were indeed cross-fat-tissue convergent regions where transcriptional activity was present. Inter-physiological state comparison demonstrated a HFD-mediated topological shift as a large portion of PGEs were only present in the obesity model. The shift of regions between treatments was defined by a genomic-range transition of at least 500kb for those optimal-PGEs. This yielded evidence about both a global topological shift on spatial regions of transcriptional activity, and information about the spatial transcriptional burden derived from suboptimal PGEs in HFD (see transcriptional information uncertainty by Shannon entropy).

- **Topological PGE-mediated single-cell decomposition**

To decompose topologically bulk-mediated hubs into single-cell PGEs in tissues of interest, we first downloaded and re-analysed single cell RNAseq data from the tabula muris consortium (see adipose tissue scRNAseq from mice). After obtaining differential expressed genes in adipose tissue, we performed PGEs per cell type (see functional embedding of positional gene enrichments). We observed cell-specific transcriptionally active regions as scPGEs from the scRNAseq data with some large PGEs with shared cell pair enrichments (e.g. mesenchymal stem cells and endothelial; B-cells and endothelial). Given that this dataset was generated in wild-type non-treated mice, we assessed whether scPGEs colocalize with hubs derived from chow-diet transcriptomic data. The overlap genomic-range was defined as > 200 kb between PGEs and we quantified the proportion of domains shared as transcriptionally active regions. To assess if the degree of overlap was higher than random expectation we performed permutations with PGEs formed with random gene sets. We showed that random overlap lingered between 20 to 35% while scPGEs displayed > 50% overlap for most cell-specific PGEs in bulk-derived chromatin domains. NK overlap was similar to random expectation while T cell overlap was around 40%. A similar approach was performed with PGEs from HFD in adipose tissue. We overlapped them with fat scPGEs and quantified the proportion of overlapping for each cell type with the HFD-mediated chromatin domains. This revealed that mesenchymal stem cells and endothelial cells scPGEs shared the highest proportion of HFD-regulated hotspots. We then assessed chow-diet specific hubs in fat that are lost after HFD and are located contiguous to syntenic metabolic-HAR domains. These genomic-regions of interest were interrogated for homotypic and heterotypic interaction using contact frequency Hi-C data (see Hi-C data analysis). We then mapped homotypic and heterotypic interactions from PGE-PGE chow or HFD locations that displayed cell-specific transcriptional activity by scPGEs. Metabolic-HAR regions were also mapped in contact frequency maps of interest. Those cell-specific topological variations induced by diet were selected for downstream experimental validation by ChIP-loop, expression and chromatin immunoprecipitation.

- **Transcriptional information uncertainty by Shannon entropy**

In information theory, entropy is defined as the uncertainty of a random variable (*163*, *164*). In other words, it quantifies the expected values of information in a given message, or how much information or surprise levels are related to an outcome. If we take event probabilities, information theory assumes that the lower the probability of an event the higher the uncertainty in the observable outcome. Now, if we have a variable taking many numbers (or events) with a finite quantifiable limit, we can then consider the distribution of its probabilities. Any time we observe an event of probability we gain information, which we quantified as bits. Thus, for whole probability distributions we can assess the average level of uncertainty for given probability of events. To extrapolate information from spatial hubs of transcriptional activity, our assumption based on our observations and previously described observations on transcriptional assemblies, was that localized genomic domains of transcription in the form of positional gene enrichments or hus (derived from high-throughput transcriptional profiles) can be used to estimate the level of randomness or uncertainty in transcription. Moreover, biophysical condensate formation decreases cellular entropy by localizing hubs of cellular activity (*7*). As shown before (see functional embedding of positional gene enrichments), compared to random gene sets, gene sets derived from high-throughput biological observations always displayed an elevated proportion of topological enrichments with > 5 genes within genomic clusters (defined before as optimal hubs). This approach can therefore be considered as an indirect measure of hub-models of transcriptional activity, where the number of observations (number of PGEs with > 5 genes / total number of genes in positional enrichments) for a given gene set defines here “event probabilities” of the variable transcription. We considered the total number of genes in positional enrichments as the full transcriptional output for a given state, which we used to obtain probabilities of localized transcriptional organization to subsequently derive Shannon entropy. The reason to use the total number of genes in positional enrichments rather than the total number of PGEs for a given state lies in our previous observations in Fig 4. These showed that, for a given state across many organs (chow-diet or high-fat diet), around >60% of genes are within PGE hubs. Of these, around 15% are located within top hub PGEs of >5 genes. This yielded evidence that the largest discrepancy in transcriptional burden lies on the number of genes being transcribed and located within suboptimal <5-genes PGEs. In that regard, qualitative hub-PGEs of >5 genes were defined by comparing the average p-values between optimal and suboptimal PGEs using all transcriptomic data (independent of state) vs random permutations. In addition, we plotted fold-enrichment vs p-value followed by non-linear regression and f-test statistics to compare random and biological models. We then perturbed optimal PGEs to show that random and biological models of suboptimal PGEs are similar. Therefore, for every organ or biological dataset we derived the “event probabilities” as the number of optimal PGEs with > 5 genes / total number of genes in positional enrichments. We used these proportions as observed event probabilities to calculate individual (per sample or organ) or global (event probabilities from organs in a given state) Shannon entropy (**Fig 4G**). This was performed using python library for scientific computing scipy. Finally, to show we can capture random transcription by single genes using PGEs in a given state, we first partitioned unidirectional genes by state. This was done by taking all differentially expressed genes across all 19 tissues, followed by venn-diagram intersections to define genes that were ‘only’ regulated by HFD or Chow-diet. These unidirectional gene sets were used to quantify PGEs per state and the proportion of total topological enrichments (**Fig 4H**). This showed HFD increased the transcriptional burden, similar to random-derived positional enrichments (**Fig 4H**).

#### ITG-V: Combinatorial disease hubs: polygenic risk scores (PRS) and causal mediators

- **Introduction**

Metabolic-related traits and other complex-traits are polygenic with >90% of variations located in non-coding regulatory regions. They also display pleiotropy as the same variant associations are shared between traits (*165*). Organismal and cellular metabolic states influence the genetic burden of other complex-traits, which highlights the ubiquitous and pervasive nature of metabolism on disease risk. Indeed, obesity and type-2 diabetes increase the risk of developing a plethora of pathological processes determined by individual genetics (*165*). It then becomes paramount to understand the intertwined relationships of shared variations among complex-traits with specific focus on metabolic disease maps. The systematic dissection of cis-regulatory-gene associations is a key step to decompose hierarchically metabolic disease and subsequently its multi-trait penetrance. To this end, we used results from previously published approaches where causal inference methods were developed to systematically assess the dependencies between genetic variations, genes, tissues and traits (*79*). Park et al. implemented an inference framework based on causal multivariate mediation analysis with extended linkage disequilibrium (CaMMEL) (*79*), to identified causal cis-regulatory genes mediating the effects of complex traits. This approach uses eQTLs in 49 tissues from Genotype-Tissue Expression (GTEx project) and jointly analysed them using summary statistics to define tissue-specific causal genes. Source code for CaMMEL is publicly available as a part of the R package for summary-based QTL / GWAS analysis (available at https://github.com/YPARK/zqtl; https://github.com/YPARK/cammel-gwas).

- **Data preparation, PRS and TWAS statistics**

Here below, we give a general description of the method for multi-tissue PRS causal mediation developed by Park et al. with specific implementations in metabolic disease (BMI and T2D) by integrative topological genomics pipelines used in this study. For **data preparation,** Genomic locations of GTEx eQTL summary statistics and genotype matrix were used, linkage disequilibrium blocks and dosage matrix of GTEx data were done using published methods (*166*, *167*). Additional variations on LD estimation can be found at Park et al. (*79*). For **Polygenic risk scores (PRS) and TWAS statistics**, a summary-based method (*168*) was used to predict PRS from GTEx individuals. Specific information and full results can be found at https://github.com/YPARK/PRS. After having the PRS of gene expression and GWAS, a linear regression analysis of the GWAS PRS on the expression PRS identifies slope parameters and standard error. These are then used to estimate the shift from zero by building the Wald test statistic. This corresponds to summary-based TWAS statistics as previously described (*169*, *170*).

- **Causal Mediation analysis**

To estimate causal loci across different complex traits by their indirect association through inferred intermediate molecular traits of gene expression and enhancer activity, CaMMEL was developed (*79*). It follows a Bayesian generalization of Mendelian Randomization (*171*, *172*) and TWAS (*169*, *170*) approach that uses eQTLs to infer the genetic component of intermediate phenotypes and correlate them with different phenotypes. To define causal mediation problem the authors assume that only cis-regulatory mechanisms drive the effects of mediation between genetic variation and phenotypes. Then the relationships between genotype, mediator and phenotype variables are generated by stepwise linear combination of genotype and mediator information with coefficients. In brief, CaMMEL focuses on finding non-causal effects or confounders, to underscore residual variation as causal genes. To correct for non-causal correlations, the algorithm uses inherent and invariant relationships related to LD blocks, and contrast the confounders with true cis-regulatory effects. Similar control-based approaches have been described before (*173*, *174*). The projection of eQTL and GWAS z-score matrices onto independent LD blocks constitutes the control matrix. Confounding factors on mediators and phenotypes are removed by matrix factorization, followed by removing their effects on GWAS traits and estimation of residual effects. This is followed by summary-based multivariate regression of corrected GWAS z-scores on the cis-eQTLs z-scores for all genes and tissues (*79*). Full method details of CaMMEL implementation and benchmarking against TWAS methods with the systematic analysis of GWAS traits and cis-eQTLs from 49 tissues (GTEx datasets), can be found as follows:

  - Source code can be found at https://github.com/YPARK/GTEx_mediation
  - Results at https://github.com/YPARK/GTEx_mediation/tree/master/docs/share/20190701

- **Multi-tissue PRS and mediators of complex metabolic traits**

Systems genetics implementation of CaMMEL by Park et al. revealed a wide range of dependency combinations that exploit the conjoint dissection of multi-resolution data through one statistical framework. This stepwise prioritization revealed genotype--eQTL-gene--phenotype associations between GWAS traits, 49 GTEx and 17608 protein coding genes. CaMMEL identification of causal relationships reveals the pleiotropic and the pervasive nature of dependencies in complex traits. To study multi-trait, multi-tissue and multi-gene dependencies, and their global and local associations with complex metabolic traits, we performed 3 complementary approaches: network integration, hierarchical prioritization of topological based enrichments (cumulative-PGEs), and hierarchical prioritization of topological enrichments with integration of chromatin structural data.

- **Hierarchical prioritization of BMI and T2D PRS-mediation topological enrichments**

To investigate topological hubs representing territorial models of transcriptional activity shared across PRS-mediation tissues from metabolic traits, we focused on GIANT-BMI and T2D-BMI-adjusted. We downloaded GTEx mediation data for these traits across multiple tissues and filtered genes with a log odds ratio cutoff of 2 per trait-tissue and a TWAS adjusted z-score > 1. Of all the mediators in each trait within the afore-mentioned constraints, we quantified the proportion of unique genes in tissues defined as metabolic and the rest of the tissues. We compared the tissue contribution per trait (BMI and T2D) for the same metabolic tissues. We then performed positional gene enrichment of genesets derived from PRS-mediation. Mult-tissue PRS-mediation genes were classified as BMI or T2D causal mediators. These gene sets were used as inputs for PGE-analysis. We then displayed PGE hubs as the proportion of total PGE by dividing the number of regulated genes in topological enrichment by the total number of regulated genes. We carried out this approach in every tissue per trait, and compared them with PGEs formed from random sets (same distribution of number of genes building PGEs of interest). To identify overlapping or cross-tissue hubs within each trait, we combined gene sets per group and evaluated combinatoria PGE-hubs (see functional embedding of positional gene enrichments). We then assessed optimal-PGEs, which were defined as PGEs with more than 5 genes (see transcriptional information uncertainty with Shannon entropy and mice phenotype networks). We compared the proportion of PGEs with >5 genes / total regions, in combined sets by trait, individual by tissue and combined random permutations. To discriminate between the average of expected proportions for individual and random combinations, with the proportions from biological combinations we used chi-square test and p-value estimation. We also estimated the dependencies of the number of genes in optimal-PGEs in multi-tissue combinations versus random permutations, by assessing the linear regression with p-value enrichments between traits and random expectation. We plotted representative optimal-PGEs in each chromosome by group (genomic cluster range defined by start and end of chromosomal PGE hub region) and defined colocalized regions of activity. In addition, we calculated the proportion of unique PGEs in each chromosome, by quantifying the total number of PGEs irrespective of trait and the ratio contribution per PRS-mediation. Metabolic-HAR domains and experimentally validated HAR-regions in human cellular models were highlighted, and used for downstream analysis.

- **Structural prioritization of topological enrichments within trans-interacting chromatin regions**

Transcriptional factories of co-regulated genes are usually located in or in close proximity to homotypic and heterotypic interacting chromosome regions (*44*, *85*). These models suggest that interacting chromatin regions serve as anchors that increase the likelihood of trans- co-regulation (*85*). This has been further supported by observations where non-random spatial proximity of adjacent chromosomes determine their DNA translocation (*175*). In addition, contiguous regions or regions within interacting chromatin hubs were found to be regulated by transcriptional factories or hotspots (*85*)(*86*). These interactions can, in a tissue- or cell-specific way, be more active from a wide range of conserved global interacting locations to define cell-specific factories (*86*). These events indicate a high-degree of plasticity, likely depending on cell-specific heterotypic transcriptional regulators (*85*). This is supported by our observations on spatially defined positional gene enrichments from high-throughput profiling (Fig 4 and Fig 6), and by the degree of higher-order conservation of structural data including TAD boundaries, frequently interacting chromatin regions (FIREs) (*129*) and trans-interacting regions (*86*, *126*). These observations also suggest a high degree of conservation of trans-interacting domains across diverse tissues and cells. The dosage and usage of the conserved trans-architecture likely depend on cell-specific transcriptionally active regions conditioning the hierarchical combination of interactions, and perhaps the nonrandom location of nuclear speckles within transcriptional factories (*176*). Supporting this, transcriptional factories have been found to be located in proximity to trans-domains, perhaps as they allow less space constraints compared to intrinsic TAD-mediated forces of homotypic chromosomal folding (*85*) (*44*). Indeed, genome-wide crosslinked methods simultaneously mapping higher-order interacting regions have revealed that physical limitations within the nucleus constrained these domains within rather large interacting hub regions (*87*)(*5*). In order to build a map of all possible combinations, we used a naive approach to investigate trans-interacting chromatin regions. This global trans-network was exploited to generate hierarchical maps of tissue and cell-specific territorial activity suited for experimental validation. Then, to investigate the structural dependencies of transcriptional hub models (using for example the PRS-mediation cross-tissue cumulative location), we defined higher-order trans-interacting regions using Hi-C data from representative cell types, hESC (see Hi-C data analysis and trans-interacting regions in hESC), and data from SPRITE (split-pool recognition of interactions by tag extension) a method based on tagging crosslinked protein-DNA-RNA complexes followed by genome-wide detection of higher-order interaction hubs in the nucleus (*87*). We then used a two tier approach in which diversity of both data and algorithmic analysis is exploited to define large trans-domains. This is followed by manual curation across representative cells with a likelihood-based integration within curated genomic windows, as previously shown (*177*). We then tuned and cross-validated these recovered interactions using discovered hubs by SPRITE. First, to define Hi-C based domains we used previously described procedures with hESC (see trans-interacting regions in hESC). We also implemented the approach developed by Belyaeva et al. (*86*) using a large average submatrix algorithm (LAS) with a binary classifier to identify trans-domains in human lung fibroblasts (IMR90) (available at https://github.com/anastasiyabel/functional_chromosome_interactions). For those interacting domain peaks, we calculated in each chromosome the genomic-range average from the genomic ranges identified by both algorithmic implementations. Following these results, we then partitioned chromosome regions by 5-Mb genomic-windows and proceeded to manually curate the presence of trans-domains using HiC data from other representative cell types (see HiC data analysis). For each chromosome, we labelled genomic windows as interacting if they display trans-domain bins from the HiC algorithmic comparison analysis. We then proceeded to the manual curation of heterotypic interaction on analysed HiC data as follows. For each biological cell type, we used a naive binary approach in which positive genomic-windows with trans-domains were labeled as 1 in each chromosome if heterotypic interaction was found with every other chromosome (absence was labeled as 0). We then calculated the likelihood of trans-domain presence in each genomic-window by dividing the sum of interactions with the total number of possible interactions. Taking into account the results from each representative cell type, we obtained a global likelihood score of trans-domain presence by averaging the likelihood scores from the other biological cell types. This final score was used to cross-validate genomic-windows with analysed interacting hubs from SPRITE datasets and with the first step of trans-domain HiC interrogation. This cross-validation was used to tune and filter the nodes that overlapped with trans-domain regions discovered by SPRITE-hubs and algorithmic trans-domains. In brief, using the genomic-range windows as nodes and the presence of interaction as edges we built an integrative trans-domain network. As previously described, we performed network-based prioritization of influencer and critical connector nodes. We used this network as all possible combinations of global genome trans-architecture for downstream modeling of overlapping disease hotspots, metabolic-HAR regions, PGC1A-network regulatory regions, and cell-specific transcriptional activity. We focused on trans-interacting domains with overlapping disease hubs from the PRS-mediation approach for metabolic traits. We then filtered a more stringent trans-interacting network with only metabolic-disease hubs. These hubs were composed of shared BMI-T2D PRS-mediation PGEs locations. This graph representation of trans-architectural metabolic disease was used for downstream analysis. To assess the degree of overlapping from other biologically derived gene sets versus random expectation (randomly generated gene sets), we used a previously described approach. In brief, overlapping between PGEs locations (from different datasets) onto our trans-interacting network is quantified and compared with PGEs location generated from random gene sets. Overlapping was defined as PGEs within trans-regions or with at least 50% overlap. The mean value of the number of regions overlapped by random PGEs composes the random expectation number. This might differ in between experiments as every biologically derived gene set is composed of different gene numbers. For this reason, each comparison stands alone with their own random permutations from gene sets forming a similar number of PGEs. For each comparison made, we calculated chi-square tests followed by p-value assessment. To assess the interpolation of loci positions from GWAS studies with our disease-hub trans-domain network, we used GWAS data for metabolic traits stored and analysed in https://t2d.hugeamp.org/. There we obtained GWAS association for BMI from GIANT UK biobank study (*178*) and for T2D BMI-adjusted from DIAMANTE study (*179*). We then assessed the overlap between GWAS locus with our metabolic disease hubs trans-domain network. The shortest distance path between superimposed loci intra-trait and inter-trait was estimated in our network, and topological distances were compared for dependencies.

- **Topological single-cell decomposition of structural metabolic disease hubs**

To decomposed topologically metabolic disease hubs (PGEs) overlapping trans-interacting domain network (see previous section) into fat-specific disease hubs and fat-specific single-cell PGEs from human obese subjects, we first downloaded and re-analysed published single cell RNAseq data from obese subjects (*90*) (see adipose tissue scRNAseq from obese human subjects). We then obtained differential expressed genes in cell-specific clusters, followed by PGEs assessments per cell type (see functional embedding of positional gene enrichments). In parallel, we obtained PRS-mediation results specific for adipose tissue (see PRS-mediation approach) and compared their spatial location with cumulative multi-tissue mediation hubs. From this, we assessed the overlap enrichment of fat-specific disease hubs within the trans-interacting domain network (see previous section). Our previous observations showed metabolic-HAR domains are enriched across chromatin interacting regions, PGE-hubs act as territorial domains of transcriptional activity, and heterotypic condensates are tuned dynamically to assemble nuclear compartments that exploit genome architecture and function. This is in line with observations on transcriptional factories assembled in proximity of interacting hubs and tuned cell-specifically with the dynamic activity of heterotypic transcriptional regulators (*44*). In the previous section, we argue that global trans-architecture could be used to infer the proximity of transcriptional factories and the cell-specific location of interacting hubs with transcriptionally active territories (see structural prioritization of topological enrichments within trans-interacting chromatin regions). Considering this, we obtained scPGEs from differentially expressed genes (see single cell analysis) and evaluated cell-specific domains in the trans-interacting disease hub network. We defined PGE overlap within each domain as previously described and highlighted cell-specific trans-interacting regions with transcriptional activity. This revealed cell-specific domains, domains with double cell-specificity, and promiscuous domains. Hierarchies of active trans-interacting domains per cell and cell-related were obtained by extracting overlapping scPGE and trans-domains. We used this decomposed network to filter regions for experimental validation of cell-specific chromatin contacts by ChiP-loop and transcriptional activity by gene expression. We evaluated APC-specific contacts, transcriptional activity and their interaction with metabolic-HAR domains mediating fasting endurance (ATG4C- and DGAT-domains). We compared the specificity and regulation of these transcriptional active interactions with neighboring and interacting regions during fasting and with metabolic overload. These revealed APC-ectopic inflammation-driven regulation of trans-regions (enriched for macrophage type 1 and monocytes) during fasting with metabolic overload.

#### ITG-VI: Functional impact of radial chromatin organization

- **Integration of GPSeq data for radial chromatin organization**

Genome architecture determines proper function of genome plasticity. Chromosomes are indeed non-randomly positioned in the nucleus with specific territorial domains with respect to the nuclear lamina (*180*)(*80*). Several studies have revealed structural associations of radial chromatin organization with gene density (*180*), while genome-wide approaches have provided high resolution maps (100kb) of genome locations along the radial axis (*80*). However, functional interpretation of radiality with ensemble multi-resolution models aimed at understanding complex disease is still missing. Therefore, we have integrated previously published data derived from genome loci positioning by sequencing (GPSeq) method, which gives relative distances between genomic-loci and the nuclear lamina (*80*) (GEO:GSE135882; GPSeq protocol available at https://doi.org/10.21203/rs.3.pex-570/v1; data available at https://github.com/ggirelli/GPSeq-source-data). Briefly, radially directed enzymatic restriction of fragmented genomic DNA is paired with sequencing. To follow enzymatic radial diffusion with microscopy, Girelli et al. used a modified in situ hybridization in which adapters ligated to the restricted fragments are detected by fluorescently labeled oligos. The adapters guided next-generation sequencing, library generation and radial estimates were inferred by comparing radial locations with 3D DNA FISH methods (*181*). For genome-wide GPSeq score calculation the authors employed custom pipelines. In brief, after quality control of the sequencing data, the genome was binned in 1-Mb overlapping windows in steps of 100-kb, followed by calculation of digestion probability considering all restriction sites cut. GPSeq score from different experiments was generated by averaging the score in each window followed by log2 scaling (available at https://github.com/ggirelli/gpseqc). We then obtained genomic-loci radial scores at 100-kb resolution, which was used to superimpose previously generated PGEs. First, we divided each genomic position in ranges of radial probabilities by peripheral, central and core genome-bins. This was carried out by estimating the average GPSeq score of core-located chromosome 19, and peripherally located chromosome 4. *GPSeq-score in genomic-positions used in this study*: We plotted these 3 categories and assessed the overlap with PGEs derived from either BMI or T2D polygenic-risk score mediation hubs (see PRS-mediation PGEs for metabolic traits). To quantify the PGEs classification, we considered the average of the GPSeq-score for the given genomic-range. To assess the radial proportion, we took into account all PGEs within each group and estimated the proportion of radial location following the range of radial probabilities. We also approximate random expectation likelihood of radial enrichments by assessing the radial proportions for PGEs derived from random gene sets (30% peripheral and 15% core). Similarly, we estimated the radial proportion location of active enhancers by VISTA, metabolic-HAR domains, frequently interacting regions (FIREs), and parameters derived from our network-based analysis of interactions including social and isolated regulatory regions.

- **TFBS-radial distribution and transcriptional PPI networks**

We used GPSeq-mediated transcription factor binding sites (TFBS) radial enrichments as previously described (*80*). In brief, predicted TFBS were obtained as previously described (*182*) and their appearances quantified within the genomic windows. GC-content in each TFBS was quantified. Correlation coefficient between the number of TFBS and GPSeq-score for each window computed against the GC TFBS content. For downstream analysis, we used transcriptional regulators with TFBS-radial peripheral or core correlation. We next assessed in each group their propensity to form heterogeneous physical interactions. To this end, we used protein-protein interaction data from the HuRi repository (see ITGs for transcriptional dependencies) (*104*), followed by network analysis of interaction parameters. We quantified the proportion of interacting transcriptional regulators in each group by dividing the number of interacting factors by the total of regulators per group. Given the differences in interaction probabilities, and our previous observations on co-activators being hub-assemblers of transcription, we quantified the number of co-activators and their interacting transcription factors within each radially enriched group. This showed co-activator-related radial enrichment in core factors. For each of the networks formed, either core-TR or peripheral-TR network, we assessed their functional embedding with reactome and gene ontology network enrichments (p-value < 0.01). This revealed interacting heterogeneity of radially enriched regulators and divergence of functional annotation.

- **Context-dependent phase separation score for radially enriched regulators**

We downloaded FASTA amino acid sequences from transcriptional regulators with radial enrichments. We then used them as input to localCIDER and obtained the ensemble parameters of phase separation and its conformational dependencies. We used the globule conformation classifiers to quantify the proportion of context-dependent phase separation among core or peripheral transcriptional regulators. We next compared both groups using the average value in all parameters obtained, including segregation of charge residues (kappa) and charge-related variations (f-, f+, NCPR, FCR, hydropathy). We built a correlation matrix to study their associations within each group.

- **Radiality of transcriptionally active loci in single cells by DNA-MERFISH**

To evaluate radial positions of transcriptional active loci in single cells and their structural relationships, we used datasets from established genome-scale approaches that use multiplexed FISH imaging technology (*77*) (available at https://zenodo.org/record/3928890#.YDgeiZNKiuV). Briefly, this approach is based on systematic integration of sequential rounds of genomic-loci imaging using in situ hybridization (*183*) at single cell resolution (*77*). This is coupled to a combinatorial strategy previously described as multiplexed error-robust FISH (MERFISH) where imaging combination rounds determine chromatin loci identities (*184*). They coined DNA-MERFISH technique to specific modifications on combinatorial rounds following observations on chromosome territorial location and polymer models. It assigns to each genomic loci a 100-bit barcode, which determines the signal across imaging rounds. To detect the barcodes imprinted in genomic loci the authors used sequential hybridization of labeled probes. With this method, a comprehensive resource of nuclear architecture was generated. This displays single cell resolution of more than 1000 genomic loci information containing both nuclear structural relationships and nascent RNA transcriptional burst activity (*77*). This systematic method images each 30-kb genomic window size for each chromosome, in IMR-90 cells. Multi-resolution Information is then available from around 1000 genomic loci in > 5000 cells, including nascent transcripts from genes located within these loci and their distances to nuclear structures (e.g. nucleoli, nuclear speckles and nuclear lamina). Data processing and script functions can be found at https://github.com/ZhuangLab/Chromatin_Analysis_2020_cell. We used genome-scale 3D position of the measured genomic loci to assess distance to lamina of specific metabolic-HAR PGEs, and other domains composing reconstructed nuclear compartment for lipid metabolism. In addition, we used genome-scale transcription of genes contiguous to the loci imaged by DNA-MERFISH. This extended multimodal approach allows the quantification of transcriptional burst activity and their structural relationships. We used this dataset to obtain transcriptional bust profiles of genes with radial enrichments. We used the structural parameter “distance to nuclear lamina” as an estimate of radiality, and partitioned genomic loci by transcriptional burst on or off. For those genomic-loci with different structural associations (nucleoli, nuclear speckles and lamina), we quantified their mean distance to lamina when they are transcriptionally active on or silent off. We compared these mean distances and estimated their p-value. We next evaluated those genomic-loci with genes with bursting activity on, the correlation between their mean distance to nuclear lamina and the proportion of genes actively transcribed. This revealed that the larger the distance from the nuclear lamina the larger the amount of genes with active transcription. Finally, we performed functional annotation of genes with burst on and located at the core of the nucleus. This revealed cell-fate, differentiation, cell-adhesion genes are transcribed actively at the single cell level despite heterogeneity in local structural measurements (*77*).

**Human brain Single nuclei RNA sequencing data analysis**

We downloaded snRNAseq data (gene expression and metadata) from 24 human brain control samples (*76*). We re-analyzed the data using SEURAT (*185*) and performed cluster-based separation of different subtypes, followed by identification of cell types based on the canonical marker genes and the annotation provided in the paper. The differentially expressed genes for each cell type were identified by FinderMarker function in Seurat package. These genes were used for further analysis in this study.

**Transcriptomic profile of human adipose precursor cells**

To compare different human adipocyte progenitor cells (APC) we used previously defined subtypes: APC beige-like, APC low lipid turnover and APC high lipid turnover (*75*). These categories represent different phenotypic behavior such as beiging ability and lipid turnover activity (*75*), and were defined using FACS analysis, transcriptomic, proteomic and metabolic profiling. We therefore downloaded datasets from this study from GEO: GSE129042, which corresponds to expression profiling by high throughput sequencing of FACS-sorted APCs from adipose stromal vascular fraction of human subjects. Counts were processed and differentially expressed genes determined using the DEGSeq package in R as previously described (*186*) For differentially expressed genes in each APC subtype we performed functional annotation and aggregate topological expression within fPGEs composing our reconstructed nuclear compartments (see aggregate topological expression).

**Adipose tissue scRNAseq from obese human subjects**

We downloaded the snRNAseq data (gene expression and metadata) from > 20 samples from adipose tissue biopsies of obese human subjects (*90*). We re-analyzed the data using Seurat (*185*) and separated cell types based on the canonical marker genes provided in the paper. The differentially expressed genes for each cell type were then identified by FinderMarker function in Seurat package. These genes were used for further analysis in this study.

**Adipose tissue scRNAseq from mice**

We downloaded the scRNAseq data (gene expression and metadata) from different tissues of the tabula muris consortium (*78*). We re-analyzed data from tissues of interest using seurat (*185*) and separated cell types based on the canonical marker genes provided in the paper. The differentially expressed genes for each cell type were then identified by FinderMarker function in Seurat package. These genes were used for further analysis in this study.

**Quantification and Statistical Analysis**

Statistical analyses related to experimental procedures were performed using GraphPad Prism Software (San Diego, CA) and all parameters are indicated in the corresponding figure legend. Quantitative data are presented as the mean ± SEM and n is indicated for each experiment. All experiments were carried out with 3 biological replicates and 3 independent times. Animal experiments were performed in more than 3 animals per group. Unpaired Student's t test was used to determine statistical significance when two groups were compared. One-way ANOVA followed by Fisher's least significant difference (LSD) test for post hoc comparisons was used to determine statistical significance when multiple groups were compared. Two-way ANOVA was used to estimate significance between groups constrained by time-based measurements, followed by Tukey test for post-hoc multiple comparisons. Correlations were calculated by Spearman rank and Pearson correlation with a two-tailed test. Statistical significance was defined as p value < 0.05 by either test and is denoted with an asterisk as follows: * p-value 0.05-0.01, ** p-value 0.01-0.001 and *** < 0.001 when indicated.

**Data and Code Availability**

The analysis was performed in R (3.5 and 3.6) and python (2.7, 3.5 and 3.7). Libraries for data analysis include numpy, pandas, scikit-learn, scanpy and scipy. The datasets reported in this paper are available in supplementary materials. Analysis software will be released upon publication at https://github.com/leandroagudelo189. When indicated, network analysis and visualization were done using the python library NetworkX, and the software GEPHI. Functions to evaluate networks have been created and validated by the Ideker Lab (archived in https://github.com/idekerlab/Network_Evaluation_Tools) and are stored on the network data exchange (NDEx) (*187*). Other functions to evaluate network topology can be found as inbuilt analysis tools in GEPHI. Tissue-specific networks for functional annotation of topological dependencies can be found at http://giant.princeton.edu and https://hb.flatironinstitute.org/. Positional gene enrichment analysis tool can be found at http://silico.biotoul.fr/pge/. Functional annotation tool from a large library of biological repositories can be found at https://maayanlab.cloud/Enrichr/. DAVID functional clustering analysis tools can be found at https://david.ncifcrf.gov/. To filter out PGE regions without cytogenetic band enrichment, we used a curated list of cytogenetic band-gene dependencies archived in http://www.gsea-msigdb.org/gsea/msigdb/index.jsp. Prediction of TFBS from DNA sequences can be found at http://alggen.lsi.upc.es/. Prediction of conserved TFBS in promoter regions of genes of interest can be found at http://www.gsea-msigdb.org/gsea/msigdb/index.jsp. Tissue-naïve and tissue-specific protein-protein interactions used for n+1 network reconstruction can be found at http://www.interactome-atlas.org/. Tissue-naïve protein-protein interactions used for full network reconstruction can be found at https://thebiogrid.org/. In silico prediction of intrinsically disordered regions in amino acid sequences can be found at http://www.pondr.com/, https://iupred3.elte.hu/, https://mobidb.bio.unipd.it/. Curated experimentally validated phase separation database stored at http://db.phasep.pro/. Context-dependent phase separation score from ensemble parameters of conformation changes present in disordered proteins. They offered discrepancies of conformations that are related to the local environment such as salt, solvation, multi-molecular interactions and ligand presence. These can be found at http://pappulab.wustl.edu/CIDER/about/ and can be implemented locally following instructions stored at http://pappulab.github.io/localCIDER/. Phylogenetic analysis tool information for protein residues conservation score and 3D modelling of residue exposure can be found at https://consurf.tau.ac.il/. Evolutionary domain conservation of protein domains with known structure can be found at https://consurfdb.tau.ac.il/. ChIP-atlas methodology, documentation and functions for analysis can be found at https://github.com/inutano/chip-atlas. Compendium of human chromatin contact maps has been created by the Ren Lab and is stored at GEO: GSE87112 (schmitt, bing ren 2016). GREAT software, available at http://great.stanford.edu/public/html/index.php, was used to evaluate biological process enrichment for neighboring genes compared to genomic background. Cis-regulatory elements relationship were obtained from GeneHancer, and can be found at http://www.genecards.org/. VISTA enhancer elements can be found at https://enhancer.lbl.gov/frnt_page_n.shtm. HACER, human active enhancer to interpret regulatory variants database is archived in http://bioinfo.vanderbilt.edu/AE/HACER/index.html. A curated database of cell-specific active regulatory regions can be found at http://www.enhanceratlas.org/index.php. Global chromatin 3D reconstruction tool and code can be found as follows: GSDB at http://sysbio.rnet.missouri.edu/3dgenome/GSDB/; and Spacewalk at http://3dg.io/spacewalk/. 3D-multiscale Monte Carlo modeling code for global and local CTCF-mediated ChIA-PET sequencing can be found at https://bitbucket.org/3dome/3dome_mmc/src/master/. Code for integration of structural variants (SVs) to spatial chromatin interaction patterns can be found at https://bitbucket.org/4dnucleome/spatial_chromatin_architecture. Data by Sadowski et al. on SVs spatial chromatin effects can be found at https://zenodo.org/record/2837248#.YCQBvZNKhPM. CTCF-binding site prediction tool can be found at https://insulatordb.uthsc.edu/. Structural variants data from the reference resource compendia developed by Collins et al. (*55*) is archived in https://gnomad.broadinstitute.org/. To isolate metabolic disease associations for a given genomic-range, we used the curated common metabolic disease portal, archived in https://hugeamp.org/. Maps of epigenomic integration of Encode and Roadmap consortium data, EpiMap can be found at http://compbio.mit.edu/epimap/. For topological comparative genomics and to evaluate chromosomal synteny, tools can be found at the synteny portal, archived in http://bioinfo.konkuk.ac.kr/synteny_portal/. CAMMEL mediation analysis and polygenic risk scores can be found at https://github.com/YPARK/GTEx_mediation and https://github.com/YPARK/PRS. We integrated in our pipelines previously published data derived from radial genome loci positioning by sequencing (GPSeq) method, which gives relative distances between genomic-loci and the nuclear lamina (*80*) (GEO:GSE135882; GPSeq protocol available at https://doi.org/10.21203/rs.3.pex-570/v1; data available at https://github.com/ggirelli/GPSeq-source-data). To evaluate radial positions of transcriptional active loci in single cells and their structural relationships, we used datasets from established genome-scale approaches that use multiplexed FISH imaging technology (*77*) (available at https://zenodo.org/record/3928890#.YDgeiZNKiuV and script functions can be found at https://github.com/ZhuangLab/Chromatin_Analysis_2020_cell)

### Supplementary Figures

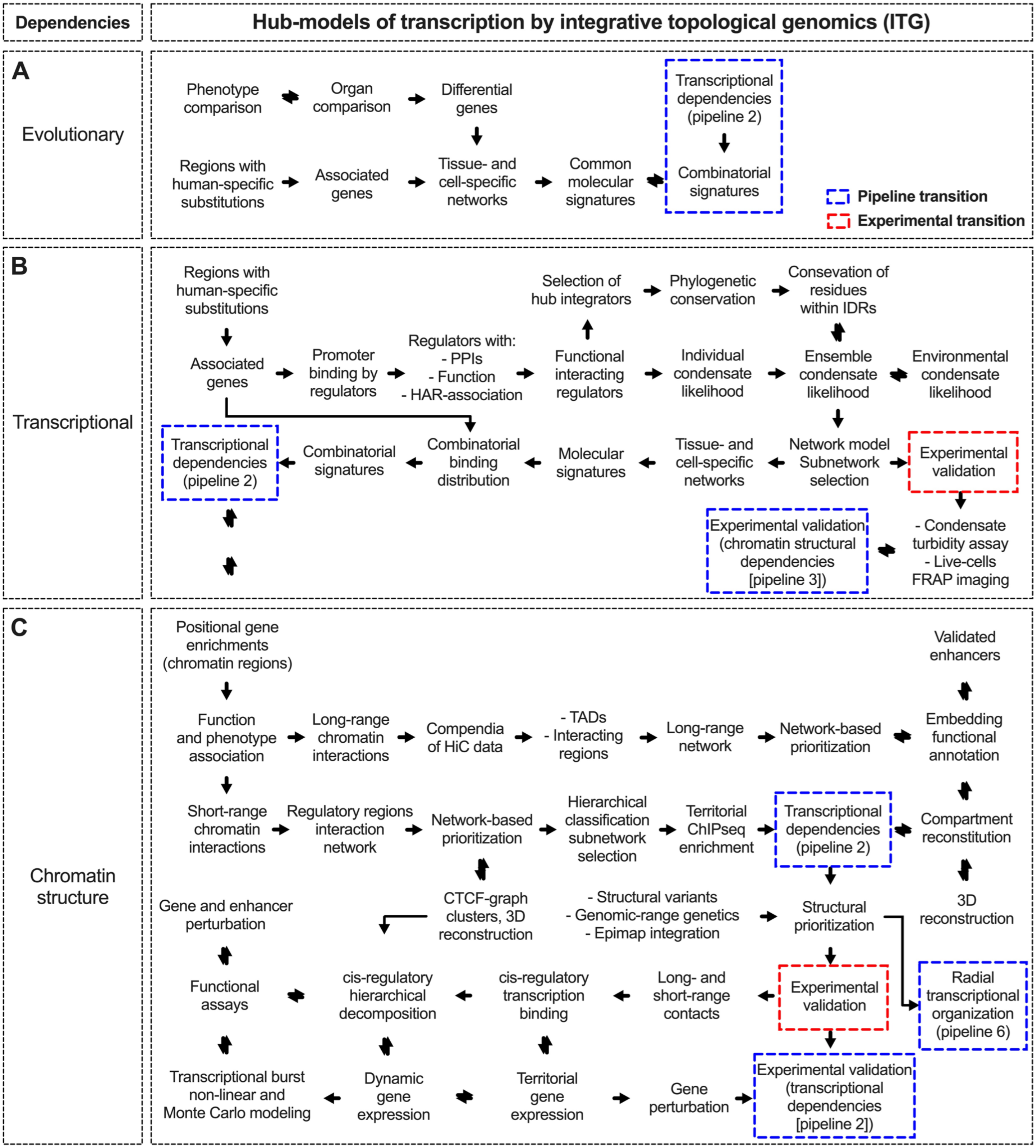


**Fig. S1. Integrative topological genomics overview: Part 1.** (**A**) Representation of pipeline steps for evolutionary relationships. (**B**) Representation of pipeline steps for transcriptional relationships. (**C**) Representation of pipeline steps for chromatin structural relationships

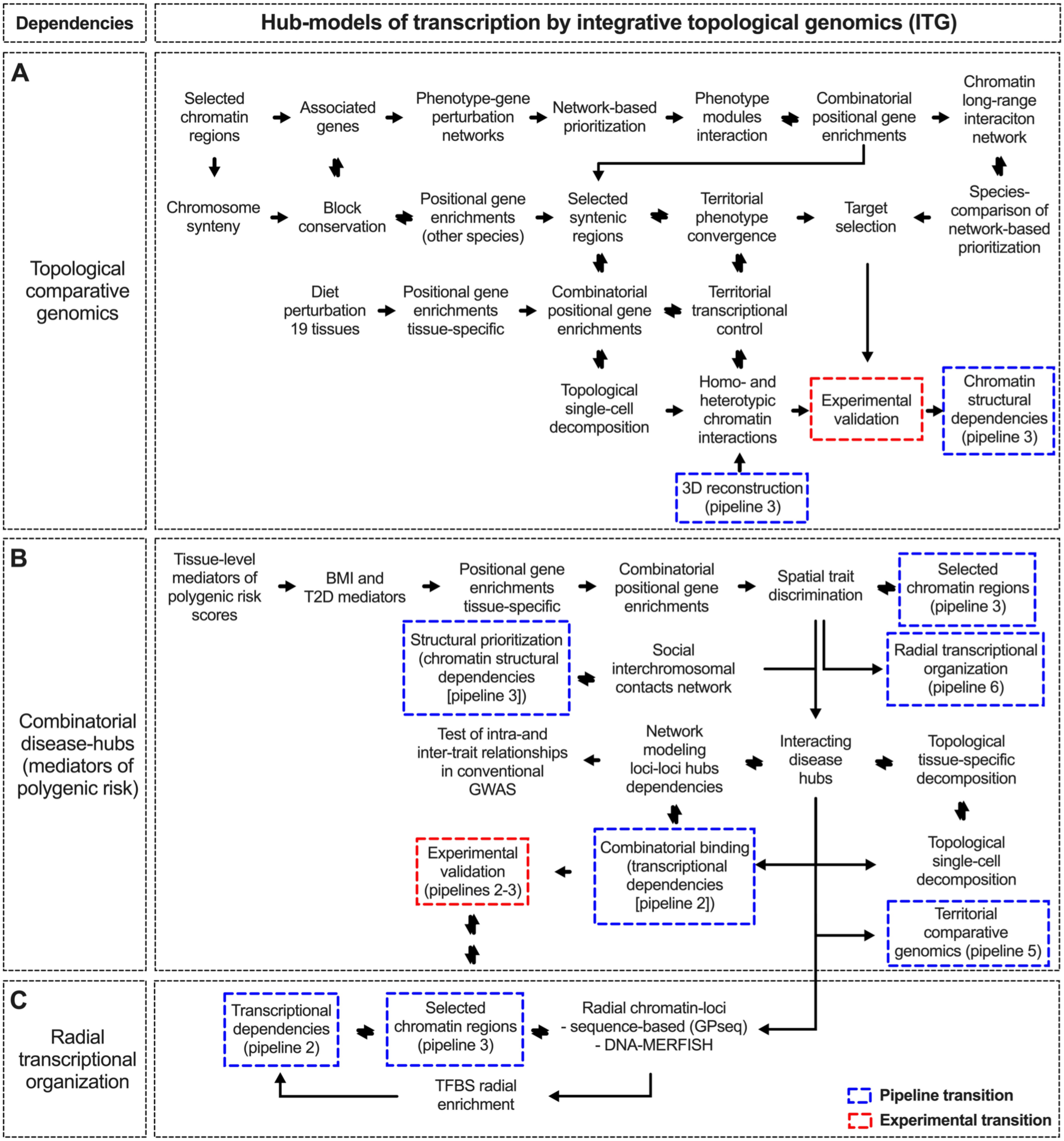


**Fig. S2. Integrative topological genomics overview: Part 2.** (**A**) Representation of pipeline steps for topological comparative genomics relationships. (**B**) Representation of pipeline steps for disease mediators of polygenic risk relationships. (**C**) Representation of pipeline steps for radial chromatin relationships

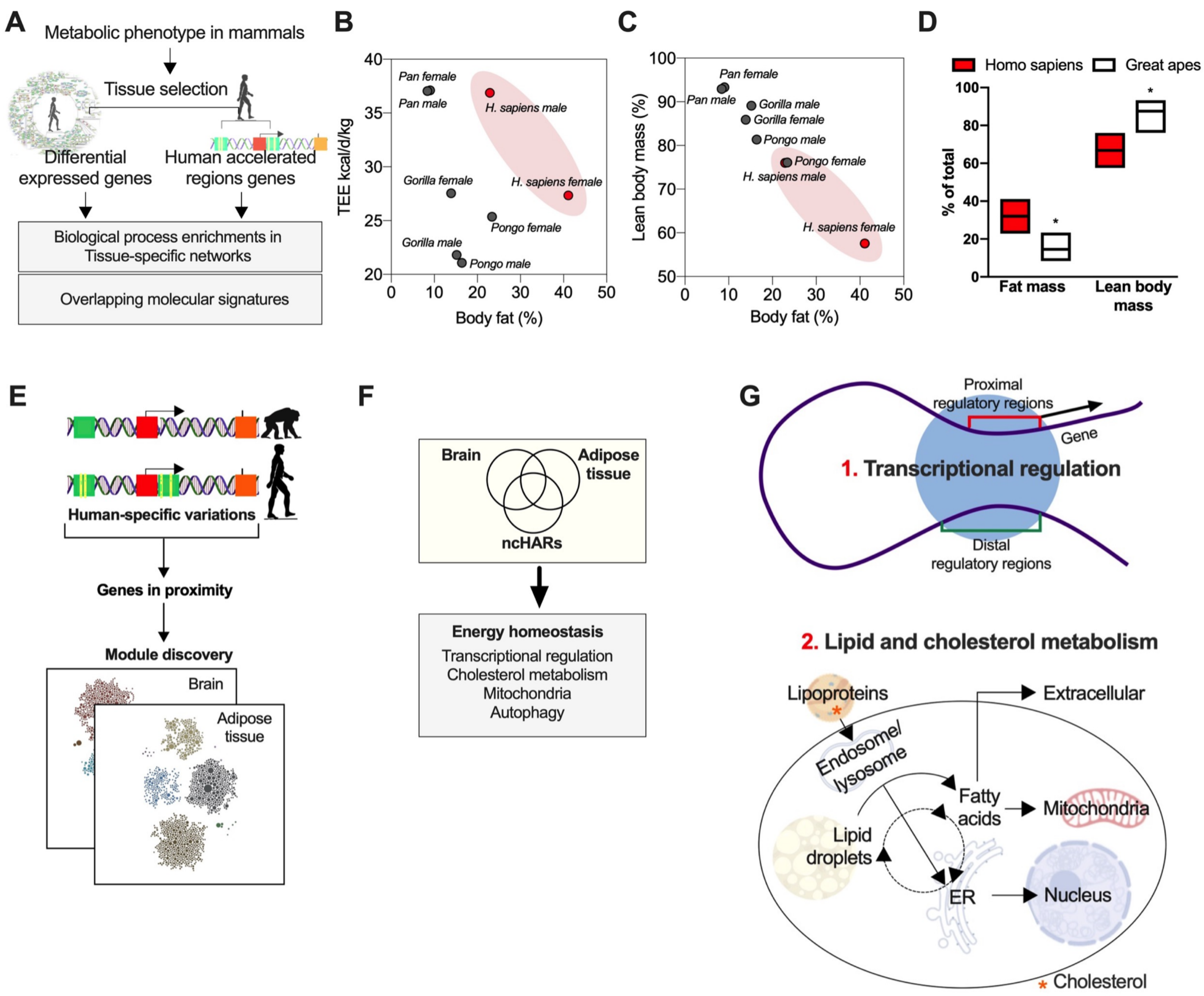


**Fig. S3. Phenotypic and genetic human accelerated metabolism** *(related to Fig. 1).* (**A**) Schematic for network-based integration of phenotype and genetic human enriched biological enrichments (part of evolutionary relationships pipeline). (**B**) Body fat composition in higher primates and relationship to total-energy expenditure and lean body mass (*23*). (**C**) Comparison of fat mass and lean body mass between homo sapiens and great apes (*23*). (**D**) Body fat composition in higher primates and relationship to total-energy expenditure and lean body mass. (**E**) Illustration showing non-coding human accelerated regions and genes in proximity followed by network based functional enrichments (part of evolutionary relationships pipeline). (**F**) Overlap of network based functional enrichments between molecular signatures increased in humans compared to primates and genes in proximity to HARs. (**G**) Schematic illustration of molecular signatures observed in (**F**) and in **Fig. 1C**. Bars show mean values. Unpaired, two-tailed student's t-test was used when two groups were compared, and ANOVA followed by fisher's least significant difference (LSD) test for post hoc comparisons for multiple groups. * p-value <0.05.

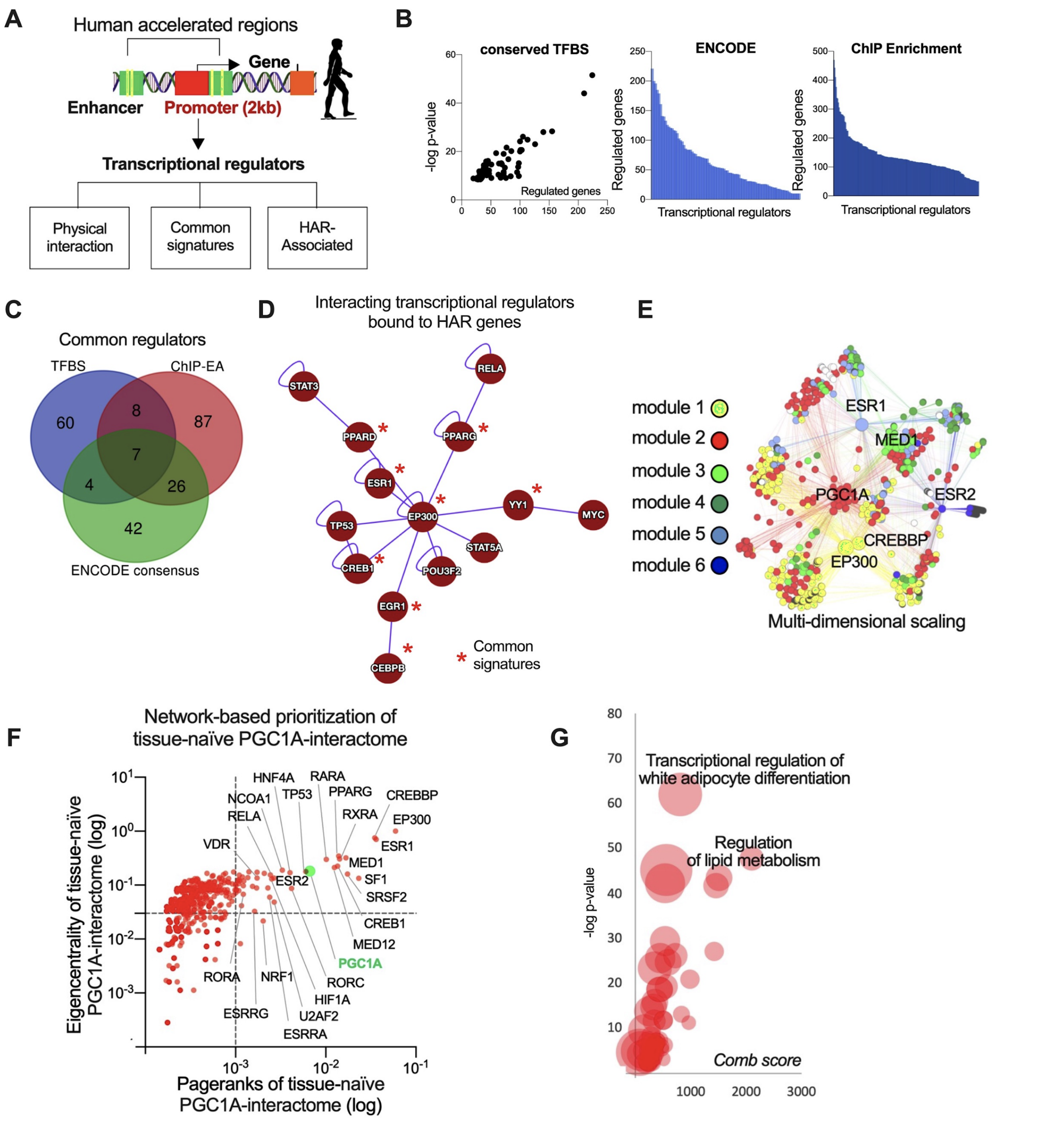


**Fig. S4. Transcriptional dependencies of human accelerated regions genes** *(related to Fig. 1)*. (**A**) Illustration showing genes in proximity of HARs and interrogation of transcriptional dependencies on their promoter regions (part of transcriptional relationships pipeline). Bound regulators with physical interactions, similar function and HAR-association. (**B**) Binding dependencies of transcriptional regulators of ncHAR-genes by TFBS-conservation, ENCODE ChIP-seq binding, and ChIP-seq atlas binding enrichment. (**C**) Venn diagram of common regulators from (**D**). Network reconstitution of common regulators of HAR genes displaying physical interactions and similar function. (**E**) Full reconstitution of interactors of common regulators followed by network analysis and representation using multi-dimensional scaling. (**F**) Network-based prioritization of interacting regulators by eigencentrality and pagerank. (**G**) Functional enrichment of top nodes from (**F**).

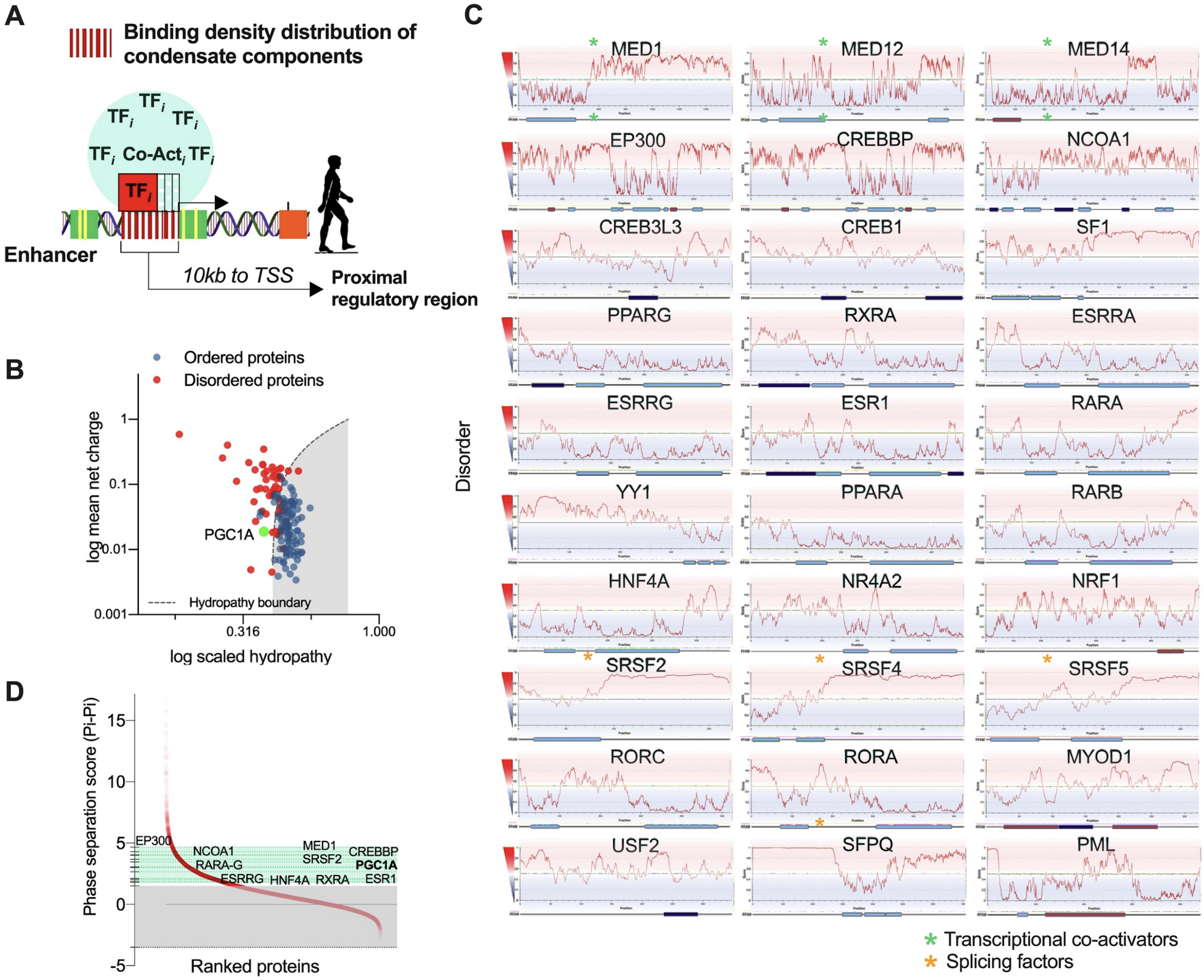


**Fig. S5. Disorder scores for interacting transcriptional regulators of HAR genes** *(related to Fig. 1).* (**A**) Illustration on regulatory binding density distribution for HAR-genes promoter regions (10kb) (part of transcriptional relationships pipeline). (**B**) net charge and scaled hydropathy of hub integrator the co-activator PGC1A, along with ordered and disordered regions from PONDR software. (**C**) Disorder scores for prioritized nodes from fully reconstituted interacting regulators network. (**D**) Genome-wide phase separation scores based on highly frequent pi-interactions in small amino acids with exposed peptide backbone (*111*) Highlighted in green range score for interacting transcriptional regulators of HAR genes.

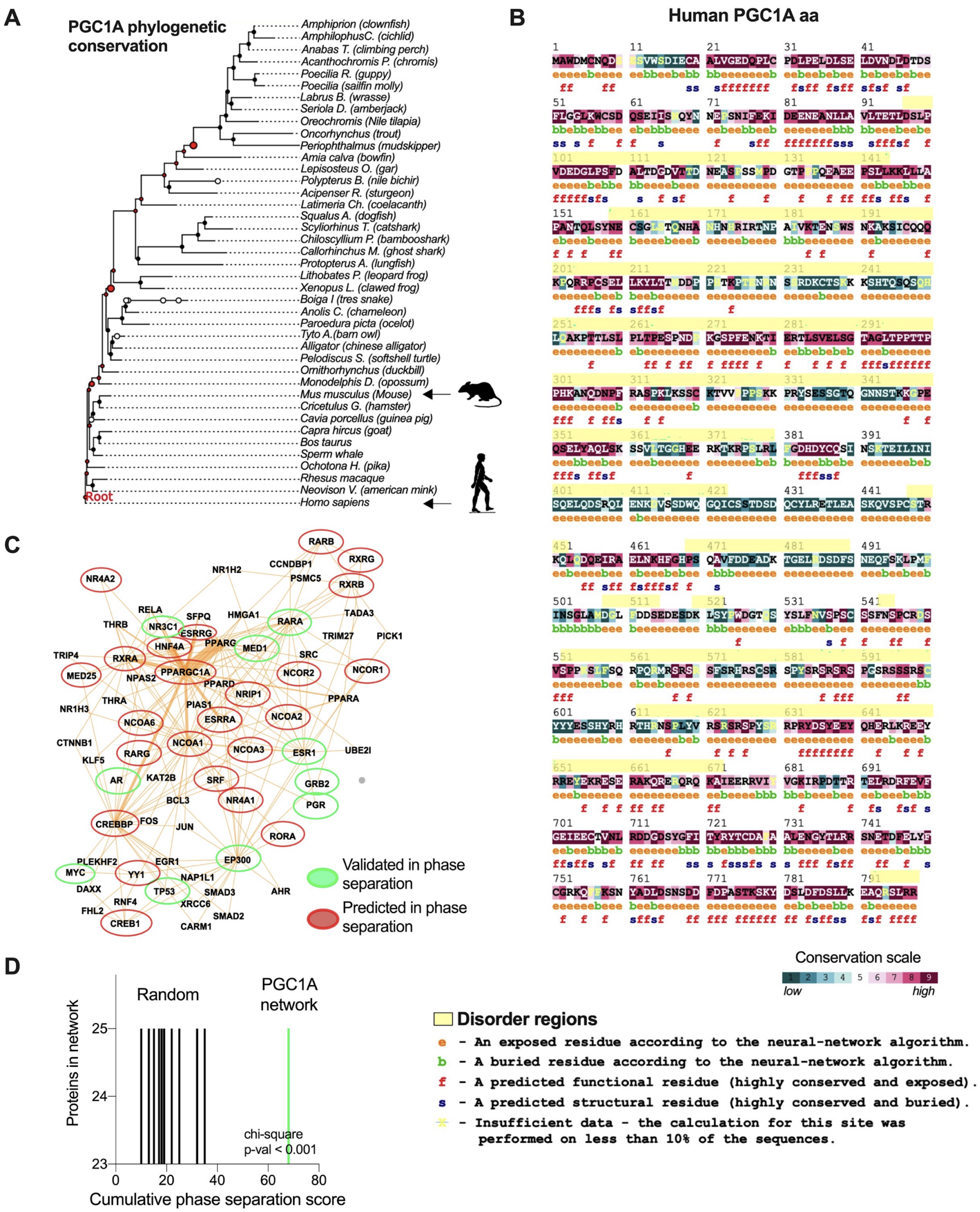


**Fig. S6. Phylogenetic conservation of hub integrator PGC1A co-activator and cumulative phase separation scores of its network** *(related to Fig. 1).* (**A**) PGC1A amino acid residues phylogenetic conservation using CONSURF. (**B**) Conservation scores in PGC1A amino acids from phylogenetic analysis. In yellow consensus disordered regions. (**C**) Brain and adipose tissue backbone PPI network, displaying interacting PGC1A regulatory network, and proteins undergoing phase separation validated experimentally in green and predicted in silico in red. (**D**) Cumulative phase separation score for PGC1A-interacting network.

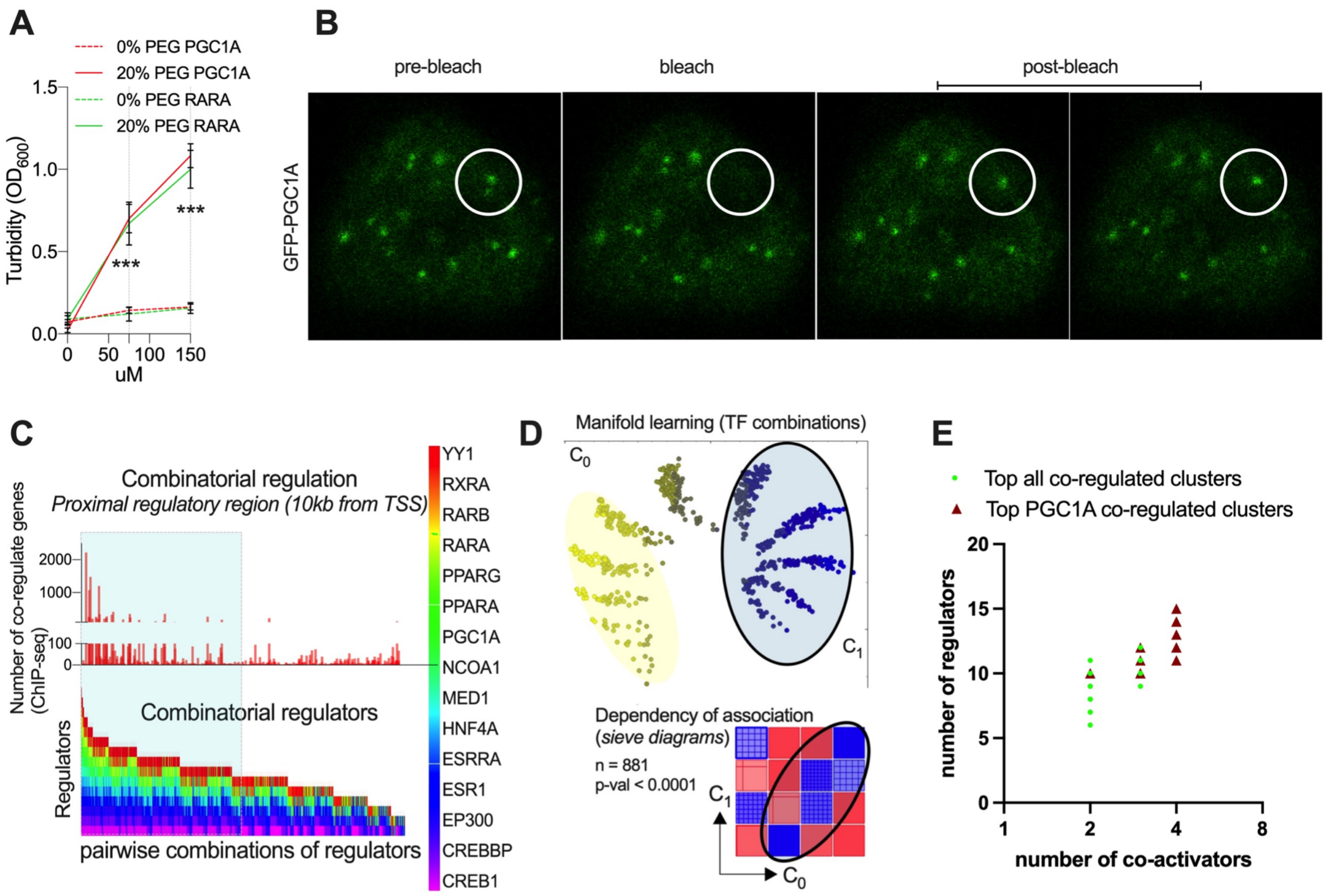


**Fig. S7. PGC1A phase separation and PGC1A-network combinatorial binding** *(related to Fig. 1)***.** (**A**) Turbidity assays of PGC1A and RARA protein at different concentrations and with and without polyethylene-glycol. (**B**) FRAP images of GFP-PGC1A condensates in HEK293 cells and treated with forskolin for 4h, and dynamic recovery of luminescence. (**C**) Combinatorial binding of PGC1A-interacting regulators showing the number of co-regulated genes from ChIP-seq experiments and pairwise combination of interactors. Candidates of regulators selected from network analysis. (**D**) Manifold representation of combinations of transcriptional regulators showing in blue top clusters with combinations co-regulating a higher number of genes. Below, sieve diagram representations assessing the dependence of association found in top clusters 1. (**E**) Top co-regulated clusters generally (all combinations) and targeted (combinations with PGC1A). Bars show mean values and standard error of the mean. Two-way anova was performed to compare the effect of group and concentration between groups, followed by two-tailed student's t-test by Tukey's multiple comparison test. P-value cutoff was set at <0.05. *** representing <0.001.

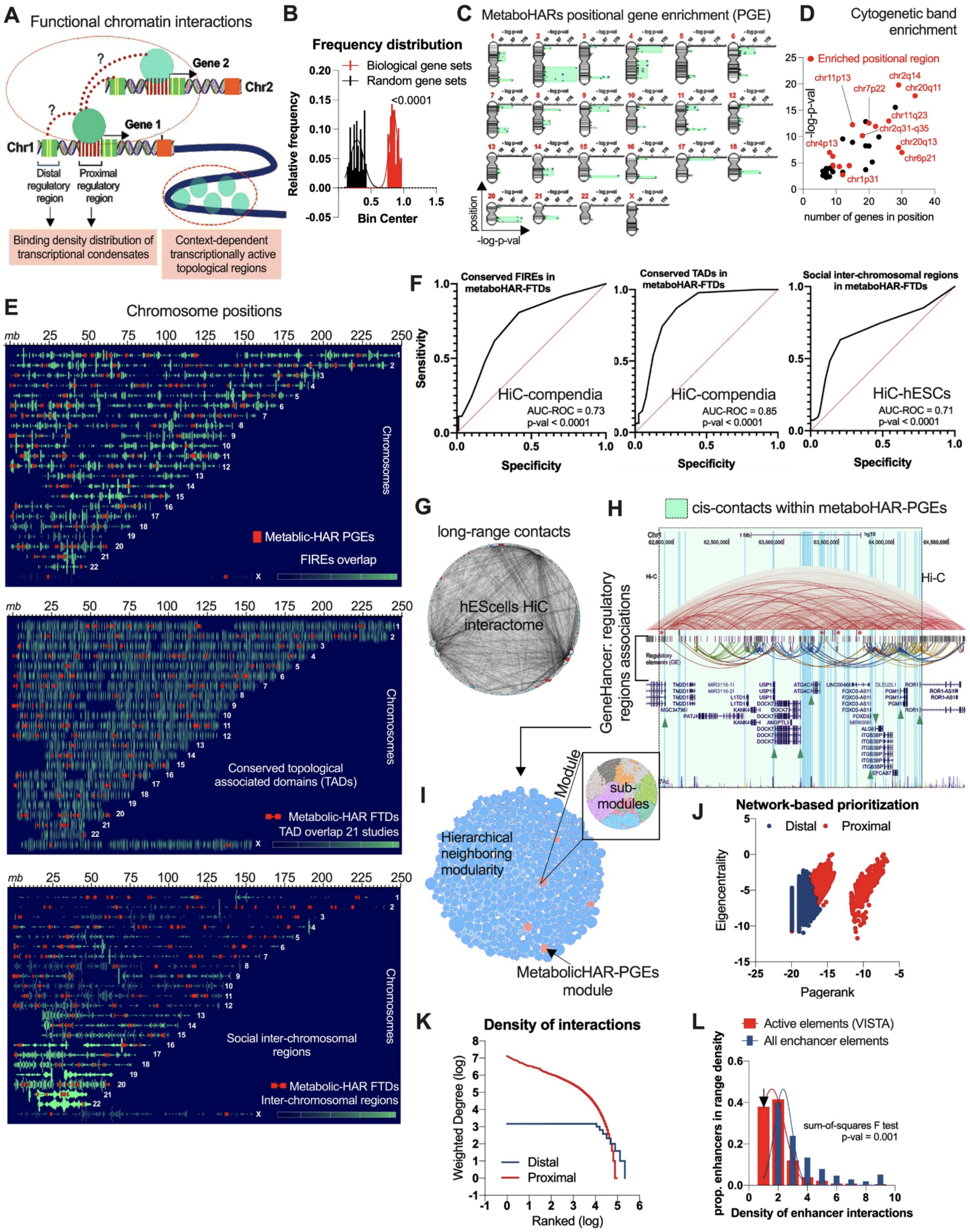


**Fig. S8. Interrogation of functional nuclear compartments through topological integration** *(related to Fig. 2)*. (**A**) Illustration showing pipeline for chromatin structural relationships. (**B**) Frequency distribution for positional gene enrichments (PGEs; > 5 genes per genomic cluster) comparing biologically derived data sets and random gene sets. (**C**) Metabolic HAR genes positional enrichments. (**D**) Cytogenetic band enrichments for metabolic-HAR genes. (**E**) Metabolic-HAR PGEs relationships with compendia of frequently interacting chromatin regions (FIREs), conserved TADs and social trans-interacting chromatin regions (see pipeline for chromatin structural relationships). (**F**) Area under the curve of the receiver operating characteristic (AUC-ROC) for analysis comparing the presence of regions from (**E**) in PGEs from metabolic-HAR genes set and random gene sets. (**G**) Trans-interacting chromatin region network from hESCs displayed in a circular layout. Node size represents pagerank analysis. (**H**) Metabolic HAR domain in chr1: 62-65mb, showing regulated genes by PGC1A-interacting network (green arrow), representative contact HiC frequency maps (GM12878), genehancer regulatory elements and interactions along with vista enhancers. (**I**) Genome-wide network of regulatory element interactions from Genehancer database, displayed in modules and with metabo-HAR regions in red. (**J**) Network-based prioritization of regulatory elements classified in proximal comprising promoter, and distal for enhancer regions. (**K**) Density of interactions by degree parameter for distal and proximal regulatory elements. (**L**) Overlapping consensus of transcriptional regulators. (**L**)  Number of links among genome-wide distal regulatory regions and ncHAR elements.

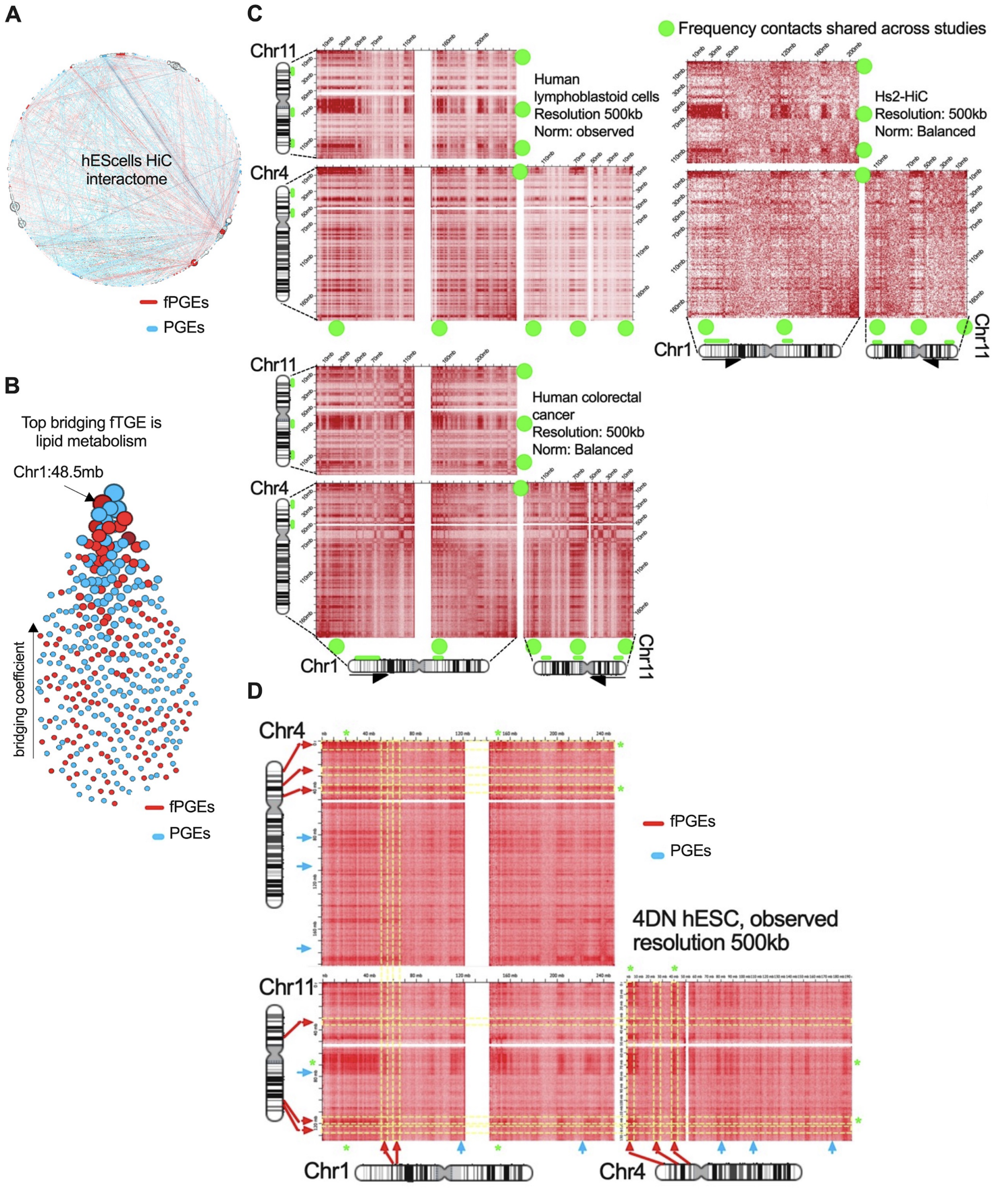


**Fig. S9. Top metabolic functional nuclear compartments** *(related to Fig. 2)*. (**A**) hESCells HiC network of trans-interacting chromatin regions, displaying overlapping positional enrichments. In red metabolic-HAR PGEs with similar functional annotation and in blue for the rest. (**B**) Bridging coefficient for overlapping metabolic HAR PGEs in trans-interacting chromatin regions. Layout upwards with top PGEs with highest score, and functional (red) or not (blue). (**C**) Pervasive of contact frequency maps across other studies for metabolic-HAR regions with similar function (top bridging coeff.) and in trans-interacting regions. (**D**) For selected chromosomes harboring metabolic-HAR regions with similar function, displaying the location of PGEs in relation to trans-interacting contacts.

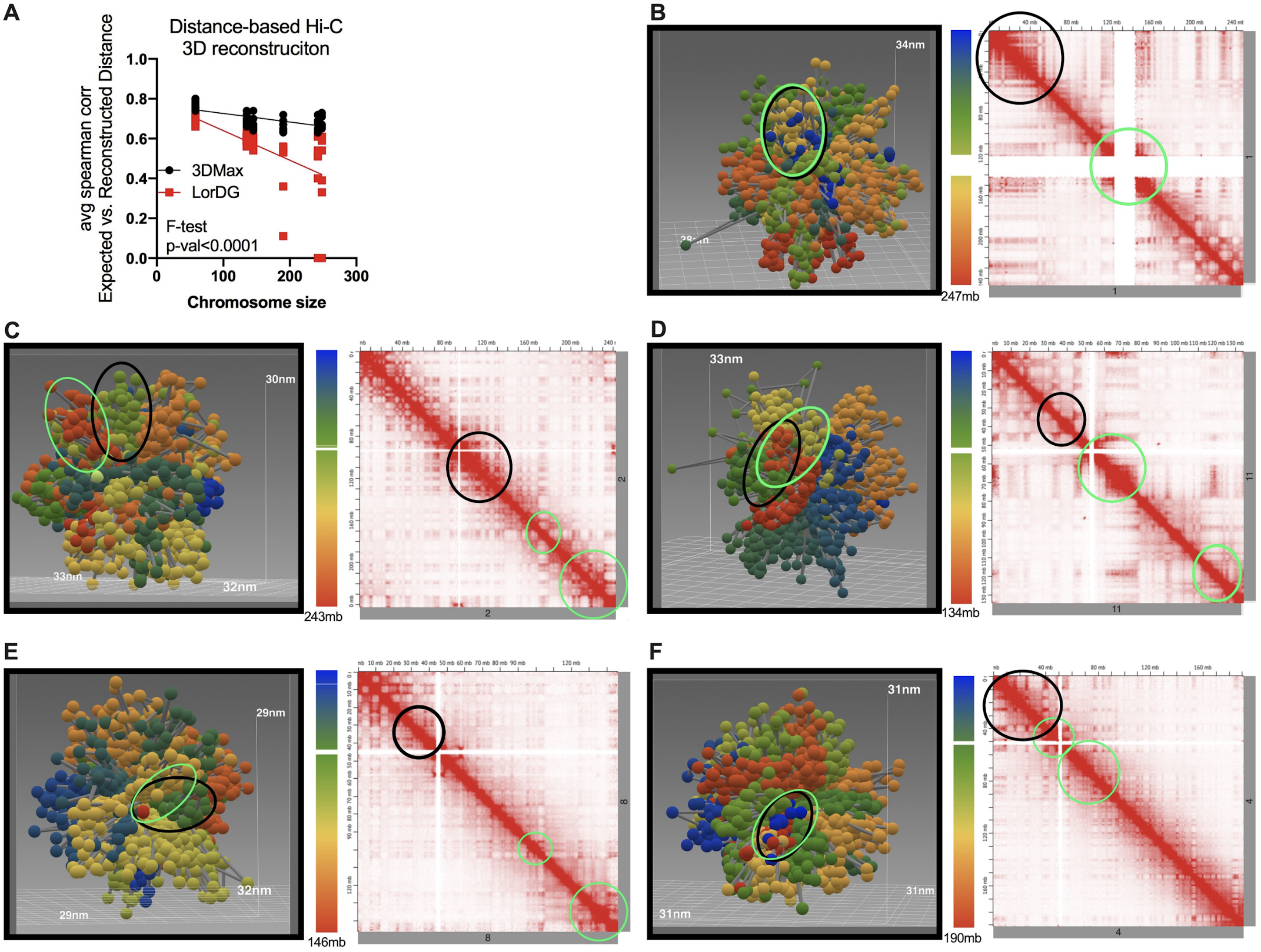


**Fig. S10. Global 3D reconstruction of chromosomes harboring metabo-HAR regions and their homotypic interactions** *(related to Fig. 2)*. (**A**) Comparison of distance based 3D HiC reconstruction for chromosome size biase, between 3DMax and LorDG softwares. Displayed expected vs reconstructed distances and chromosome size. 3D picture reconstruction using 3DMax for chromosomes harboring selected metabolic-HAR regions (black circle) and top homotypic interaction (green circle) for Chr1 (**B**), Chr2 (**C**), Chr11 (**D**), Chr8 (**E**) and  Chr4 (**F**).

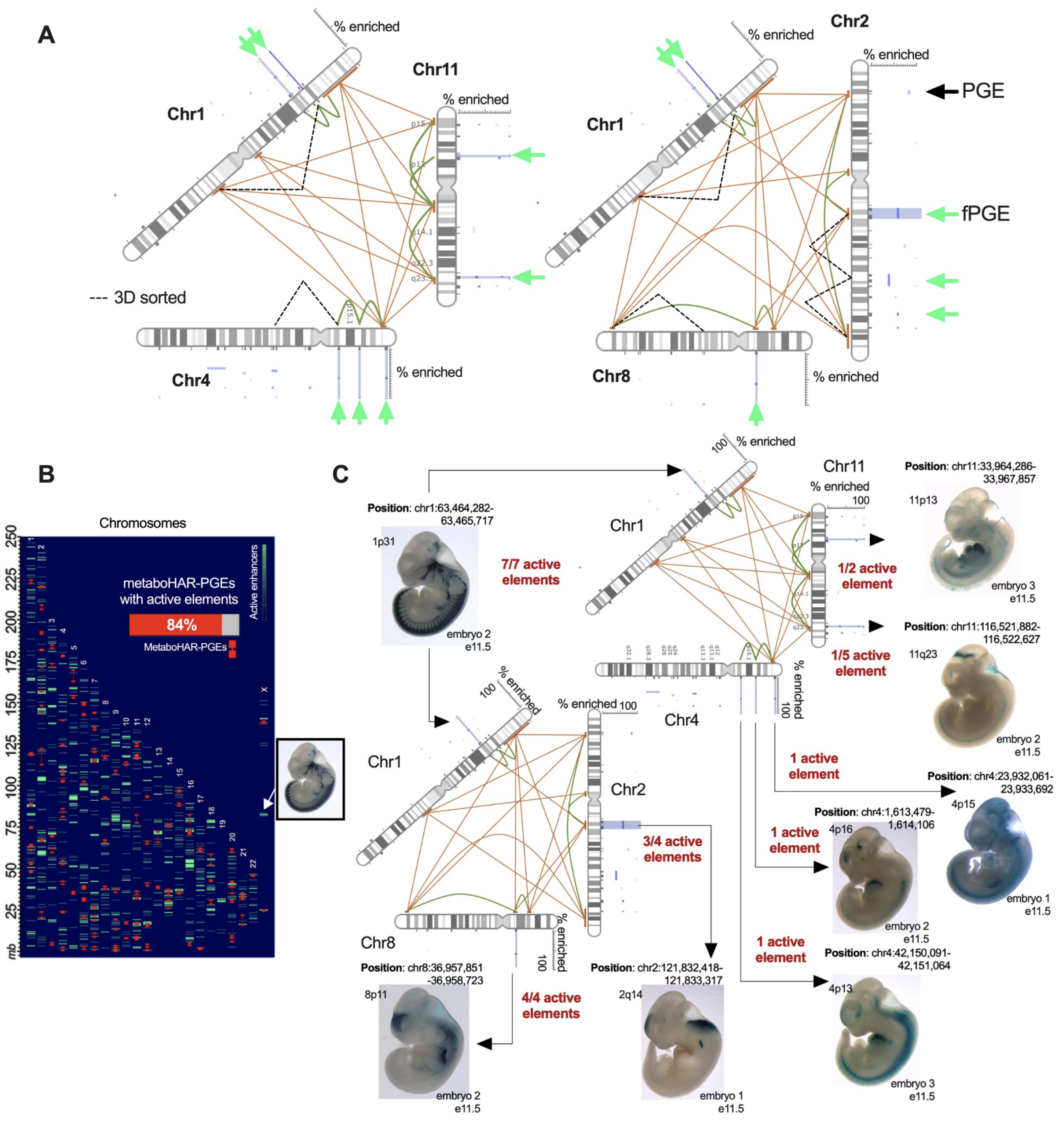


**Fig. S11. Reconstructed functional nuclear compartments and experimentally validated enhancer activity within regions** *(related to Fig. 2)*. (**A**) Schematic representation for bridging-score mediated reconstruction of interactions between chromosomes harboring metabolic-HAR domains with similar function. Displayed fPGEs and PGEs in green and black arrows respectively, homo- and heterotypic interactions, along with 3D interacting prioritization. (**B**) Metabolic-HAR domains relationships with active enhancers validated and found in the VISTA database. Displayed regions with high density of active enhancers and percentage overlap with all metabolic HAR domains. (**C**) Schematic representation of reconstructed functional compartments displaying number of active elements within metabo-HAR region and representative picture of enhancer activity found in VISTA database.

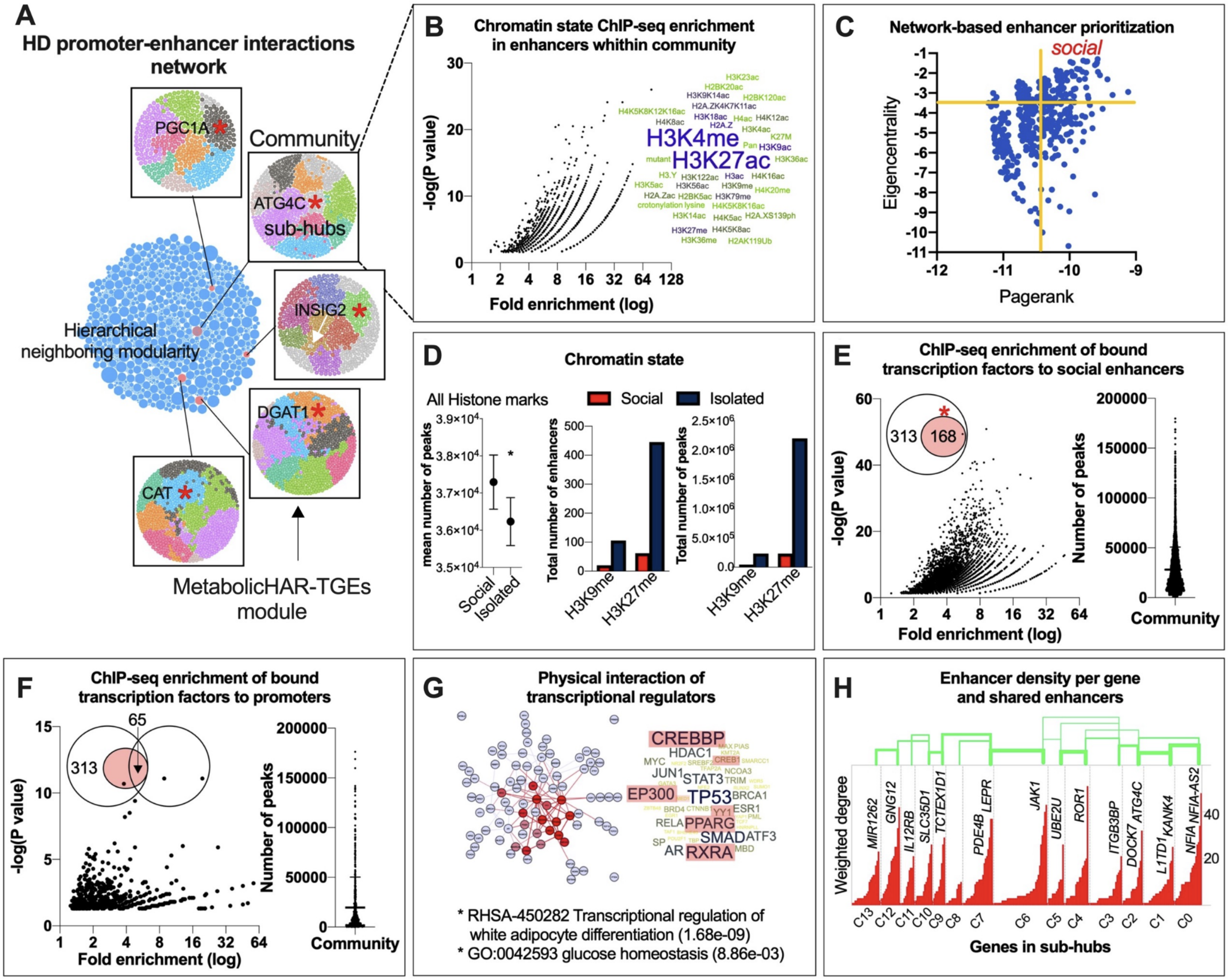


**Fig. S12. High-definition short-range territorial networks** *(related to Fig. 2)*. (**A**) Short-range regulatory elements interaction network, displayed with a modularity based layout. Highlighted in red metabolic-HAR domains modules and submodule territories. Gene in proximity to HAR elements and submodule location (asterisk). (**B**) ChIP-seq enrichment analysis for elements composing metabolic-HAR domain and specific for chromatin states. In a word-cloud subpanel represented top chromatin modifiers within elements. (**C**) Network-based prioritization of metabolic-HAR regions module. (**D**) Chromatin state ChIPseq enrichment comparison for highly social and isolated enhancers. First right panel showing peaks for all histone marks, number of enhancers and number of peaks for repressive histone marks H3K9me and H3K27me. (**E**) Transcriptional regulator ChIP-seq enrichment analysis for social enhancers within metabolic-HAR module and number of peaks within module. Venn diagram showing number of regulators across all elements (313) and top regulators in social regulators (168) (**F**) Transcriptional regulator ChIP-seq enrichment analysis for promoters within metabolic-HAR module and number of peaks within module. Venn diagram showing number of regulators across all elements (313) and overlapping top regulators in promoter and social enhancers. (**G**) PPI Network for interacting transcriptional regulators selected in F. Word-cloud of top regulators (in red) and functional annotation. (**H**) Promoter gene degree distribution of submodules within metabolic-HAR region and density of shared enhancers between submodules.

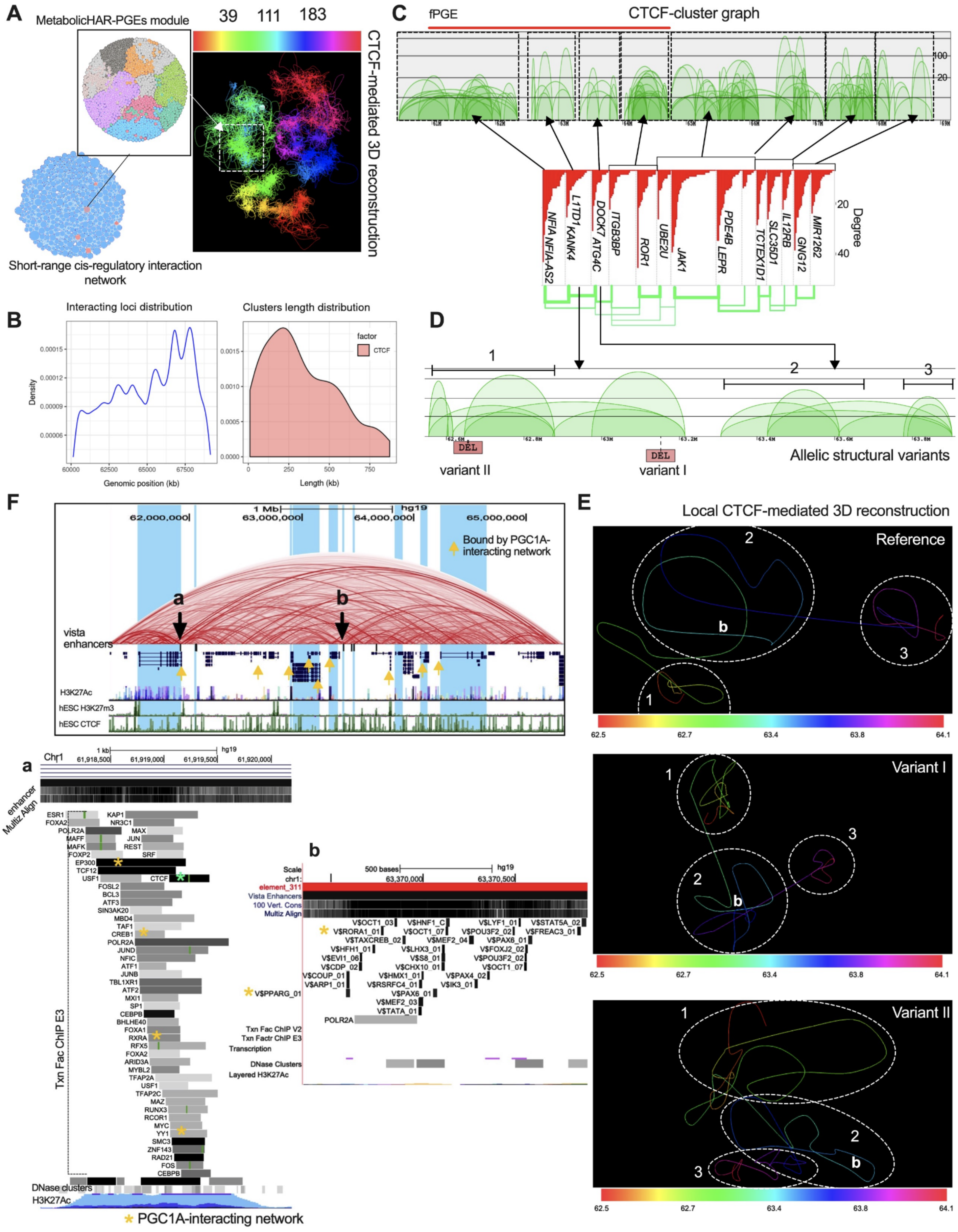
A
MetabolicHAR-PGEs module
39 111 183
CTCF-mediated 3D reconstruction
Short-range cis-regulatory interaction network
B
Interacting loci distribution
Clusters length distribution
Density
Genomic position (kb)
factor
CTCF
Length (kb)
C
fPGE
CTCF-cluster graph
NFIA NFIA-AS2
L1TD1 KANK4
DOCK7 ATG4C
ITGB3BP
ROR1
UBE2U
JAK1
PDE4B LEPR
TCTEX1D1
SLC35D1
IL12RB
GNG12
MIR1262
Degree
D
DEL
variant II
DEL
variant I
Allelic structural variants
F
1 Mb
hg19
62,000,000
63,000,000
64,000,000
65,000,000
Bound by PGC1A-interacting network
vista enhancers
H3K27Ac
hESC H3K27m3
hESC CTCF
E
Local CTCF-mediated 3D reconstruction
Reference
Variant I
Variant II
a
Chr1
enhancer
Multiz Align
Txn Fac ChIP E3
DNase clusters
H3K27Ac
PGC1A-interacting network
b
Scale
chr1:
500 bases
63,370,000
63,370,500
element_311
Vista Enhancers
100 Vert. Cons
Multiz Align
V$PPARG_01
Txn Fac ChIP V2
Txn Factr ChIP E3
Transcription
DNase Clusters
Layered H3K27Ac

**Fig. S13. Local CTCF-mediated 3D reconstructions and territorial submodules as chromatin domains** *(related to Fig. 2)***.** (**A**) Short-range network and metabolic-HAR region submodules with CTCF-based 3D reconstruction and localization of HAR domain within chromosome 1. (**B**) Density of interacting loci in genomic range, and cluster length of CTCF interactions. (**C**) CTCF-based cluster graph density of interaction within metabolic-HAR domain (top panel and fPGE in red).Submodule territorial overlap between interacting frequency network and CTCF specific clusters. (**D**) CTCF-graph cluster in 2 neighboring submodules within metabolic-HAR domain, and allelic structural variant location. (**E**) Local CTCF-based 3D reconstruction for regions in D showing reference structure and both allelic structural variant effects on density of interactions. (**F**) Genome browser picture of metabolic-HAR region showing 2 selected enhancers (a, b) for downstream analysis with TFBS and ChIP-binding location of PGC1A-interacting network.

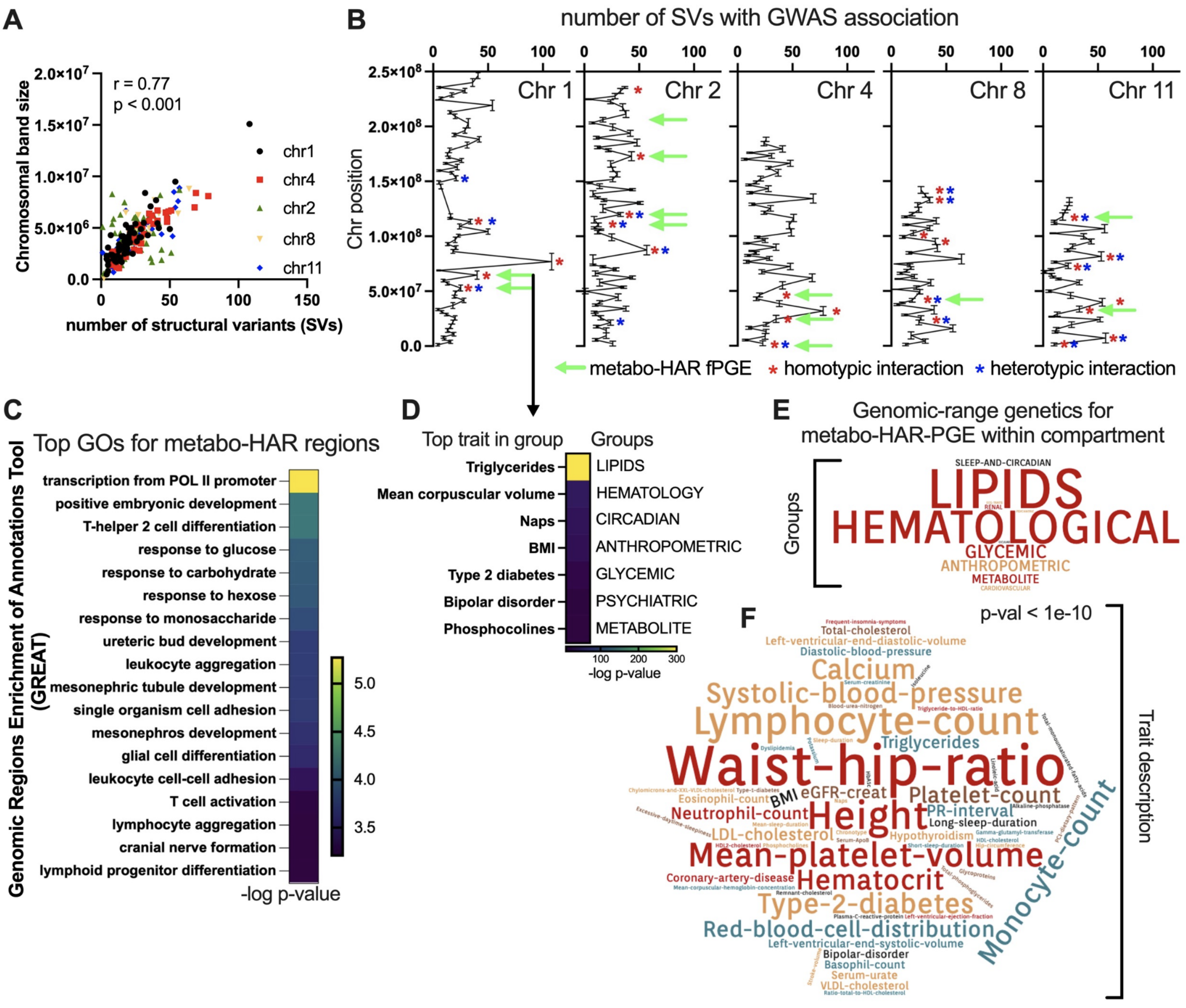


**Fig. S14. Structural variants density and genomic-range genetics in reconstructed nuclear compartments** *(related to Fig. 2)***.** (**A**) Correlation between number of structural variants with GWAS association and chromosome harboring selected metabolic-HAR domains. (**B**) Number of structural variants with GWAS association per cytogenetic band in chromosomes harboring metabolic-HAR. Green arrow shows metabolic-HAR location and red and blue asterisks for homo- and heterotypic interacting regions within compartments. (**C**) Genomic-range functional enrichment for metabolic-HAR domains using GREAT annotation tool. (**D**) Genomic-range genetics for selected metabolic-HAR regions with a representative example in Chr1:61-65mb range, using the HugeAmp tool. Displayed top trait within trait-group class. (**E**) Genomic range genetic results for selected metabolic-HAR regions within compartment displaying top trait-group repetition as a word-cloud. (**F**) Genomic range genetic results for selected metabolic-HAR regions within compartment. Displaying top traits description repetition as a word-cloud.

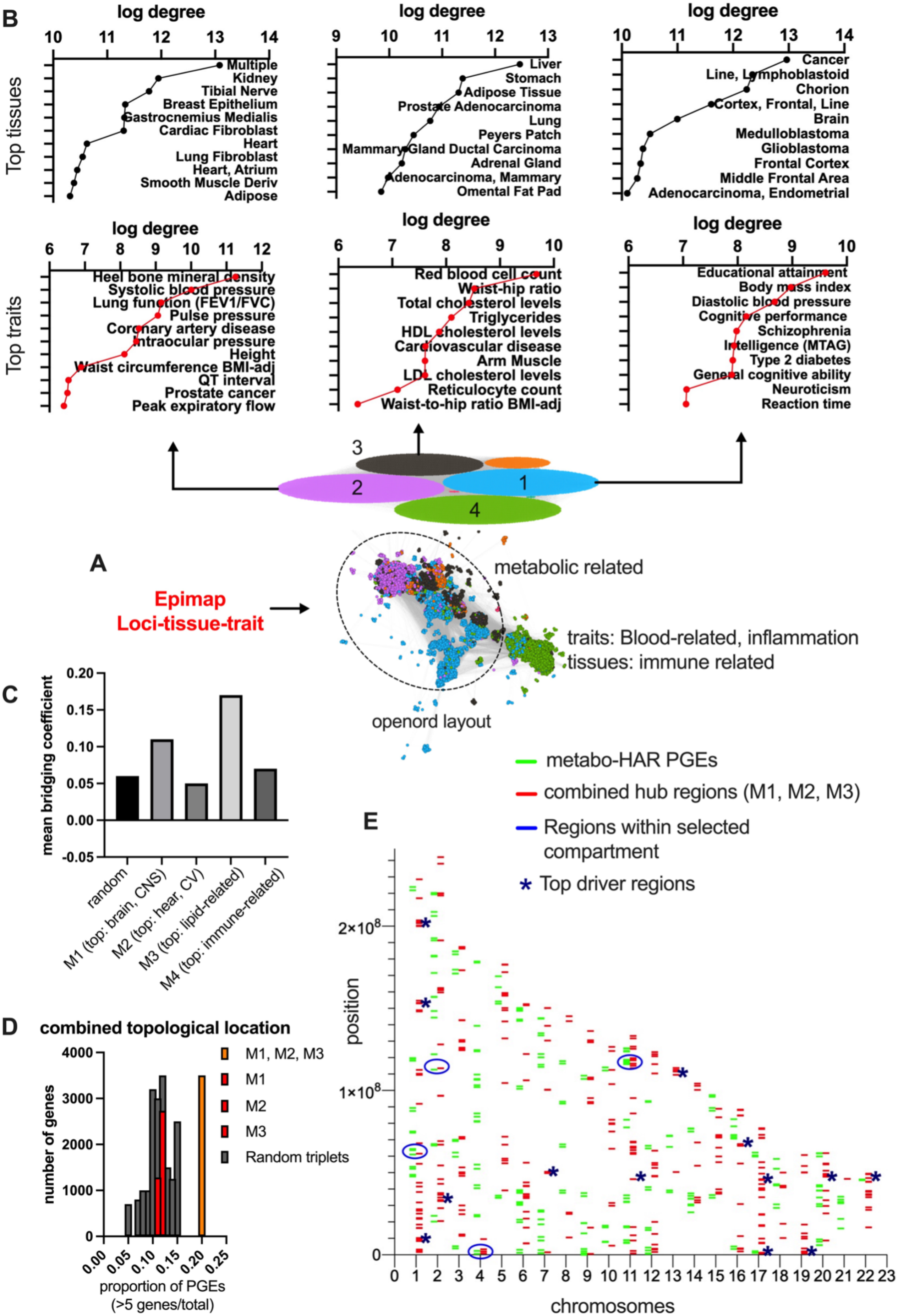

B
log degree
Top tissues
Multiple
Kidney
Tibial Nerve
Breast Epithelium
Gastrocnemius Medialis
Cardiac Fibroblast
Heart
Lung Fibroblast
Heart, Atrium
Smooth Muscle Deriv
Adipose
Liver
Stomach
Adipose Tissue
Prostate Adenocarcinoma
Lung
Peyers Patch
Mammary Gland Ductal Carcinoma
Adrenal Gland
Adenocarcinoma, Mammary
Omental Fat Pad
Cancer
Line, Lymphoblastoid
Chorion
Cortex, Frontal, Line
Brain
Medulloblastoma
Glioblastoma
Frontal Cortex
Middle Frontal Area
Adenocarcinoma, Endometrial
Top traits
Heel bone mineral density
Systolic blood pressure
Lung function (FEV1/FVC)
Pulse pressure
Coronary artery disease
Intraocular pressure
Height
Waist circumference BMI-adj
QT interval
Prostate cancer
Peak expiratory flow
Red blood cell count
Waist-hip ratio
Total cholesterol levels
Triglycerides
HDL cholesterol levels
Cardiovascular disease
Arm Muscle
LDL cholesterol levels
Reticulocyte count
Waist-to-hip ratio BMI-adj
Educational attainment
Body mass index
Diastolic blood pressure
Cognitive performance
Schizophrenia
Intelligence (MTAG)
Type 2 diabetes
General cognitive ability
Neuroticism
Reaction time
A
Epimap
Loci-tissue-trait
metabolic related
traits: Blood-related, inflammation
tissues: immune related
openord layout
C
mean bridging coefficient
random
M1 (top: brain, CNS)
M2 (top: hear, CV)
M3 (top: lipid-related)
M4 (top: immune-related)
D
combined topological location
number of genes
proportion of PGEs (>5 genes/total)
M1, M2, M3
M1
M2
M3
Random triplets
E
metabo-HAR PGEs
combined hub regions (M1, M2, M3)
Regions within selected compartment
* Top driver regions
position
chromosomes

**Fig. S15. EpiMap network integration and characterization of lipid-traits relationships** *(related to Fig. 2)*. (**A**) Epimap network reconstruction of loci with tissue and trait enrichments. Module classification in openord layout displaying metabolic trait relationships in interconnected modules (module 1-3) and immune-related traits. (**B**) Trait and tissue top degree scores in each module with strong metabolic relationships. (**C**) Mean bridging coefficient across modules and random network. (**D**) Combined topological coalescence of positional enrichments for top genes within each interconnected metabolic trait, followed by chromosomal landscape positioning and colocalization with metabolic-HAR regions, and top driver regions from combinatorial-hub assessment (**E**).

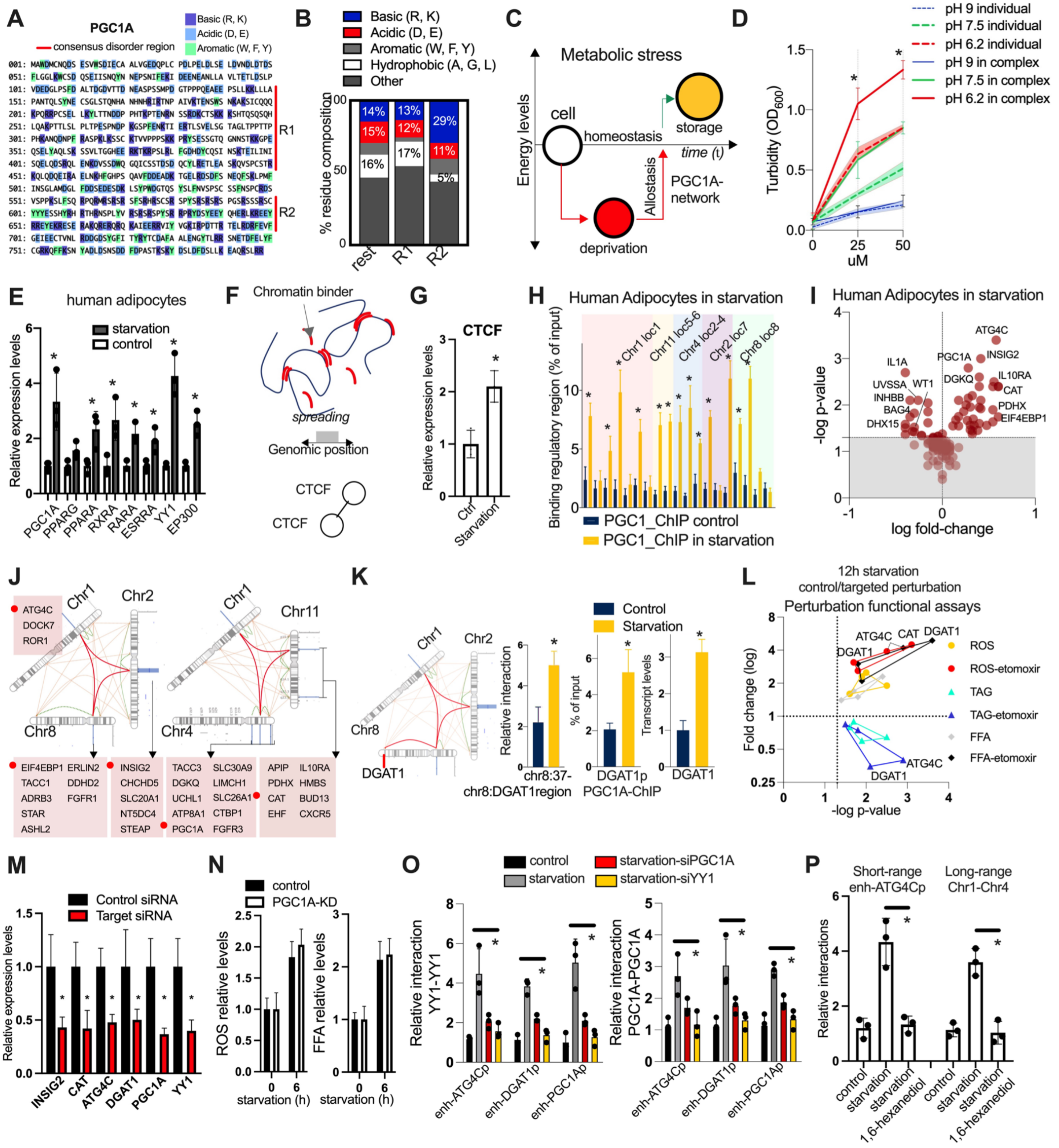

**Fig. S16. Metabolic control of nuclear compartmentalization** *(related to Fig. 3)*. (**A**) PGC1A protein sequence with consensus disorder regions, basic, acidic and aromatic residues. (**B**) Proportion comparison of residue composition features between disorder regions and the rest of PGC1A protein sequence. (**C**) Illustration of homeostatic and allostatic mechanisms regulated by PGC1A-network (**D**) Turbidity assays at different concentrations and pH variations for PGC1A alone or in complex with RARA. (**E**) Expression of transcriptional regulators in glucose starvation (12h). (**F**) Illustration showing genomic spreading between chromatin interactions mediated by structuring candidates such as CTCF. (**G**) CTCF expression as in (E). (**H**) ChIP-qPCR of PGC1A to HAR elements within selected metabolic-HAR compartments. (**I**) Expression of metabolic-HAR genes within selected compartments during glucose starvation. Top regulated genes in red and compartments in panel (**J**). (**K**) Chromosomes 1-2-8 compartment and DGAT1 loci location, CTCF-mediated long range homotypic interaction in chr8, PGC1A binding to DGAT1 promoters and DGAT1 expression levels. All assays were done as in (E). (**L**) Functional assays in starvation with or without CPT1A-channel inhibitor etomoxir, in cells perturbed by siRNA for top regulated genes in metabolic-HAR regions. Etomoxir exacerbates targeted perturbation phenotypes. (**M**) Gene expression after siRNA knockdowns. (**N**) Functional assays in control and PGC1A knockdown cells after 6h of starvation induction. (**O**) Relative short-range interactions in starvation and in targeted siRNA downregulation for top regulated metabolic-HAR genes. (**P**) YY1-short- and CTCF-long-range mediated interactions in starvation and abolished by 1,6-hexanediol. Bars show mean values and error bars indicate SEM. Unpaired, two-tailed student's t-test was used when two groups were compared, and ANOVA followed by fisher's least significant difference (LSD) test for post hoc comparisons for multiple groups. * p-va<0.05

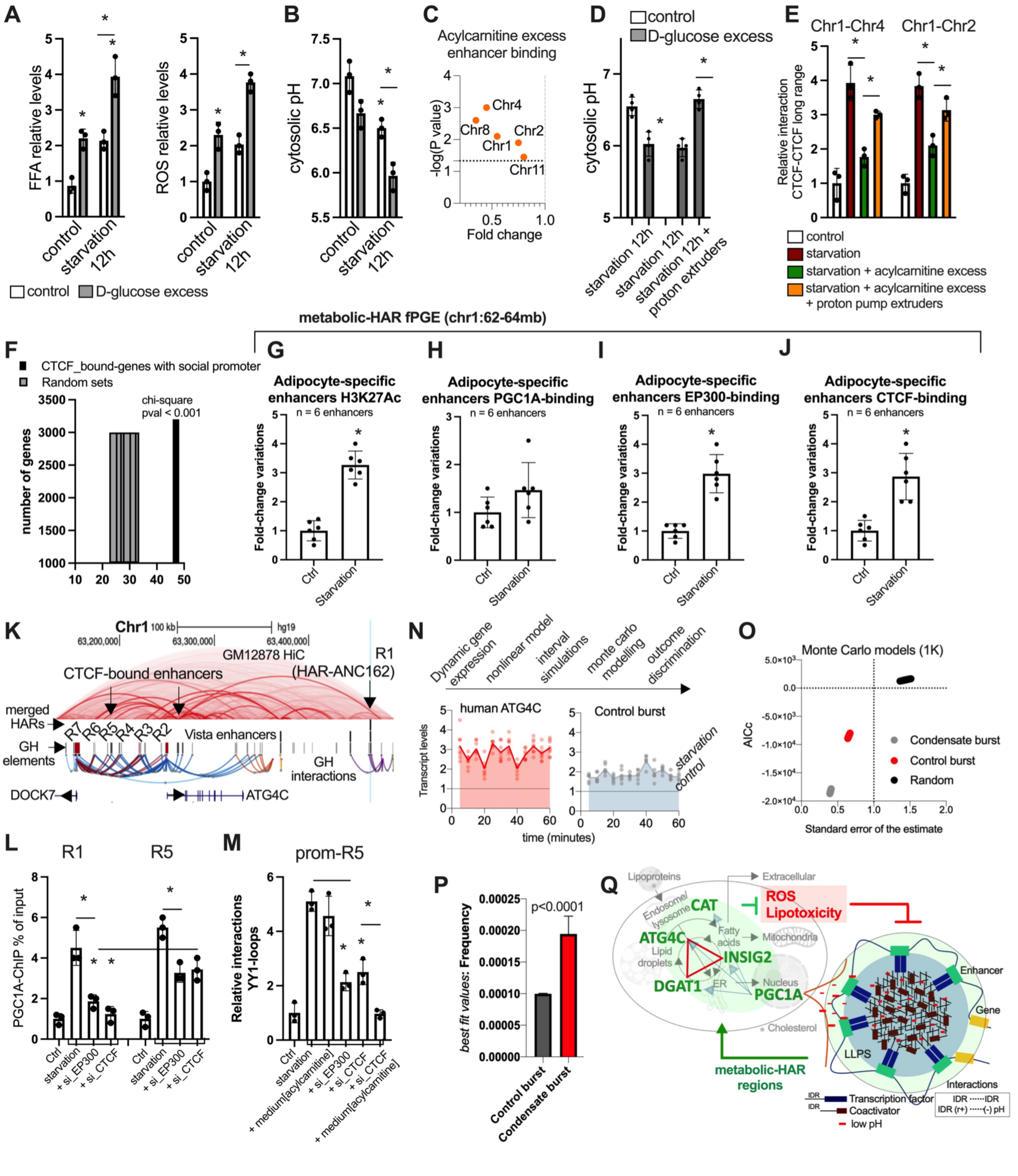


**Fig. S17. Metabolic control of nuclear compartmentalization** *(related to Fig. 3)*. (**A**) FFA and ROS functional assays in human adipocytes in starvation and with glycolysis inhibition by D-glucose. (**B**) Cytosolic pH in the same conditions as in A. (**C**) PGC1A binding to selected HAR-elements during starvation and with acylcarnitine excess (20uM). (**D**) Cytosolic pH in starvation and after glycolysis inhibition and activators of proton pump extruders such as Zinc. (**E**) CTCF long range chromatin interactions in starvation, starvation with acylcarnitine excess as in C, and proton pump activators as in D. (**F**) Proportion of CTCF-bound genes displaying social promoters from our regulatory regions interaction network, compared to random permutations. (**G**) H3K27Ac ChIP-qPCR in adipocyte specific enhancers within selected metabolic HAR region during starvation (12h). (**H**) PGC1A-binding by ChIP-qPCR in adipocyte specific enhancers as in G. (**I**) EP300 ChIP-qPCR as in G. (**J**) CTCF-binding by ChIP-qPCR as in G. (**K**) Human genome track showing zoom in top target within metabolic-HAR domain in chr1:62-64mb: ncHARs (ANC162), regulatory elements and interactions, GM12878-Hi-C data, CTCF-bound elements and promoter locations of ATG4C and DOCK7. (**L**) Exacerbated loss of PGC1A binding to HAR by EP300 and CTCF knockdown. PGC1A ChIP-qPCR during starvation and with EP300 and CTCF knockdown in R1(HAR) element and R5 (CTCF bound) element. (**M**) Loss of resilience and sensitization to metabolic overload in structural loops by EP300 and CTCF knockdown. YY1-mediated loop interactions between ATG4C-promoter and R5-CTCF bound element in starvation, in medium acylcarnitine excess (10uM), with EP300 and CTCF knockdowns and together with medium AC excess. (**N**) Pipeline steps description of Monte Carlo modeling of dynamic gene expression data in stimulated conditions (starvation). Panels below are dynamic transcript levels in starvation of regulated genes: ATG4C gene (social promoter by regulatory interaction networks) and ANGPTL3 (isolated promoter). (**O**) Akaike information criterion and standard error of the estimate for Monte Carlo burst models from dynamic gene expression comparing genes in N and random models. (**P**) Frequency best-fit-values parameters from Monte Carlo discrimination of burst models. (**Q**) Illustration of mechanistic insight for lipid cycling regulation of nuclear compartment formation. Top genes within interacting metabolic-HAR domains regulated by PGC1A-network during starvation and controlling lipid-derived oxidative stress and toxicity. Bars show mean values and error bars indicate SEM. Unpaired, two-tailed student's t-test was used when two groups were compared, and ANOVA followed by fisher's least significant difference (LSD) test for post hoc comparisons for multiple groups. * p-value <0.05.

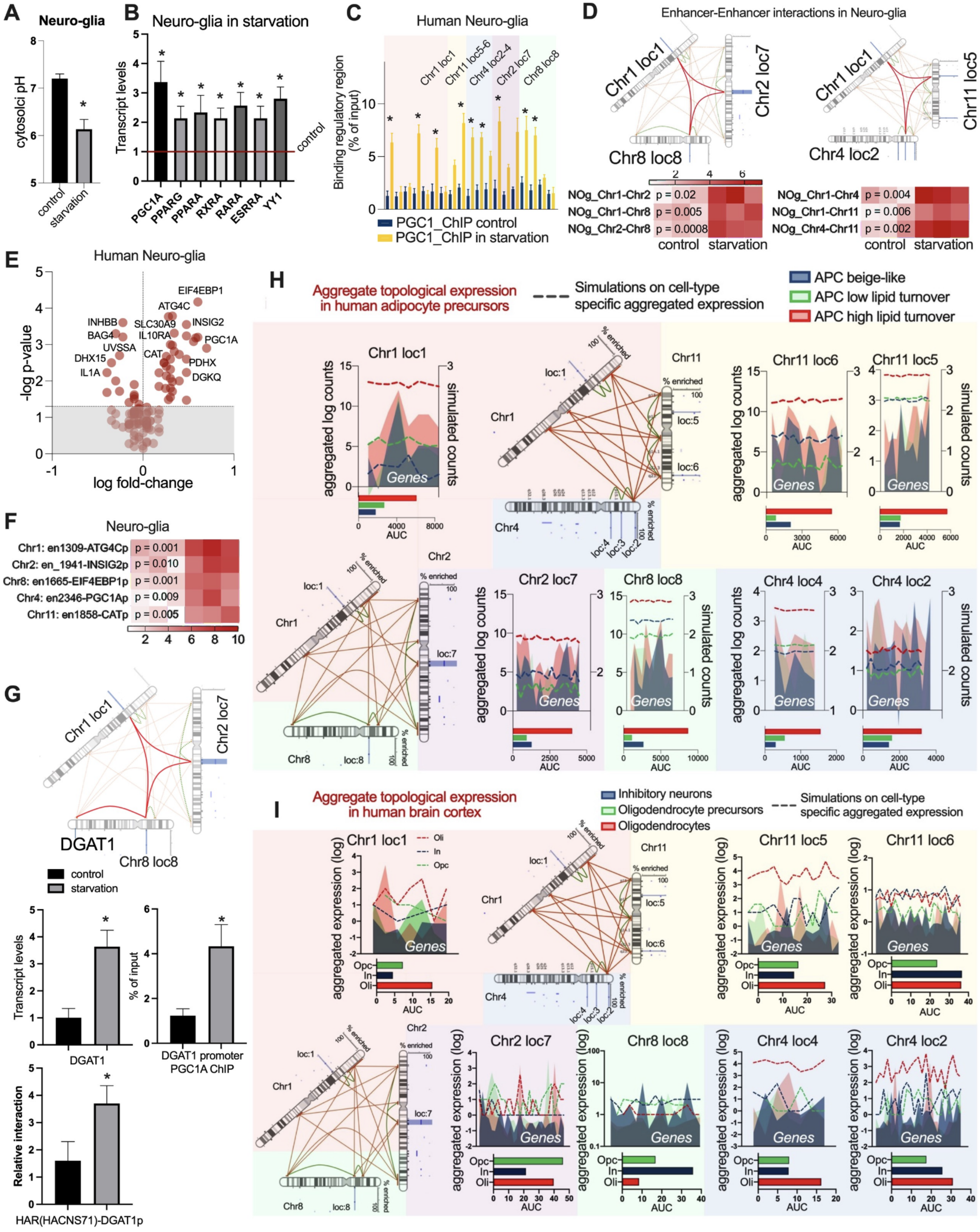


**Fig. S18. Metabolic control of nuclear compartmentalization in APCs and brain cells** *(related to Fig. 3)*. (**A**) Cytosolic pHin starvation in human neuro-glia cultures.. (**B**) Expression of PGC1A-network of transcriptional regulators during starvation (12h) in human neuro-glial cell cultures. (**C**) PGC1A binding by ChIP-qPCR to HAR elements in selected metabolicHAR domains. (**D**) ChIp-loop of metabolic-HAR interactions within selected nuclear compartments.(**E**) Expression of metabolic-HAR genes within selected compartments during glucose starvation. (**F**) Enhancer-promoter loops between HAR-elements and starvation-regulated genes within metabolic-HAR domain. (**G**) HAR-DGAT1 CTCF-mediated long-range interaction by ChIP-loop, DGAT1 expression levels, PGC1A-binding to DGAT1 promoter and CTCF-mediated short range HAR-promoter interactions. (**H**) Aggregate domain expression of genes composing metabolic-HAR domains in human-derived adipose precursor cells subtypes. (**I**) Aggregate domain expression of genes composing metabolic-HAR domains in human-derived single nuclei RNA seq brain cell subtypes. Bars show mean values and error bars indicate SEM. Unpaired, two-tailed student's t-test was used when two groups were compared, and ANOVA followed by fisher's least significant difference (LSD) test for post hoc comparisons for multiple groups. * p-value <0.05.

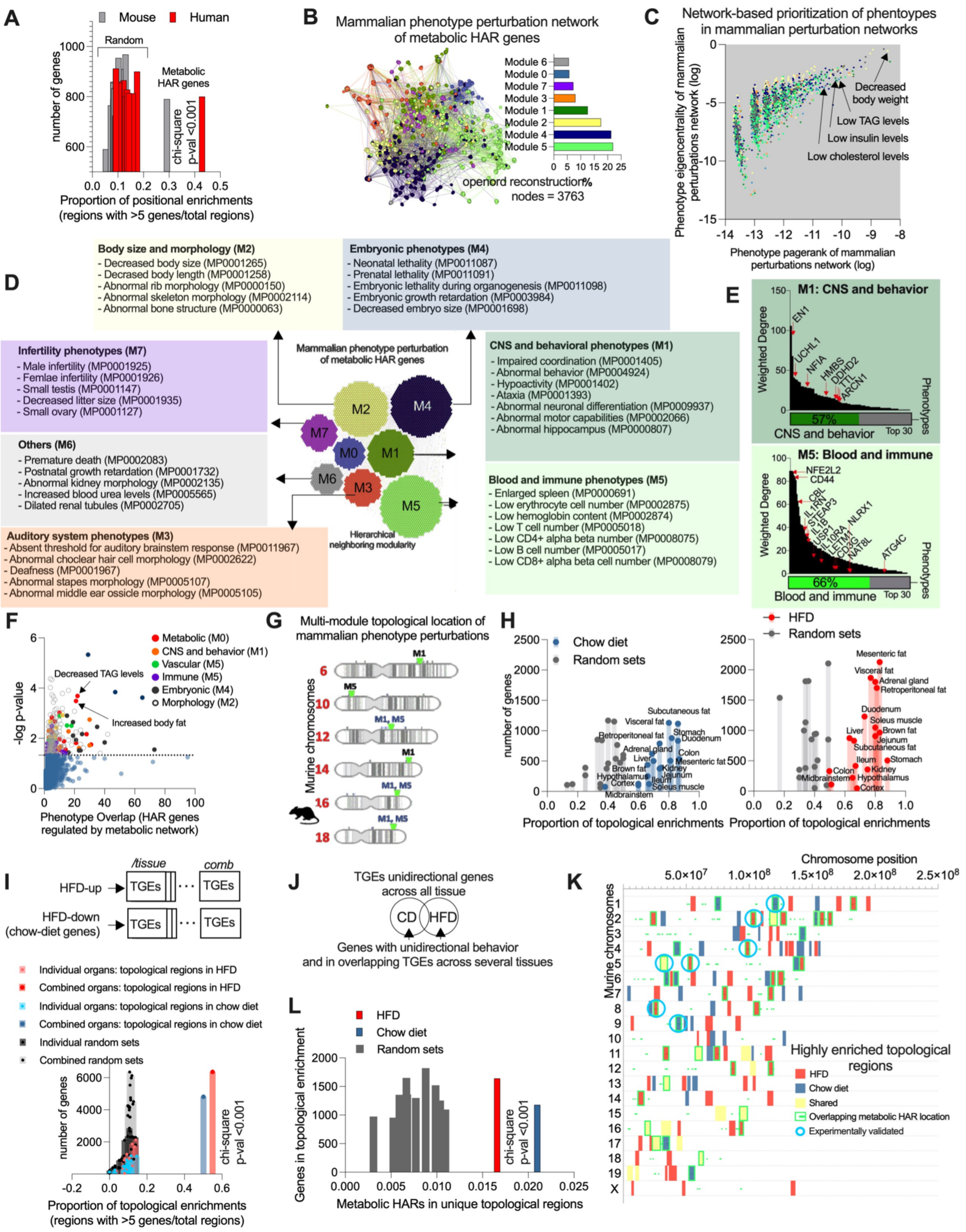
A
Mouse
Human
Random
Metabolic HAR genes
chi-square p-val <0.001
number of genes
Proportion of positional enrichments (regions with >5 genes/total regions)
B
Mammalian phenotype perturbation network of metabolic HAR genes
Module 6
Module 0
Module 7
Module 3
Module 1
Module 2
Module 4
Module 5
openord reconstruction
%
nodes = 3763
C
Network-based prioritization of phentoypes in mammalian perturbation networks
Decreased body weight
Low TAG levels
Low insulin levels
Low cholesterol levels
Phenotype eigencentrality of mammalian perturbations network (log)
Phenotype pagerank of mammalian perturbations network (log)
D
Body size and morphology (M2)
- Decreased body size (MP0001265)
- Decrased body length (MP0001258)
- Abnormal rib morphology (MP0000150)
- Abnormal skeleton morphology (MP0002114)
- Abnormal bone structure (MP0000063)
Embryonic phenotypes (M4)
- Neonatal lethality (MP0011087)
- Prenatal lethality (MP0011091)
- Embryonic lethality during organogenesis (MP0011098)
- Embryonic growth retardation (MP0003984)
- Decreased embryo size (MP0001698)
Infertility phenotypes (M7)
- Male infertility (MP0001925)
- Femlae infertility (MP0001926)
- Small testis (MP0001147)
- Decreased litter size (MP0001935)
- Small ovary (MP0001127)
Mammalian phenotype perturbation of metabolic HAR genes
M2
M4
M7
M0
M1
M6
M3
M5
Hierarchical neighboring modularity
CNS and behavioral phenotypes (M1)
- Impaired coordination (MP0001405)
- Abnormal behavior (MP0004924)
- Hypoactivity (MP0001402)
- Ataxia (MP0001393)
- Abnormal neuronal differentiation (MP0009937)
- Abnormal motor capabilities (MP0002066)
- Abnormal hippocampus (MP0000807)
Others (M6)
- Premature death (MP0002083)
- Postnatal growth retardation (MP0001732)
- Abnormal kidney morphology (MP0002135)
- Increased blood urea levels (MP0005565)
- Dilated renal tubules (MP0002705)
Blood and immune phenotypes (M5)
- Enlarged spleen (MP0000691)
- Low erythrocyte cell number (MP0002875)
- Low hemoglobin content (MP0002874)
- Low T cell number (MP0005018)
- Low CD4+ alpha beta number (MP0008075)
- Low B cell number (MP0005017)
- Low CD8+ alpha beta cell number (MP0008079)
Auditory system phenotypes (M3)
- Absent threshold for auditory brainstem response (MP0011967)
- Abnormal choclear hair cell morphology (MP0002622)
- Deafness (MP0001967)
- Abnormal stapes morphology (MP0005107)
- Abnormal middle ear ossicle morphology (MP0005105)
E
M1: CNS and behavior
Weighted Degree
EN1
UCHL1
NFIA
HMBS
DDHD2
TTL
ARCN1
57%
CNS and behavior
Top 30
Phenotypes
M5: Blood and immune
NFE2L2
CD44
CBL
IL1RN
STEAP3
IL1B
USP1
IL10RA
LETM1
CD3G
NAT8L
NLRX1
ATG4C
66%
Blood and immune
F
Metabolic (M0)
CNS and behavior (M1)
Vascular (M5)
Immune (M5)
Embryonic (M4)
Morphology (M2)
Decreased TAG levels
Increased body fat
-log p-value
Phenotype Overlap (HAR genes regulated by metabolic network)
G
Multi-module topological location of mammalian phenotype perturbations
Murine chromosomes
M1
M5
M1, M5
H
Chow diet
Random sets
HFD
number of genes
Subcutaneous fat
Visceral fat
Retroperitoneal fat
Stomach
Duodenum
Adrenal gland
Colon
Liver
Mesenteric fat
Brown fat
Kidney
Hypothalamus
Jejunum
Ileum
Cortex
Soleus muscle
Midbrainstem
Proportion of topological enrichments
I
/tissue
comb
HFD-up
TGEs
HFD-down (chow-diet genes)
Individual organs: topological regions in HFD
Combined organs: topological regions in HFD
Individual organs: topological regions in chow diet
Combined organs: topological regions in chow diet
Individual random sets
Combined random sets
chi-square p-val <0.001
Proportion of topological enrichments (regions with >5 genes/total regions)
J
TGEs unidirectional genes across all tissue
CD
HFD
Genes with unidirectional behavior and in overlapping TGEs across several tissues
L
HFD
Chow diet
Random sets
Genes in topological enrichment
chi-square p-val <0.001
Metabolic HARs in unique topological regions
K
Chromosome position
Murine chromosomes
Highly enriched topological regions
HFD
Chow diet
Shared
Overlapping metabolic HAR location
Experimentally validated

**Fig. S19. Mammalian convergence and tuning of spatial genome architecture** *(related to Fig. 4)*. (**A**) Proportion of positional gene enrichments (>5 genes per genomic cluster) for metabolic-HAR genes in humans, mice, and random gene sets. (**B**) Network integration of mammalian genetic perturbations for metabolic-HAR genes in openord layout and modularity (right panel). (**C**) Network-based prioritization of mammalian perturbation relationships for metabolic-HAR genes showing top phenotypes. (**D**) Top phenotypes in each module.(**E**) Node degree classification of gene drivers per phenotype in CNS and blood-immune modules. (**F**) Phenotype enrichment for metabolic-HAR genes showing preservation of network modules and metabolic phenotype as top enrichments. (**G**) Multi-module topological convergence of metabolic interacting modules in murine chromosome regions. (**H**) Mammalian HFD perturbations followed by RNAseq in 19 tissues and spatial-PGEs analysis. Differentially expressed genes for both chow diet and HFD were used for hub-positional enrichments compared to random sets. (**I**) Combinatorial hub-positional enrichments for diet-induced genes comparing organ-effect and cumulative hub positions across all organs assessed. (**J**) Venn diagram illustration of topological positional enrichments (TGE or PGE) for differential genes specifically regulated by either diet intervention. (**K**) Chromosomal landscape hub-positions for PGEs regulated by diet intervention, with overlapping syntenic metabolic-HAR regions and those that were experimentally validated. (**L**) Quantification of metabolic-HAR domain overlap with diet induced cumulative hubs.

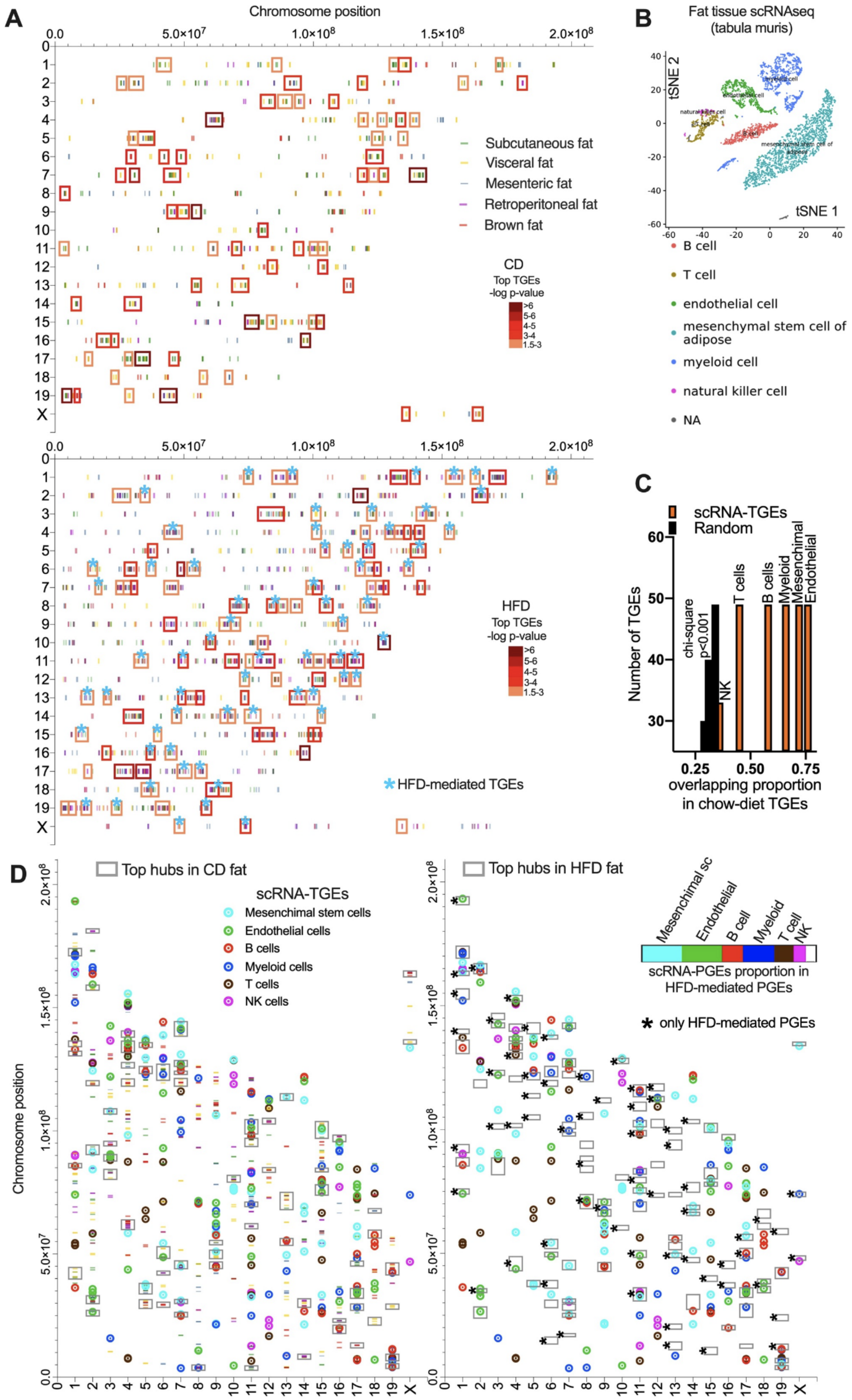


**Fig. S20. Single cell decomposition of spatial transcriptional hubs** *(related to Fig. 4)*. (**A**) Chromosomal landscape of hub-positions for PGEs regulated by diet intervention in different adipose depots. With an asterisk highlighted top positional shifts caused by HFD. (**B**) t-SNE cluster plots for mice adipose tissue scRNA-seq from Taboula Muris consortia. (**C**) Proportion of hub-overlap between scRNAseq transcriptional hubs and chow-diet RNA-seq hubs compared to random hotspots. (**D**) Chromosomal landscape of hub-transcriptional positions for scRNA-hotspots and chow- and HFD mediated hubs. Proportion of overlap between top hubs for HFD and cell type.

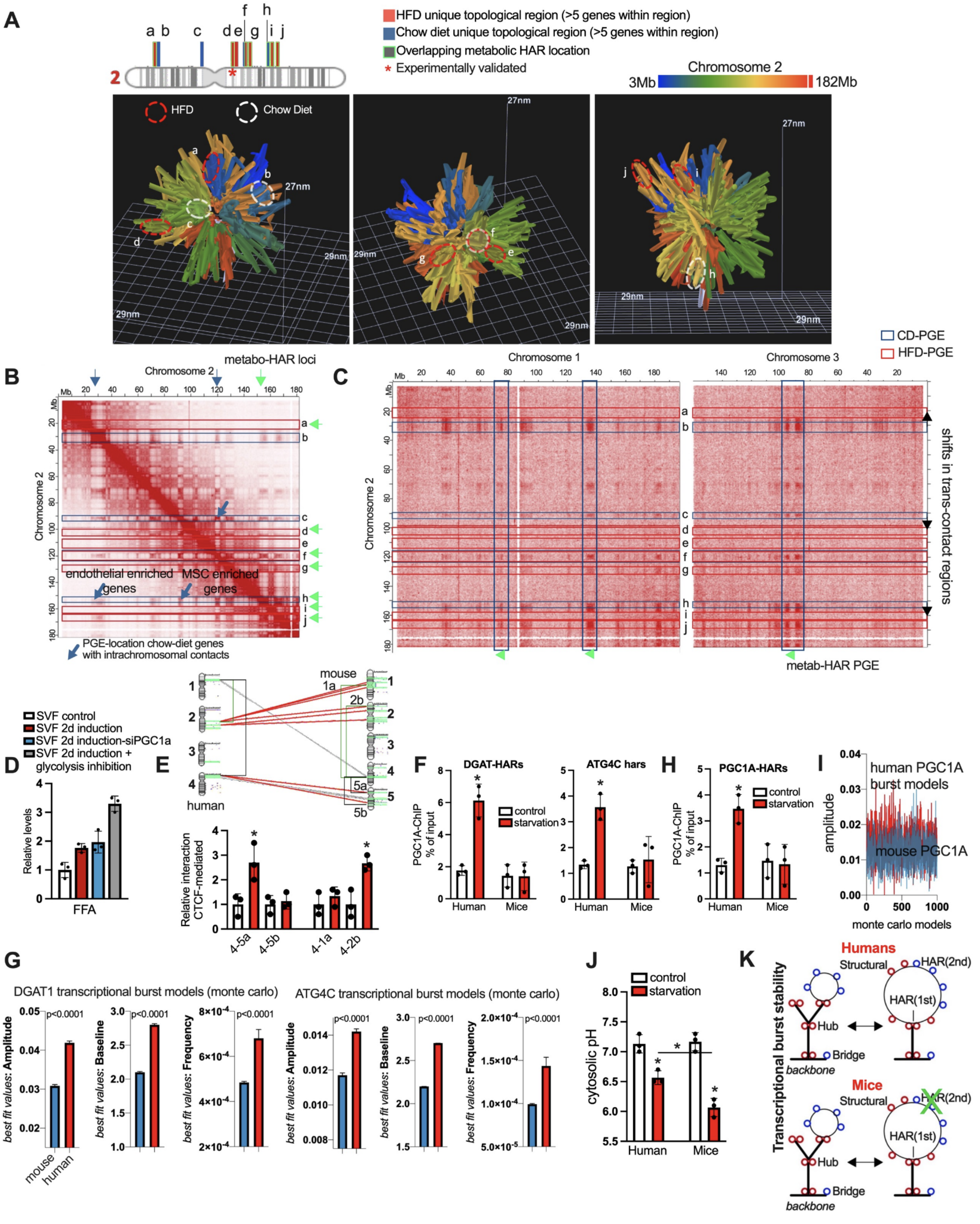


**Fig. S21. Mammalian tuning of spatial genome architecture** *(related to Fig. 4)*. (**A**) Chromosome 2 hotspot regulated by diet and overlapping synteny metabolic-HAR regions. With an asterisk experimentally validated HAR. In lower panels, 3DMax reconstruction of homotypic distances from HiC data displaying diet-specific hotspot and positional shifts caused by HFD. (**B**) Representative HiC contact frequency map for chromosome 2 from mESCs (*132*). Blue squares represent chow-diet specific hotspots while red HFD hubs. Green arrow shows syntenic metabolic-HAR domain locations and blue arrows show single cell hubs located in homotypic interacting chromosome regions, and preferentially enriched in chow-diet hotspots. c-h homotypic interactions with MSCs hotspots and b-h with endothelial hotspots (**C**) Diet-induced shift in hub locations from trans-interacting in chow-diet to neighboring non interacting chromatin regions in HFD. Green arrow shows syntenic metabolic-HAR domain locations. (**D**) Free-fatty acid assay in stromal-vascular fraction cell cultures from mice fat tissue after induction of differentiation with PGC1A perturbation and glycolysis inhibition. (**E**) Top panel shows 3 experimentally validated metabolic-HAR regions in humans, connected to their syntenic mice domains in grey if it is fully preserved, and in red if it is split in mice. Lower panel shows CTCF-mediated long-range interactions for nuclear compartment in mice cell cultures during starvation. (**F**) PGC1A-binding by ChIP-qPCR in selected HAR elements and in mouse orthologue positions for top metabolic-HAR regions ATG4C and DGAT1. (**G**) Monte Carlo burst models from dynamic gene expression in starvation for selected genes DGAT and ATG4C from mice and human cell culture systems. Showing best-fit-values from discriminatory outcomes for amplitude, baseline and frequency. (**H**) PGC1A-binding by ChIP-qPCR in selected HAR elements and in mouse orthologue positions for top metabolic-HAR regions PGC1A domain. (**I**) Monte Carlo burst models from dynamic gene expression in starvation for selected gene PGC1A from mice and human cell culture systems. Showing best-fit-values from discriminatory outcomes for amplitude. (**J**) Functional assays for cytosolic pH in cell cultures during starvation at 12h. (**K**) Illustration showing nested model transcriptional variations between human and mice, and the transcriptional optimization and stability offered by cis-HAR anchoring of chromatin conformations. Bars show mean values and error bars indicate SEM. Unpaired, two-tailed student's t-test was used when two groups were compared, and ANOVA followed by fisher's least significant difference (LSD) test for post hoc comparisons for multiple groups. * p-value <0.05.

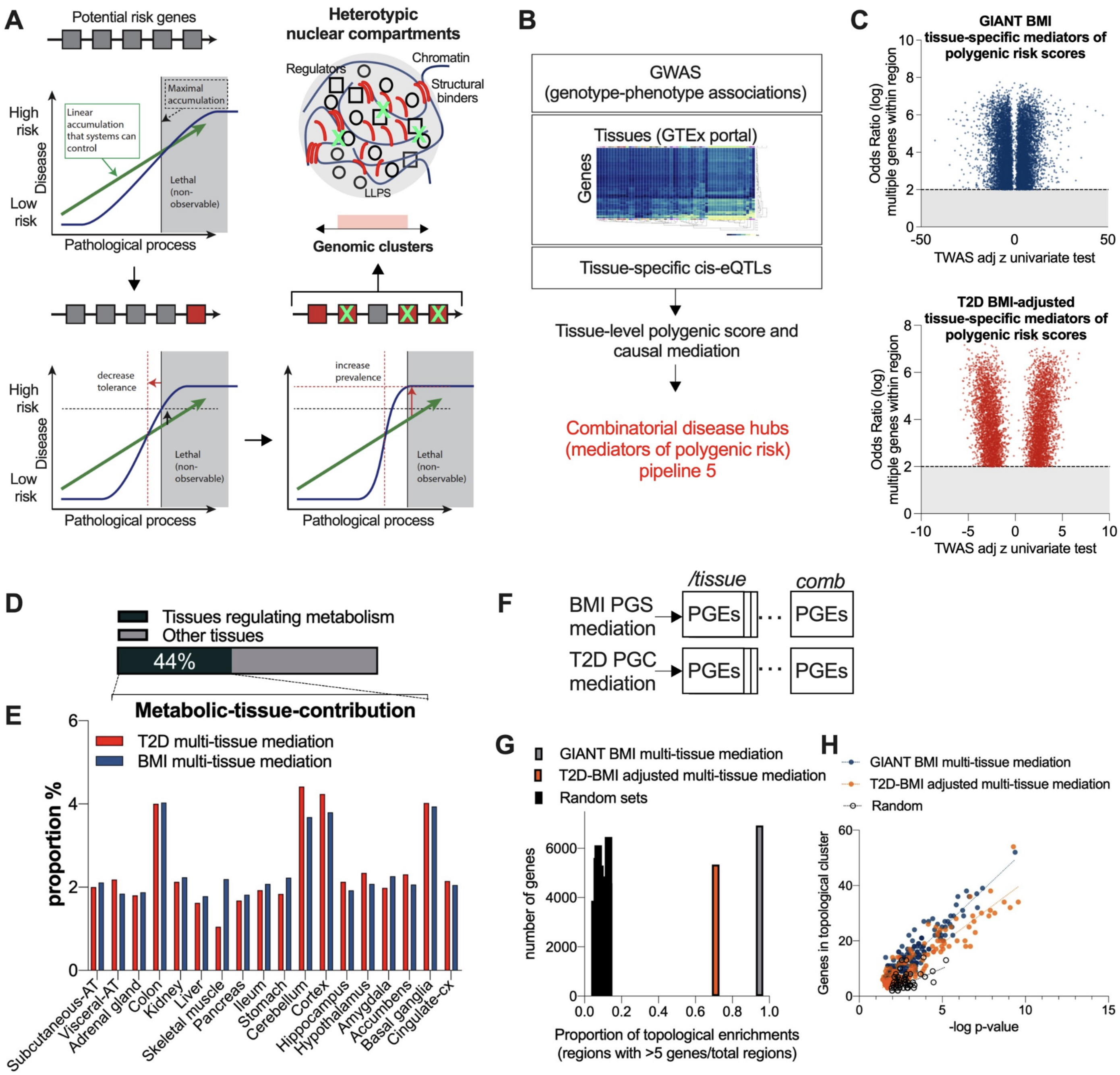


**Fig. S22. Genomic clusters of mediators of polygenic risk for metabolic disease** *(related to Fig. 5)*. (**A**) Polygenic risk model illustration showing the cumulative effect of genetic variation on disease risk prediction. Here we exploit the cumulative effect of related variations and genes affected thereby, to find genomic risk clusters of co-regulated transcriptional compartments. (**B**) Illustration of mediation pipeline that integrates GWAS-association with tissue-specific cis-eQTLs and tissue-specific gene expression profiles from GTEx portal. Causal mediators of trait-risk are used as input for integrative topological genomic pipelines described in this paper. (**C**) Volcano plots of tissue-specific mediator genes of polygenic risk for BMI and T2D disease risk. (**D**) Proportion of mediator enrichments in metabolic tissues. (**E**) Metabolic tissue contribution to polygenicity for T2D and BMI traits. (**F**) Illustration showing individual and combinatorial positional hubs for gene mediators of polygenic risk for T2D and BMI. (**G**) Proportion of genomic clusters with > 5 genes per hotspot from combined topological enrichment for T2D and BMI mediators. (**A**) Positional hub enrichment p-values and number of genes composing each cluster.

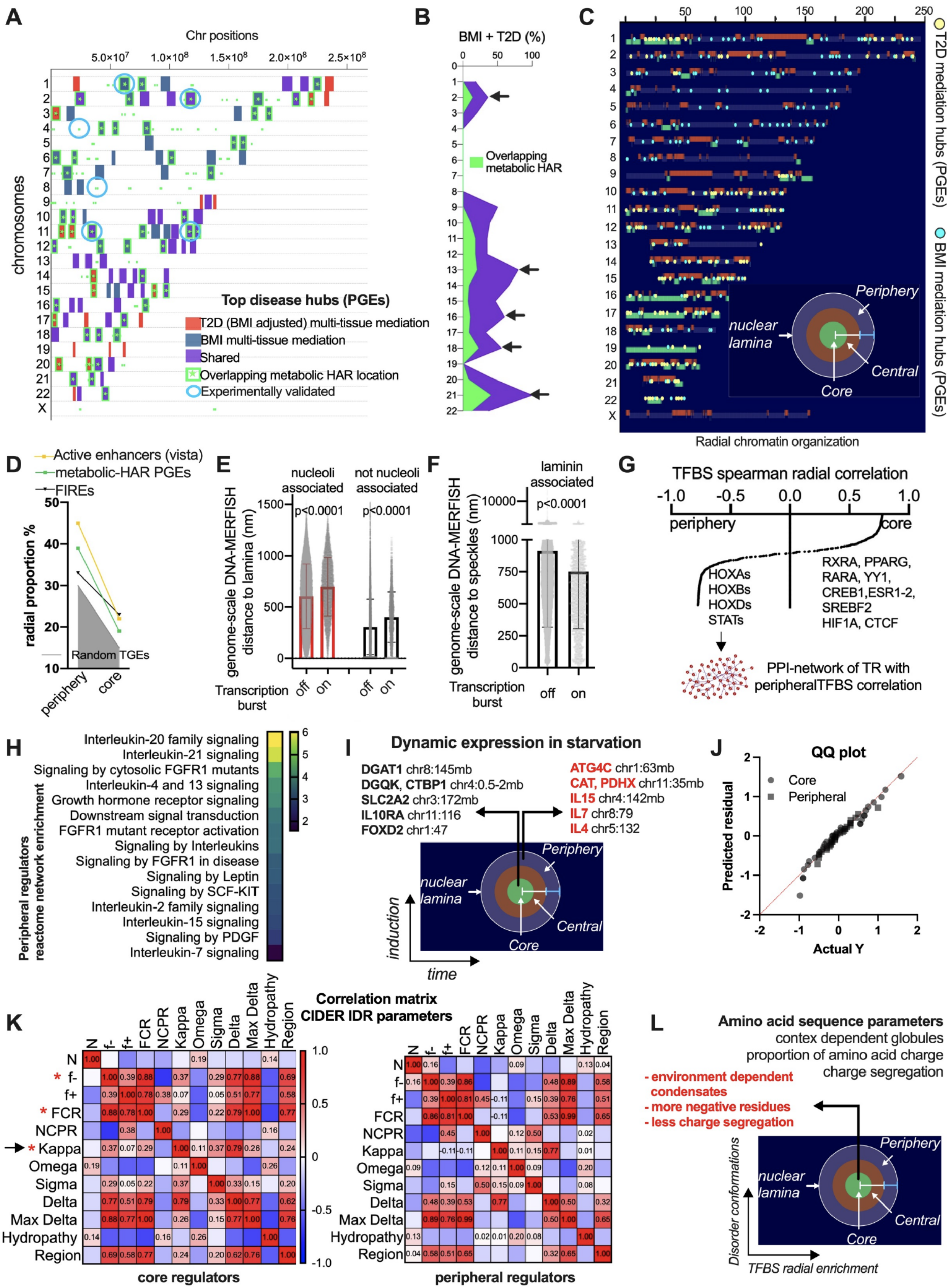
A
Chr positions
chromosomes
Top disease hubs (PGEs)
T2D (BMI adjusted) multi-tissue mediation
BMI multi-tissue mediation
Shared
Overlapping metabolic HAR location
Experimentally validated
B
BMI + T2D (%)
Overlapping metabolic HAR
C
T2D mediation hubs (PGEs)
BMI mediation hubs (PGEs)
Periphery
nuclear lamina
Central
Core
Radial chromatin organization
D
Active enhancers (vista)
metabolic-HAR PGEs
FIREs
radial proportion %
Random TGEs
periphery
core
E
nucleoli associated
not nucleoli associated
genome-scale DNA-MERFISH distance to lamina (nm)
p<0.0001
Transcription burst
off
on
F
laminin associated
genome-scale DNA-MERFISH distance to speckles (nm)
p<0.0001
Transcription burst
off
on
G
TFBS spearman radial correlation
periphery
core
HOXAs
HOXBs
HOXDs
STATs
RXRA, PPARG, RARA, YY1, CREB1,ESR1-2, SREBF2 HIF1A, CTCF
PPI-network of TR with peripheralTFBS correlation
H
Peripheral regulators reactome network enrichment
Interleukin-20 family signaling
Interleukin-21 signaling
Signaling by cytosolic FGFR1 mutants
Interleukin-4 and 13 signaling
Growth hormone receptor signaling
Downstream signal transduction
FGFR1 mutant receptor activation
Signaling by Interleukins
Signaling by FGFR1 in disease
Signaling by Leptin
Signaling by SCF-KIT
Interleukin-2 family signaling
Interleukin-15 signaling
Signaling by PDGF
Interleukin-7 signaling
I
Dynamic expression in starvation
DGAT1 chr8:145mb
DGQK, CTBP1 chr4:0.5-2mb
SLC2A2 chr3:172mb
IL10RA chr11:116
FOXD2 chr1:47
ATG4C chr1:63mb
CAT, PDHX chr11:35mb
IL15 chr4:142mb
IL7 chr8:79
IL4 chr5:132
induction
time
J
QQ plot
Core
Peripheral
Predicted residual
Actual Y
K
Correlation matrix CIDER IDR parameters
N
f-
f+
FCR
NCPR
Kappa
Omega
Sigma
Delta
Max Delta
Hydropathy
Region
core regulators
peripheral regulators
L
Amino acid sequence parameters
contex dependent globules
proportion of amino acid charge
charge segregation
- environment dependent condensates
- more negative residues
- less charge segregation
Disorder conformations
TFBS radial enrichment

**Fig. S23. Radial divergent nuclear compartmentalization: impact to disease, and transcriptional control** *(related to Fig. 5)*. (**A**) Chromosomal landscape of top disease hubs, trait-specific and shared, as well as overlapping metabolic-HAR regions and experimentally validated domains. (**B**) Chromosomal proportion of shared disease hubs from multi-tissue polygenic scores and causal mediation for BMI and T2D disease variants. (**C**) Radial chromosome organization using GPseq-data (*80*) and PGEs location for disease variant genes from **A**. In different colors radial nuclear location for chromatin loci. (**D**) Radial proportion enrichment for vista active enhancers, metabolic-HAR regions, frequently interacting regions (FIREs) and random gene set derived PGEs. (**E-F**) DNA-MERFISH data (*77*) from transcriptionally active loci (on or off), and their relation to different organelles as well as their distance to nuclear lamina. (**G**) TFBS radial correlation (*80*) with HOX family of transcription factors as top peripherally correlated. Lower panel showing PPI-network of peripherally enriched regulators, and their functional annotation by Reactome (**H**). (**I**) Illustration showing selected genes from metabolic-HAR regions with specific radial enrichments for experimental validation in starvation. (**J**) QQ plot or normality test from experimental validation of dynamic gene expression comparison for radial enriched loci, peripheral vs core locations. (**K**) CIDER Correlation matrix parameters based on amino acid sequences (*59*)for ensemble disorder features from peripheral and core enriched regulators. With an asterisk, charged-mediated variations, and with an arrow segregation of charge within disorder regions. (**L**) Illustration of top parameter variation for core-enriched transcriptional regulators. Bars show mean values and error bars indicate SD. Unpaired, two-tailed student's t-test was used when two groups were compared, and ANOVA followed by fisher's least significant difference (LSD) test for post hoc comparisons for multiple groups.

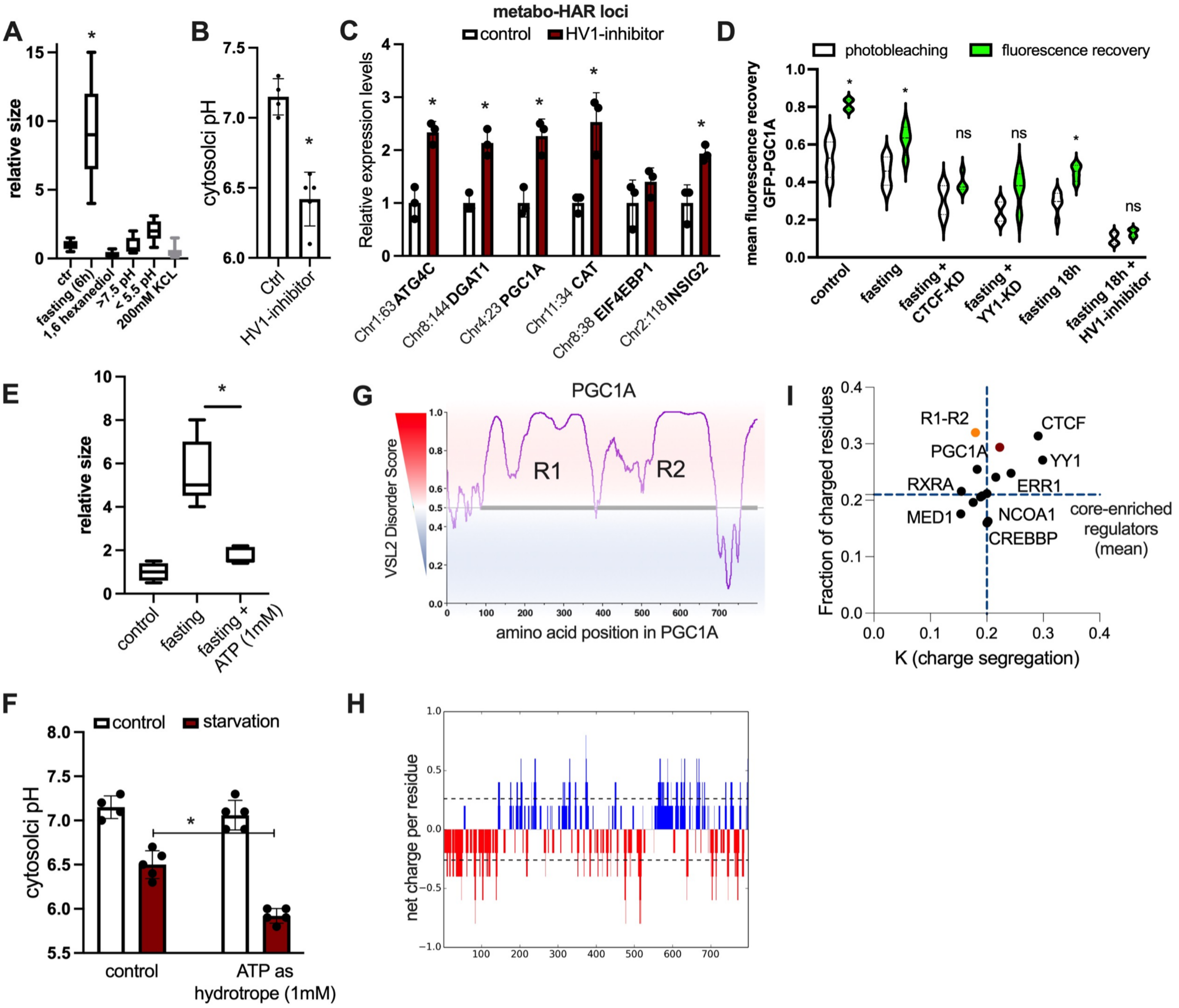

**Fig. S24. Transition from liquid-like to gel-like condensates controls fasting endurance** *(related to Fig. 5)*. (**A**) GFP-PGC1A transcriptional condensate size in microscopy of living cells, after starvation and blocked by 1,6 hexanediol, extreme pH and KCL variations. (**A**) Cytosolic pH variation after proton pump HV1 inhibitor treatment. (**C**) Expression of top regulated genes within metabolic-HAR regions after HV1 treatment. (**C**) Fluorescence recovery mean-variation after refractory period of 6 seconds once bleaching was induced. Comparison in human cells in starvation and with transcriptional regulator perturbations as well as extreme starvation and Hv! Co-treatments. (**E**) GFP-PGC1A transcriptional condensates relative size in living cells microscopy after starvation and blocked by ATP excess. (**F**) Cytosolic pH variation after fasting and ATP excess. (**G**) PONDR VSL2 disorder score of PGC1A amino acid sequence showing the R1and R2 consensus disorder regions.(**H**) Net charge per residue parameter for PGC1A sequence. (**I**) Fraction of charge residues and charge segregation across full length PGC1A, PGC1A disorder regions R1 and R2, PGC1A-interacting regulators and mean nuclear core-enriched transcriptional regulators. Bars show mean values and error bars indicate SEM. Unpaired, two-tailed student's t-test was used when two groups were compared, and ANOVA followed by fisher's least significant difference (LSD) test for post hoc comparisons for multiple groups. * p-value <0.05.

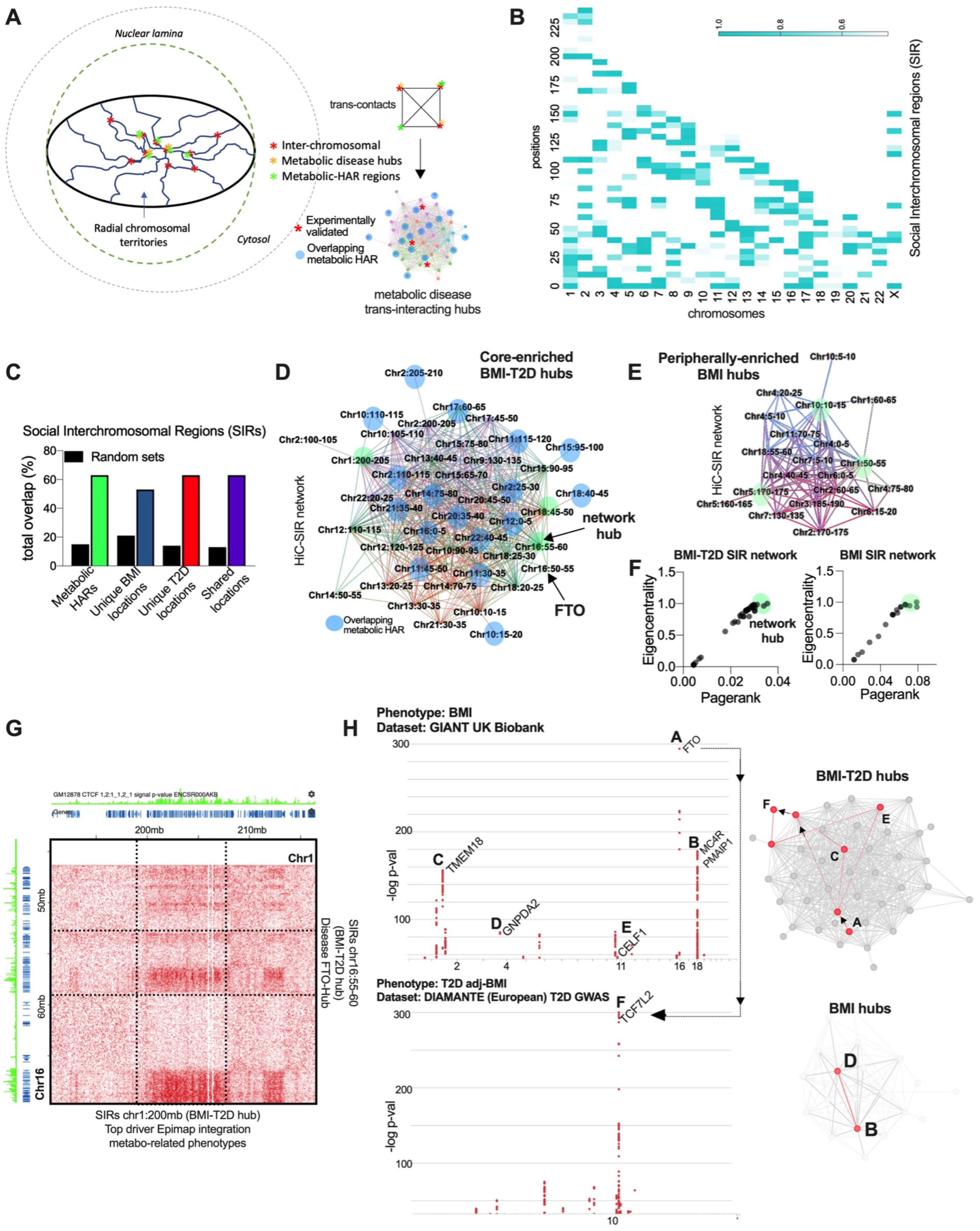
A
Nuclear lamina
Radial chromosomal territories
Cytosol
trans-contacts
Inter-chromosomal
Metabolic disease hubs
Metabolic-HAR regions
Experimentally validated
Overlapping metabolic HAR
metabolic disease trans-interacting hubs
B
positions
chromosomes
Social Interchromosomal regions (SIR)
C
Social Interchromosomal Regions (SIRs)
Random sets
total overlap (%)
Metabolic HARs
Unique BMI locations
Unique T2D locations
Shared locations
D
Core-enriched BMI-T2D hubs
HiC-SIR network
network hub
FTO
Overlapping metabolic HAR
E
Peripherally-enriched BMI hubs
HiC-SIR network
F
BMI-T2D SIR network
BMI SIR network
Eigencentrality
Pagerank
network hub
G
200mb
210mb
Chr1
Chr16
50mb
60mb
SIRs chr16:55-60 (BMI-T2D hub) Disease FTO-Hub
SIRs chr1:200mb (BMI-T2D hub)
Top driver Epimap integration
metabo-related phenotypes
H
Phenotype: BMI
Dataset: GIANT UK Biobank
-log p-val
FTO
TMEM18
GNPDA2
CELF1
MC4R
PMAIP1
Phenotype: T2D adj-BMI
Dataset: DIAMANTE (European) T2D GWAS
TCF7L2
BMI-T2D hubs
BMI hubs

**Fig. S25. Topological systems genetics** *(related to Fig. 6)*. (**A**) Illustration showing genome organization and clusters of co-localization between interacting chromosomal regions, metabolic disease hubs, and metabolic-HAR domains. In the right panel representations of loci-loci interactions between trans-domains and related metabolic-HAR regions. (**B**) Chromosomal landscape displaying social interchromosomal regions (SIR), used to define trans-domains for co-localization with disease hubs and HAR domains. (**C**) Proportion of colocalization between social interchromosomal regions, disease hubs and metabolic-HAR domains. (**D-E**) SIR Network reconstruction of shared T2D-BMI top hubs, unique BMI hubs, and metabolic-HAR domains. (**F**) Network-based analysis of SIR-network. In green showing top hubs in D and E. (**G**) Representative conservation of trans-interacting domains in HiC contact frequency maps (GM12878) for disease hub mediators. Displaying interaction between top hub in T2D-BMI SIR network FTO domain and top driver region in EpiMap integration for metabolic-related phenotypes. (**H**) Intra- and inter-trait relationships among conventional GWAS association studies. Top GWAS with their associated genes and their topological relationships in SIR-disease hub networks.

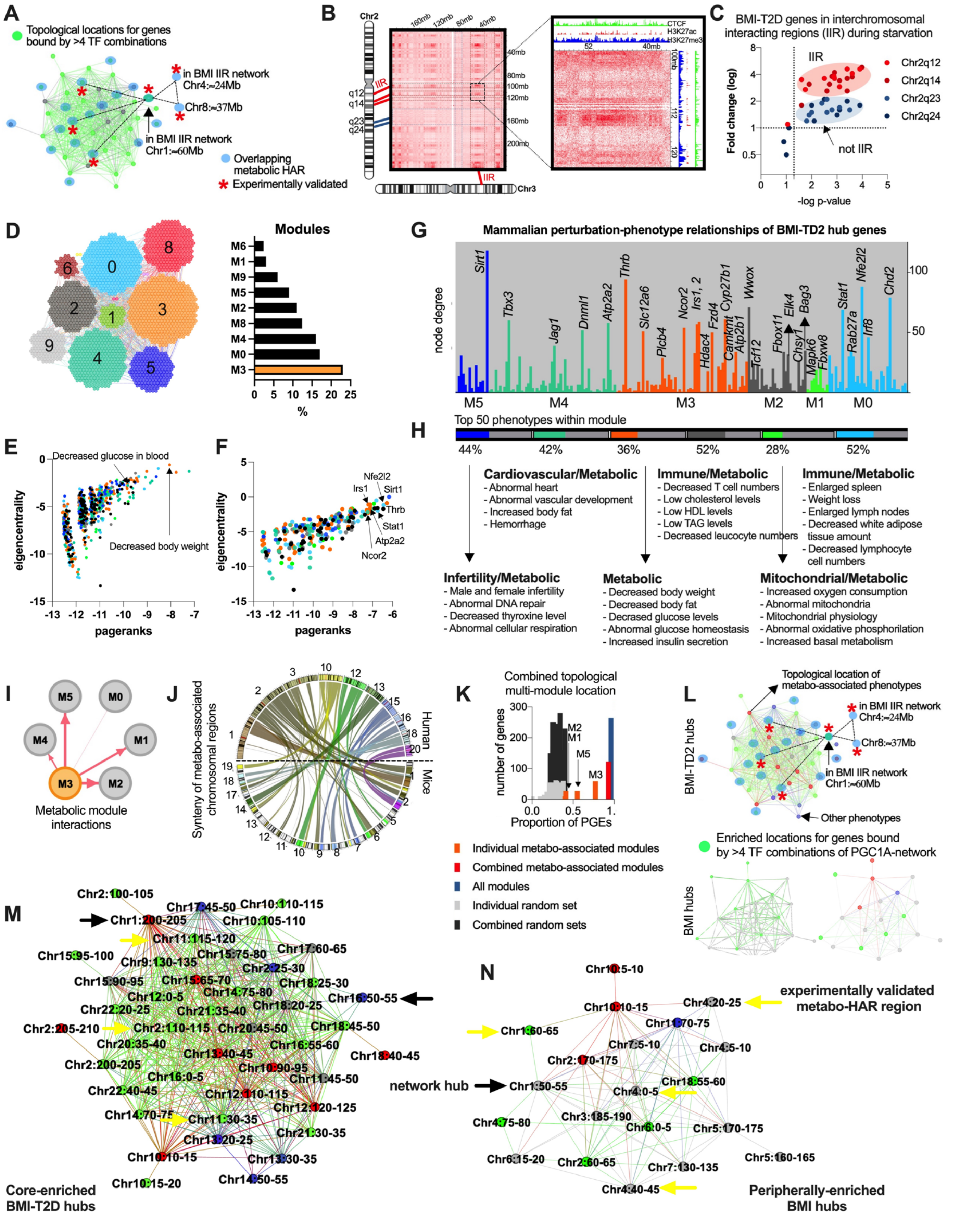


**Fig. S26. Topological systems genetics and integration of mammalian syntenic domains with metabolic phenotype associations** *(related to Fig. 6)*. (**A**) SIR-disease hub network, highlighting in green location of disease-hub genes regulated by combination of PGC1A-interacting regulators. With an asterisk experimentally validated metabolic-HAR regions. (**B**) Trans-interacting chromatin regions in HiC contact frequency maps, showing SIR-domains, and neighboring isolated regions, both harbouring metabolic-HAr and PGC1A regulated genes. (**C**) Experimental validation for regions in B comparing their fold-change variation in starvation. (**D**) Module layout of network integration of mammalian perturbation phenotypes for metabolic disease hubs, mediator of polygenic risk. In orange, top module with overall enrichment for metabolic phenotypes. (**E-F**) Network-based prioritization of phenotype and gene drivers for D. (**G**) Node degree for gene drivers in each module and phenotype enrichments (**H**). (**I**) Inter-module relationships for metabolic phenotype module M3. (**J**) Syntenic conservation in mice for positional gene enrichment from human disease genes driving top interacting modules. (**K**) Combinatorial transcriptional hub enrichments for genes driving phenotypes in interacting modules and located in syntenic regions. (**L**) Colocalization of syntenic regions displaying metabolic phenotype associations in SIR-network. In green, regions regulated by a combination of PGC1A-interacting regulators. (**M-N**) Full SIR-network showing loci with phenotype associations and experimentally validated metabolic-HAR regions.

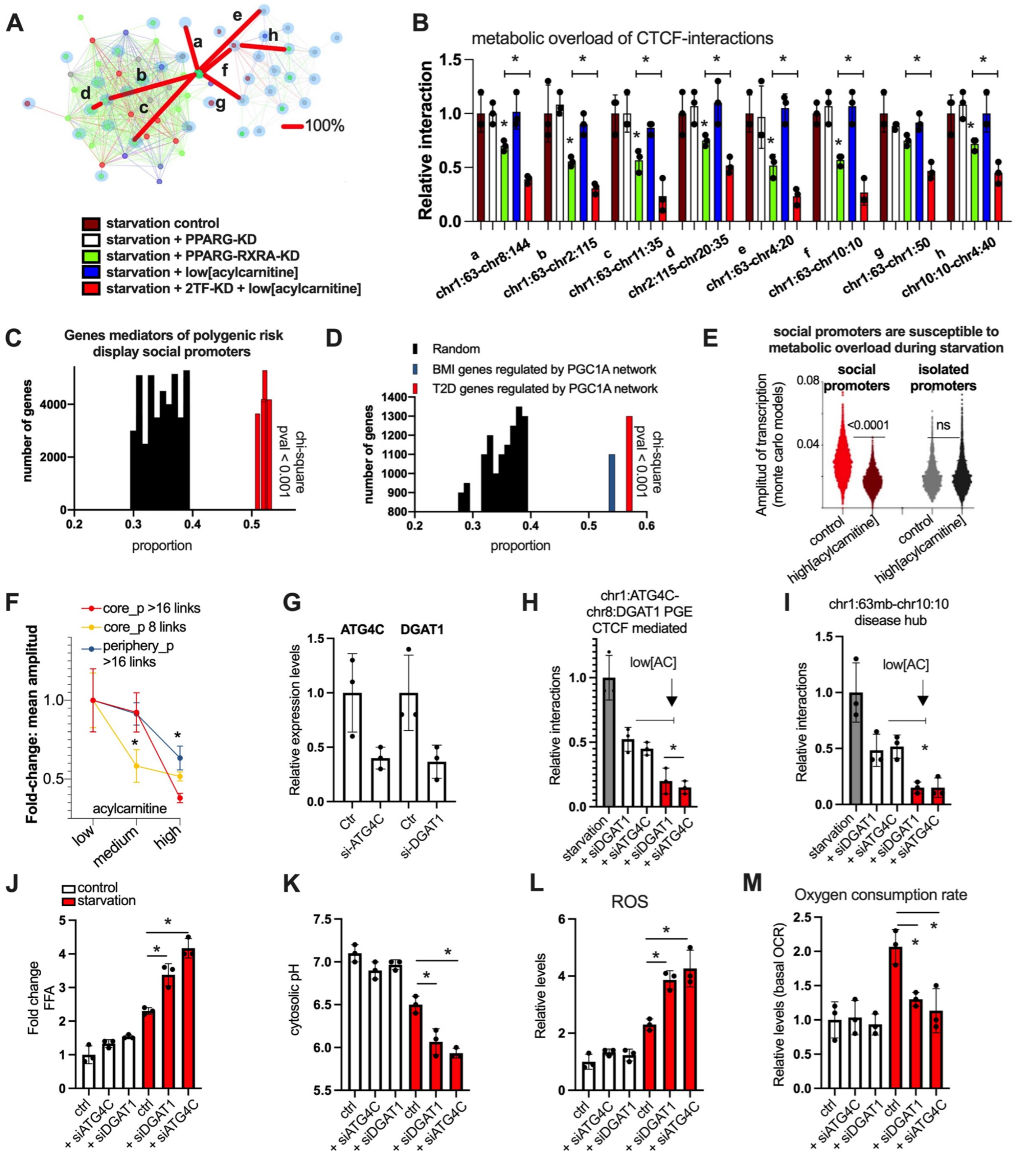


**Fig. S27. Combinatorial binding PGC1A-network to SIR-PRS network** *(related to Fig. 6)*. (**A**) BMI-T2D SIR- and BMI SIR-networks connected through top-bridging chromatin loci Chr1:60-65mb, harboring metabolic-HAR ATG4C region. Red lines showing starvation-mediated regulation of long-range interactions among diverse loci. (**B**) CTCF-mediated long-range interactions among loci displayed in A and with combinatorial transcriptional perturbation as well as metabolic overload wti low concentration of acylcarnitine (1uM). (**C**) Proportion of mediators of polygenic risk displaying social promoters as defined by their links in regulatory interaction networks in Fig 2. (**D**) Proportion of mediators of polygenic risk displaying social promoters as defined by their links in regulatory interaction networks in Fig 2, and being combinatorially bound by PGC1A-interacting regulators. (**E**) Amplitude for transcriptional burst models by Monte Carlo for genes within metabolic-HAr domains with either social and isolated promoters during starvation and with metabolic overload (acylcarnitine excess 20uM). (**F**) Mean amplitude of transcriptional models of core and peripheral genes with social promoters (>16 enhancer-promoter interactions) and mid-social promoters (8 interactions) in human adipocytes after starvation and treated with different concentrations of acylcarnitine (low=1uM; medium=10uM; high=20uM). (**G**) Gene expression levels after targeted perturbation with siRNA. (**H**) metabolic-HAR region interactions in starvation and after targeted perturbation of top regulated genes ATG4C and DGAT1, and with low acylcarnitine. (**I**) Long-range interactions for different disease loci hubs. (**J**) Functional assays in the same conditions as in H. Free-fatty acids (**J**), cytosolic pH (**K**), oxidative stress (**L**) and oxygen consumption rate by seahorse (**M**). Bars show mean values and error bars indicate SEM. Unpaired, two-tailed student's t-test was used when two groups were compared, and ANOVA followed by fisher's least significant difference (LSD) test for post hoc comparisons for multiple groups. * p-value <0.05.

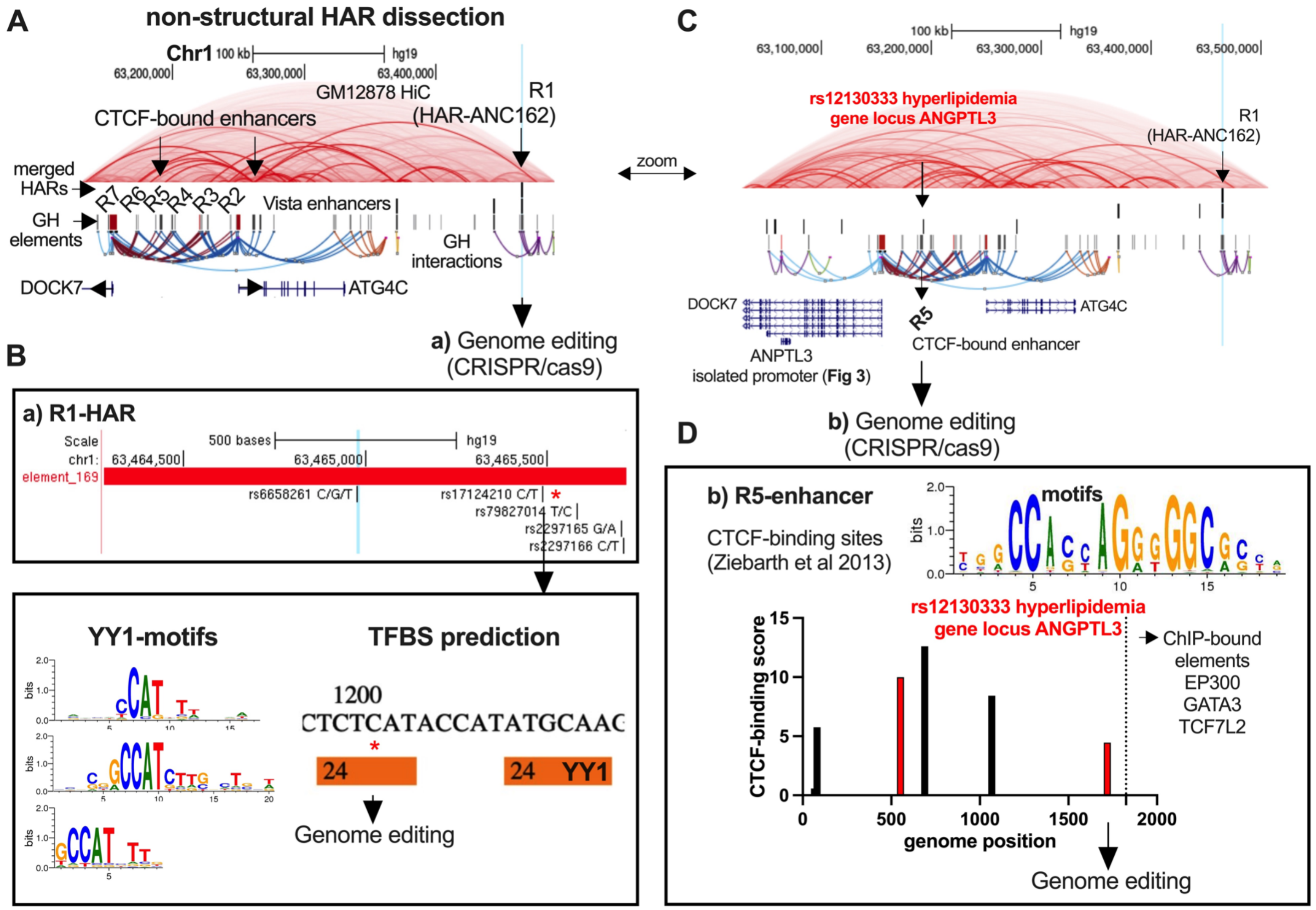


**Fig. S28. Disease circuit dissection and genome editing of metabolic-HAR ATG4C-domain** *(related to Fig. 6)*. (**A**) Human genome track showing top targets within metabolic-HAR domain in chr1:62-63mb: ncHARs (ANC162), regulatory elements and interactions, GM12878-Hi-C data, CTCF-bound elements, and promoter locations for ATG4C and DOCK7. (**B**) HAR-ANC162 (element_169) displaying variants within the element. With an arrow, location of TFBS for YY1 transcription factor, which was selected for genome editing. (**C**) Human genome track as in A, showing with arrows selected enhancers, in which R5 element (CTCF-bound) harbours a SNP related to hyperlipidemia (rs12130333). (**D**) R5 element was scanned for CTCF-binding sites as described by Ziebarth et al (*101*). A CTCF-binding site in close proximity to the variant rs12130333 was selected for genome editing.

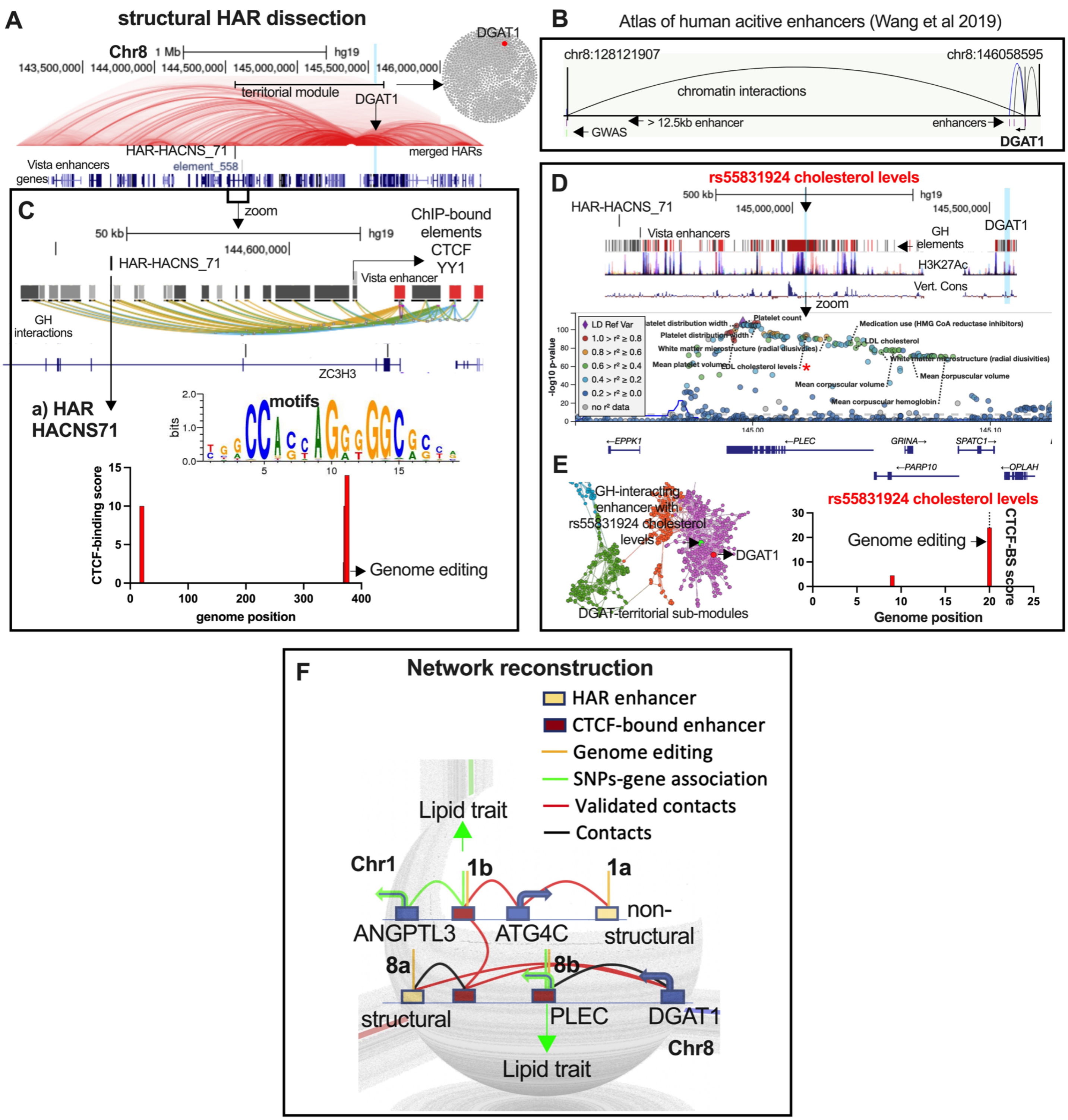


**Fig. S29. Disease circuit dissection and genome editing of metabolic-HAR DGAT1-domain** *(related to Fig. 6)*. (**A**) Human genome track showing top targets within metabolic-HAR domain in chr8:144-146mb: ncHARs (HACNS_71), vista regulatory element 558, GM12878-Hi-C data, DGAT1-promoter location, and DGAT1-module reconstruction from regulatory elements interaction network. (**B**) Enhancer long-range regulation of DGAT1 from the atlas of human active enhancers database (*147*). (**C**) HAR-HACNS_71, vista regulatory element 558, regulatory elements and interactions by GeneHancer. With an arrow to the lowest panel showing CTCF-binding sites in the HAR element, one of which was selected for genome editing. (**D**) Human genome track as in A, showing with an arrow selected variant related to cholesterol levels in regulatory element, followed in middle panel by all SNPs within region by HugAmp metabolic database and with an asterisk the lipid-related variant rs55831924. (**E**) Force atlas layout of DGAT1 module and submodules from genome-wide regulatory region interactions. Highlighted in green the lipid-variant and in red DGAT1 promoter element. In the right panel, CTCF-binding sites for the element harbouring the lipid-variant. (**F**) Full trans-network reconstruction Chr1-Chr8 coregulated metabolic HAR domains. In green arrow, lipid-related variants within structural enhancers, HAR-locations, target genes, chromatin contacts as well as notation for genome editing interventions.

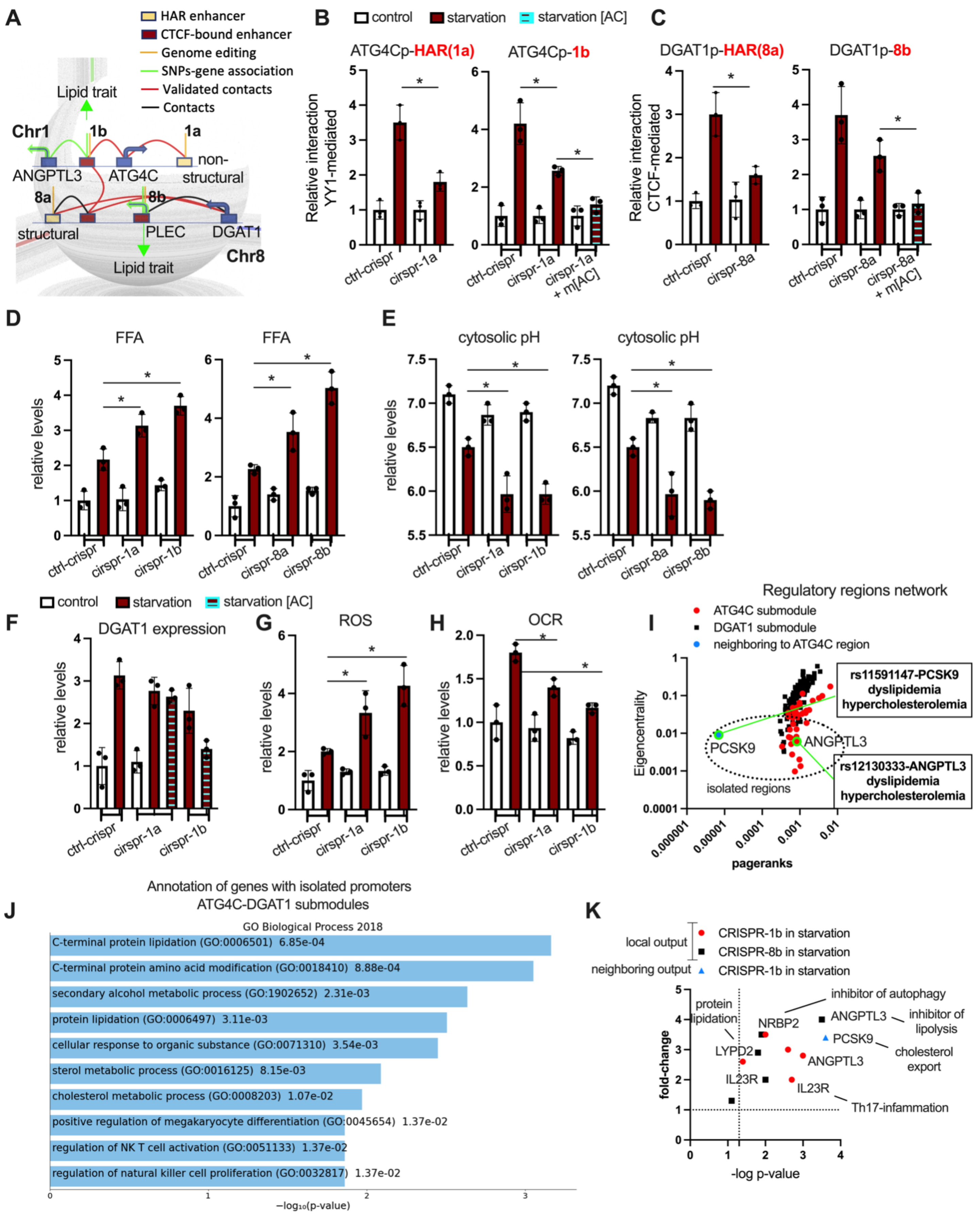

**Fig. S30. Disease circuit dissection and genome editing of trans-interacting metabolic-HAR domains** *(related to Fig. 6)***.** (**A**) Full trans-network reconstruction Chr1-Chr8 corregulated metabolic HAR domains. In green arrow, lipid-related variants within structural enhancers, HAR-locations, target genes, chromatin contacts as well as notation for genome editing interventions. (**B**) Left panel shows relative YY1-mediated loops of HAR-ATG4C_promoter in control mutation and perturbation of HAR YY1 binding site during starvation. Right panel shows enhancer 1b-ATG4C_promoter loops in the same conditions (control mutation and HAR-mutation) and with metabolic overload with acylcarnitine (medium concentration of 10uM). (**C**) Left panel shows relative CTCF-mediated loops of HAR-DGAT1_promoter in control mutation and perturbation of HAR CTCF-binding site during starvation. Right panel shows enhancer 8b-DGAT1_promoter loops in the same conditions (control mutation and HAR-mutation) and with metabolic overload with acylcarnitine (medium concentration of 10uM). (**D**) Free-fatty acids functional assays in the same conditions comparing control mutations, HAR mutations (a) and mutation in key-structural enhancers (b). (**E**) Cytosolic pH functional assays in the same conditions comparing control mutations, HAR mutations (a) and mutation in key-structural enhancers (b). (**F**) Trans-regulatory effects on chr8 DGAT1 expression from chr1-mutations in starvation and with acylcarnitine excess (10uM). (**G**) Oxidative stress and (**H**) oxygen consumption rate functional assays in the same conditions and in response to mutations in chr1 (a and b). (**I**) Node-prioritization scores for ATG4C, DGAT1 and ATG4C-neighboring regulatory submodules. Highlighted in dotted circle promoter regions with few connections and defined as isolated in Fig. 2, and in green those regions sharing strong genetic association to lipid-related traits. (**J**) Functional annotation of isolated promoters in both submodules. (**K**) Gene expression of top isolated genes in the same conditions as in B comparing control and key-structural mutations. Bars show mean values and error bars indicate SEM. Unpaired, two-tailed student's t-test was used when two groups were compared, and ANOVA followed by fisher's least significant difference (LSD) test for post hoc comparisons for multiple groups. * p-value <0.05.

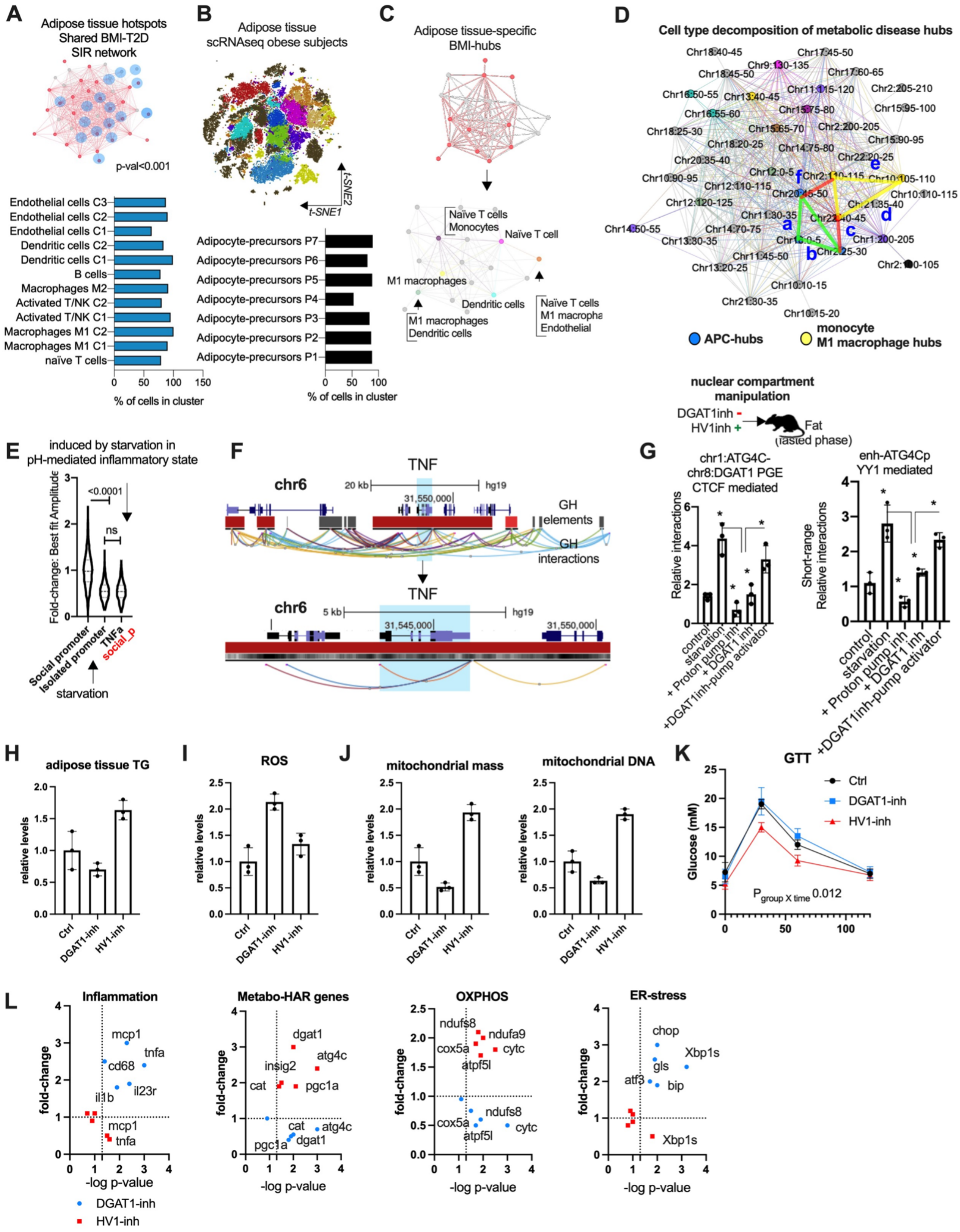
A
Adipose tissue hotspots
Shared BMI-T2D
SIR network
p-val<0.001
Endothelial cells C3
Endothelial cells C2
Endothelial cells C1
Dendritic cells C2
Dendritic cells C1
B cells
Macrophages M2
Activated T/NK C2
Activated T/NK C1
Macrophages M1 C2
Macrophages M1 C1
naïve T cells
% of cells in cluster
B
Adipose tissue
scRNAseq obese subjects
t-SNE1
t-SNE2
Adipocyte-precursors P7
Adipocyte-precursors P6
Adipocyte-precursors P5
Adipocyte-precursors P4
Adipocyte-precursors P3
Adipocyte-precursors P2
Adipocyte-precursors P1
% of cells in cluster
C
Adipose tissue-specific
BMI-hubs
Naïve T cells
Monocytes
Naïve T cell
M1 macrophages
Dendritic cells
M1 macrophages
Dendritic cells
Naïve T cells
M1 macropha
Endothelial
D
Cell type decomposition of metabolic disease hubs
APC-hubs
monocyte
M1 macrophage hubs
nuclear compartment
manipulation
DGAT1inh -
HV1inh +
Fat
(fasted phase)
E
induced by starvation in
pH-mediated inflammatory state
Fold-change: Best fit Amplitude
<0.0001
ns
Social promoter
Isolated promoter
TNFa
social_p
starvation
F
chr6
TNF
20 kb
hg19
31,550,000
GH
elements
GH
interactions
5 kb
31,545,000
G
chr1:ATG4C-
chr8:DGAT1 PGE
CTCF mediated
Relative interactions
enh-ATG4Cp
YY1 mediated
Short-range
Relative interactions
control
starvation
+ Proton pump inh
+ DGAT1 inh
+DGAT1inh-pump activator
H
adipose tissue TG
relative levels
Ctrl
DGAT1-inh
HV1-inh
I
ROS
J
mitochondrial mass
mitochondrial DNA
K
GTT
Glucose (mM)
Ctrl
DGAT1-inh
HV1-inh
P group X time 0.012
L
Inflammation
Metabo-HAR genes
OXPHOS
ER-stress
fold-change
-log p-value
mcp1
tnfa
cd68
il23r
il1b
dgat1
atg4c
insig2
cat
pgc1a
ndufs8
ndufa9
cox5a
cytc
atpf5l
chop
Xbp1s
gls
atf3
bip
DGAT1-inh
HV1-inh

**Fig. S31. Single cell decomposition of metabolic disease hubs in human fat tissue and mammalian in-vivo manipulation of nuclear compartment formation** *(related to Fig. 6)*. (**A**) Social interchromosomal region (SIR) network of trans-interacting chromatin regions and associated metabolic-HAR domains (blue nodes). In red, nodes harbouring fat-specific transcriptional hubs from mediation of polygenic risk for T2D and BMI in fat tissue. Overall modularity enrichment by z-score and p-value <0.05. (**B**) Cluster specific t-SNE plots from human adipose tissue scRNA seq and percentage of cell types within. (**C**) SIR-network and cell-specific decomposition of transcriptional hotspots found in differential expression scRNA seq by positional enrichments. (**D**) Full SIR-network coordinates with different cell-specific hubs. Highlighted in blue and connected by green lines adipose precursor cell hubs. In yellow lines monocyte and M1 macrophages cells specific hubs. (**E**) Mean amplitude from Monte Carlo models of dynamic expression profiles in adipocytes during starvation. Comparing mean amplitude for social promoter gene DOCK7, isolated promoter gene ANGPTL3, and TNF burst kinetics. (**F**) Human genome track picture showing chr6 region harboring TNF gene, enhancer elements and interactions. In the lower panel, TNF gene span with local interactions. (**G**) Top panel shows illustration for acute in vivo intervention by intraperitoneal injections of fasting mice with either DGAT1-inhibitor or HV1-inhibitor (2mg/kg). In lower panels, CTCF-long range and YY1 short-range relative interactions in the same conditions. (**H**) Adipose tissue functional assays after chronic manipulation of nuclear compartment formation in mice treated daily with either inhibitor as in G during 7 days: (**H**) triglycerides, (**I**) oxidative stress, (**J**) mitochondrial mass and mitochondrial DNA. (**K**) Glucose tolerance test in the same conditions as in H. (**L**) Gene expression profile in adipose tissue in the same conditions as in H. Bars show mean values and error bars indicate SEM. Unpaired, two-tailed student's t-test was used when two groups were compared, and ANOVA followed by fisher's least significant difference (LSD) test for post hoc comparisons for multiple groups. * p-value <0.05.

#### Supplementary Tables

**Table S1. Metabolic phenotype comparison across mammals**

A. Metabolic parameters and fasting endurance score in nonprimate mammals
B. Metabolic parameters and fasting endurance score in primate mammals
C. Metabolic parameters and fasting endurance score in hominoids
D. Tissue mass and metabolic parameters across mammals

Link: https://www.dropbox.com/s/7czlr2dkukutsxl/Table%20S1.xlsx?dl=0

**Table S2. Differential human specific molecular signatures in fat and brain**

A. GREAT analysis from ATACseq comparison between human-primate in fat: human increase
B. GREAT analysis from ATACseq comparison between human-primate in fat: human decrease
C. Fat branch-specific acceleration and positive selection from ATACseq comparison between humans and primates
D. Accelerated evolution signature genes in brain

Link: https://www.dropbox.com/s/h0bva3hjg0jx37c/Table%20S2.xlsx?dl=0

**Table S3. Human-accelerated regulatory regions**

A. List of human accelerated regions coordinates
B. HAR-associated genes discovery from Capra et al. 2013
C. Full list of HAR-genes compiled for this study
D. Functional annotation of HAR genes in brain-specific networks
E. Functional annotation of HAR genes in fat-specific networks

Link: https://www.dropbox.com/s/j89lhmskpxoo8x7/Table%20S3.xlsx?dl=0

**Table S4. Transcriptional regulatory networks of HAR-genes**

A. ChIP-seq enrichment ChIP-atlas of HAR-genes
B. Adipocytes ChIP-seq enrichment ChIP-atlas of HAR-genes
C. Neural ChIP-seq enrichment ChIP-atlas of HAR-genes
D. Stem cells ChIP-seq enrichment ChIP-atlas of HAR-genes
E. Blood ChIP-seq enrichment ChIP-atlas of HAR-genes
F. Muscle ChIP-seq enrichment ChIP-atlas of HAR-genes
G. Liver ChIP-seq enrichment ChIP-atlas of HAR-genes
H. TFBS enrichment from MSDB
I. TFBS enrichment form genome browser
J. ENCODE and ChEA consensus enrichment

https://www.dropbox.com/s/1sgdr5741pqg6t0/Table%20S4.xlsx?dl=0

**Table S5. Transcriptional regulatory networks, and PPI reconstitution**

A. Node + 1 network reconstruction of PPIs for HAR-genes regulators
B. Network topological prioritization of network in A
C. Full PGC1A-network reconstruction of protein-protein interactions
D. Network topological prioritization of full PGC1A-network
E. Reactome functional annotation of HAR cooperative regulators
F. Gene-ontology functional annotation of HAR cooperative regulators
G. Fat and brain tissue-specific PPIs for PGC1A-network

https://www.dropbox.com/s/brp2olmem6o5zgy/Table%20S5.xlsx?dl=0

**Table S6. Intrinsically disordered regions of HAR-gene regulators**

A. Consensus disordered regions in PGC1A protein
B. Hydropathy boundary ordered and disordered proteins
C. Multivalent pi-pi disordered score human proteins
D. DisProt disordered score human proteins

https://www.dropbox.com/s/idqxtlylhl1lg24/Table%20S6.xlsx?dl=0

**Table S7. Phylogenetic conservation of co-activator protein residues**

A. Consurf amino acid variation of PGC1A residues
B. Amino acid conservation scores for PGC1A co-activator

https://www.dropbox.com/s/57k3ql68793bwql/Table%20S7.xlsx?dl=0

**Table S8. Combinatorial transcriptional regulation of HAR-genes**

A. Functional interacting transcriptional regulators co-regulated genes by promoter ChIP-seq enrichment
B. Combinatorial co-regulated HAR-genes (metabolic-HAR genes)

https://www.dropbox.com/s/0zq6kakrtw5q773/Table%20S8.xlsx?dl=0

**Table S9. Functional topological loci decomposition of metabolic-HAR regions**

A. Positional gene enrichments of metabolic-HAR genes
B. Cytogenetic band enrichments of metabolic-HAR genes
C. Functional annotation of metabolic-HAR PGE-domains

https://www.dropbox.com/s/10wlpjohnu4n83u/Table%20S9.xlsx?dl=0

**Table S10. Compendia of HiC data**

A. HiC chromatin interacting regions
B. Information on HiC compendia from Schmitt et al. 2016
C. Frequently interacting regions FIREs
D. Conserved TADs

https://www.dropbox.com/s/uejs9r1h405i5n2/Table%20S10.xlsx?dl=0

**Table S11. hESC interacting interchromosomal regions**

A. Network from hESC-HiC interacting chromatin segments and metabolic-HAR regions overlap
B. Network ID of interacting segments in A
C. Topological prioritization and network analysis of hESC-HiC interacting chromatin segments

https://www.dropbox.com/s/4hl6jvb7l87enbb/Table%20S11.xlsx?dl=0

**Table S12. Genome-wide short-range enhancer-promoter relationships**

A. GeneHancer database of enhancer-promoter associations.

https://www.dropbox.com/s/ih9mof2smzyc6os/Table%20S12.xlsx?dl=0

**Table S13. Genome-wide short-range regulatory elements network**

A. Regulatory elements associations
B. Network analyses of regulatory elements associations
C. Module-based and degree prioritization
D. Degree-based prioritization of enhancers
E. Social promoter genes and genomic cluster positions
F. Nonsocial promoter genes and genomic cluster positions

https://www.dropbox.com/s/4b4u8hmns6u3el9/Table%20S13.xlsx?dl=0

**Table S14. metabolic-HAR domains short-range regulatory subnetwork from reconstituted nuclear compartment**

A. ATG4C-submodule short-range relationships
B. DGAT1-submodule short-range relationships
C. PGC1A-submodule short-range relationships
D. CAT-submodule short-range relationships
E. ATG4C-submodule territorial ChIP-seq enrichment for social enhancer and promoters
F. DGAT-submodule territorial ChIP-seq enrichment for social enhancer and promoters
G. PGC1A-submodule territorial ChIP-seq enrichment for social enhancer and promoters
H. CAT-submodule short-range relationships
I. PPI network reconstruction of territorial ChIP-seq enrichment of regulators within ATG4C submodule

https://www.dropbox.com/s/dcy05kn2hjbcaz2/Table%20S14.xlsx?dl=0

**Table S15. Active enhancer elements**

A. Vista active enhancer elements human and mouse coordinates and associated genes
B. Enhancer identity and human coordinates

https://www.dropbox.com/s/2tpw0kos0xqyndv/Table%20S15.xlsx?dl=0

**Table S16. Structural variants, genomic range function and genetics of metabolic-HAR compartments**

A. Common structural variants with SNVs
B. Number of SVs in reconstituted nuclear compartment with metabolic-HAR domains
C. ATG4C metabolic-HAR domain genomic range genetics (chr1:62mb)
D. INSIG2 metabolic-HAR domain genomic range genetics (chr2:110mb)
E. PGC1A metabolic-HAR domain genomic range genetics (chr4:25mb)
F. CAT metabolic-HAR domain genomic range genetics (chr8:38mb)
G. APOA5 metabolic-HAR domain genomic range genetics (chr11:117mb)
H. GREAT analysis metabolic-HAR genomic coordinates

https://www.dropbox.com/s/gun54eshm93xl3u/Table%20S16.xlsx?dl=0

**Table S17. Epimap network modeling of metabolic-HAR regions**

A. Epimap all SNPs-enhancer intersections
B. Epimap all GWAS-SNPs associations
C. Tissue and trait network topological dependencies
D. Chromatin loci network topological dependencies
E. Transcriptional hubs from Epimap-modular dependencies

https://www.dropbox.com/s/jth6erg3syhkf4c/Table%20S17.xlsx?dl=0

**Table S18. ChIP-MS structuring candidates and PGC1A-network**

A. List of proteins identified by chromatin proteomic profiling in mESC dual enhancer-promoter
B. Interrogation of PPIs for PGC1A-network and ChIP-MS structuring candidates
C. IDs node summary annotation for B
D. Functional network annotation
E. Network topological dependencies

https://www.dropbox.com/s/c8helx3ck1engfa/Table%20S18.xlsx?dl=0

**Table S19. Tissue- and cell-specific enhancer activity**

A. Tissue- and cell-specific enhancer activity
B. Global mean and sum of activity for each enhancer coordinate
C. Adipocyte-specific enhancer

https://www.dropbox.com/s/10ojb9escn0h9zm/Table%20S19.xlsx?dl=0

**Table S20. Human active enhancers from GRO-seq, PRO-seq and CAGE data**

https://www.dropbox.com/s/whsch6czar35mrz/Table%20S20.txt?dl=0

**Table S21. Human adipocyte precursors profiling and brain cortex snRNA**

A. Cell-specific genes in snRNAseq from human cortex
B. Human adipocyte precursors cell profiling

https://www.dropbox.com/s/dnw1anv8jqznk9e/Table%20S21.xlsx?dl=0

**Table S22. Synteny of metabolic-HAR regions and mammalian perturbation networks**

A. Murine transcriptional hubs of metabolic-HAR regions
B. Metabolic-HAR gene perturbation-phenotype association
C. Gene network topological dependencies
D. Phenotype network topological dependencies

https://www.dropbox.com/s/7aq1nhhnwsbp5fq/Table%20S22.xlsx?dl=0

**Table S23. Differentially expressed genes in diet-induced obesity across 19 organs**

A. Mesenteric adipose tissue. B. Brown adipose tissue. C. Visceral adipose tissue. D. Retroperitoneal adipose tissue. E. Subcutaneous adipose tissue. F. Hypothalamus. G. Ileum. H. Jejunum. I. Kidney. J. Liver. K. Stomach. L. Duodenum. M. Soleus. N. Colon. O. Cortex. P. Adrenal gland. Q. Cerebellum. R. Midbrain stem.

https://www.dropbox.com/s/lmd3k8k8xaiveth/Table%20S23.xlsx?dl=0

**Table S25. Transcriptional hubs controlled by diet**

A. Mesenteric adipose tissue. B. Brown adipose tissue. C. Visceral adipose tissue. D. Retroperitoneal adipose

tissue. E. Subcutaneous adipose tissue. F. Hypothalamus. G. Ileum. H. Jejunum. I. Kidney. J. Liver. K. Stomach. L. Duodenum. M. Soleus. N. Colon. O. Cortex. P. Adrenal gland. Q. Cerebellum. R. Midbrain stem.
https://www.dropbox.com/s/8e2w8wrzxtvxsws/Table%20S24.xlsx?dl=0

**Table S25. Fat-specific transcriptional hubs**

A. Fat-specific transcriptional hubs controlled by diet
https://www.dropbox.com/s/bq1fe09wox9krkr/Table%20S25.xlsx?dl=0

**Table S26. Single cell topological decomposition of fat transcriptional hubs**

A. Cell-specific genes in murine adipose from Tabula Muris consortium
B. Natural killer cell-specific transcriptional hubs in murine adipose tissue
C. T cell-specific transcriptional hubs in murine adipose tissue
D. Myeloid cell-specific transcriptional hubs in murine adipose tissue
E. B cell-specific transcriptional hubs in murine adipose tissue
F. Mesenchymal cell-specific transcriptional hubs in murine adipose tissue
G. Endothelial cell-specific transcriptional hubs in murine adipose tissue
https://www.dropbox.com/s/oesjvj972mu0dna/Table%20S26.xlsx?dl=0

**Table S27. Multi-tissue mediators of polygenic risk scores for BMI and T2D**

A. Tissue-specific mediators of polygenic risk for T2D (BMI-adjusted)
B. Tissue-specific mediators of polygenic risk for BMI (GIANT)
C. Genomic hubs for BMI mediators of polygenic risk
D. Genomic hubs for T2D mediators of polygenic risk
https://www.dropbox.com/s/efr9d3oq1ihikh4/Table%20S27.xlsx?dl=0

**Table S28. Radial chromatin organization**

A. GPseq radial chromatin loci coordinates
B. DNA-MERFISH chromatin organization
C. Ontology of core and nuclear speckles associated genes with transcriptional burst on from B
D. Biological enrichment of core and nuclear speckles associated genes with transcriptional burst on
https://www.dropbox.com/s/n6iayzilav1x8if/Table%20S28.xlsx?dl=0

**Table S29. Radial transcriptional regulators**

A. TFBS density and GPseq score radial correlations
B. Functional interacting regulators with peripheral enrichment
C. Node + 1 network reconstruction of B
D. Functional interacting regulators with core enrichment
E. Node + 1 network reconstruction of D
https://www.dropbox.com/s/ymx5kshsev9kpn1/Table%20S29.xlsx?dl=0

**Table S30. Radial transcriptional condensate features**

A. Amino acid sequences for core enriched regulators
B. Amino acid sequences for peripherally enriched regulators
C. Context-dependent condensate features for core enriched regulators
D. Context-dependent condensate features for peripherally enriched regulators
https://www.dropbox.com/s/v69f5yb6p7ap73y/Table%20S30.xlsx?dl=0

**Table S31. Social interchromosomal network: interacting metabolic disease hubs and metabolic-HAR regions**

A. Social interchromosomal regions: 5mb range interacting scores
B. Interacting metabolic disease hubs for shared hotspots for BMI and T2D polygenic risk
C. Interacting metabolic disease hubs coordinates for BMI mediators of polygenic risk
D. Interacting metabolic disease hubs coordinates for T2D mediators of polygenic risk
E. Mammalian phenotype integration of gene perturbations for shared mediators of BMI and T2D polygenic risk
F. Network topological dependencies for mediators
G. Network topological dependencies for phenotypes
https://www.dropbox.com/s/47fnm8cqlijs5ki/Table%20S31.xlsx?dl=0

**Table S32. Interacting metabolic disease hubs regulated by PGC1A-network**

A. Combinatorial regulation of T2D hotspots by PGC1A-network
B. Combinatorial regulation of BMI hotspots by PGC1A-network
C. Gene list > 4 combinatorial PGC1A-network for T2D-BMI mediators of polygenic risk
D. Genomic clusters by cytogenetic band enrichments
E. Genomic clusters by positional gene enrichments

https://www.dropbox.com/s/0b95i0r8x3ng4n4/Table%20S32.xlsx?dl=0

**Table S33. Single cell topological decomposition of interacting metabolic disease hubs**

A. Subcutaneous progenitors genes in obese fat scRNAseq
B. Visceral progenitors genes in obese fat scRNA
C. Immune cell-specific genes in obese fat scRNAseq
D. Endothelial cell-specific genes in obese fat scRNAseq
E. Genomic hubs from fat progenitors scRNAseq genes
F. Genomic hubs from fat immune scRNAseq genes

https://www.dropbox.com/s/ml1zqsonmgezt6h/Table%20S33.xlsx?dl=0

179. A. Mahajan, D. Taliun, M. Thurner, N. R. Robertson, J. M. Torres, N. W. Rayner, A. J. Payne, V. Steinthorsdottir, R. A. Scott, N. Grarup, J. P. Cook, E. M. Schmidt, M. Wuttke, C. Sarnowski, R. Mägi, J. Nano, C. Gieger, S. Trompet, C. Lecoeur, M. H. Preuss, B. P. Prins, X. Guo, L. F. Bielak, J. E. Below, D. W. Bowden, J. C. Chambers, Y. J. Kim, M. C. Y. Ng, L. E. Petty, X. Sim, W. Zhang, A. J. Bennett, J. Bork-Jensen, C. M. Brummett, M. Canouil, K.-U. Ec Kardt, K. Fischer, S. L. R. Kardia, F. Kronenberg, K. Läll, C.-T. Liu, A. E. Locke, J. 'an Luan, I. Ntalla, V. Nylander, S. Schönherr, C. Schurmann, L. Yengo, E. P. Bottinger, I. Brandslund, C. Christensen, G. Dedoussis, J. C. Florez, I. Ford, O. H. Franco, T. M. Frayling, V. Giedraitis, S. Hackinger, A. T. Hattersley, C. Herder, M. A. Ikram, M. Ingelsson, M. E. Jørgensen, T. Jørgensen, J. Kriebel, J. Kuusisto, S. Ligthart, C. M. Lindgren, A. Linneberg, V. Lyssenko, V. Mamakou, T. Meitinger, K. L. Mohlke, A. D. Morris, G. Nadkarni, J. S. Pankow, A. Peters, N. Sattar, A. Stančáková, K. Strauch, K. D. Taylor, B. Thorand, G. Thorleifsson, U. Thorsteinsdottir, J. Tuomilehto, D. R. Witte, J. Dupuis, P. A. Peyser, E. Zeggini, R. J. F. Loos, P. Froguel, E. Ingelsson, L. Lind, L. Groop, M. Laakso, F. S. Collins, J. W. Jukema, C. N. A. Palmer, H. Grallert, A. Metspalu, A. Dehghan, A. Köttgen, G. R. Abecasis, J. B. Meigs, J. I. Rotter, J. Marchini, O. Pedersen, T. Hansen, C. Langenberg, N. J. Wareham, K. Stefansson, A. L. Gloyn, A. P. Morris, M.

Boehnke, M. I. McCarthy, Fine-mapping type 2 diabetes loci to single-variant resolution using high-density imputation and islet-specific epigenome maps. *Nat. Genet.* **50**, 1505–1513 (2018).
